# A progesterone derivative linked to a stable phospholipid activates breast cancer cell response without leaving the cell membrane

**DOI:** 10.1101/2023.05.14.540708

**Authors:** Jofre Font-Mateu, Pol Sanllehí, Jesús Sot, Beatriz Abad, Nicolas Mateos, Juan Andres Torreno-Pina, Roberto Ferrari, Roni H.G. Wright, Maria Garcia-Parajo, Jesús Joglar, Félix M. Goñi, Miguel Beato

## Abstract

In hormone-responsive breast cancer cells, progesterone (P4) has been shown to act via its nuclear receptor (PR), a ligand-activated transcription factor. A small fraction of PR is palmitoylated and anchored to the cell membrane (mbPR) forming a complex with estrogen receptor alpha (ERα). Upon hormone exposure, either directly or via interaction with ERα mbPR activates the SRC/RAS/ERK kinase pathway leading to phosphorylation of PR by ERK. Kinase activation is essential for P4 gene regulation, as the ERK and MSK1 kinases are recruited by the PR to its genomic binding sites and trigger chromatin remodeling. An interesting open question is whether activation of mbPR can result in gene regulation in the absence of ligand binding to intracellular PR. This matter has been investigated in the past using P4 attached to serum albumin, but the attachment is leaky and albumin can be endocytosed and degraded, liberating P4. Here we propose a more stringent approach to address this issue by ensuring attachment of P4 to the cell membrane via covalent binding to a stable phospholipid. This strategy identifies the actions of P4 independent of hormone binding to intracellular PR. We found that a membrane-attached progestin can activate mbPR, the ERK signalling pathway leading to PR phosphorylation, initial gene regulation and entry into the cell cycle, in the absence of detectable intracellular progestin.

## INTRODUCTION

The dominant view of steroid hormone action is that these amphipathic molecules diffuse through the cell membrane and interact with their intracellular receptors, members of the large nuclear receptor family, which shuttle between the cytoplasm and cell nucleus.^1^ In response to binding their specific hormone ligand, the intracellular receptors dissociate from chaperones, dimerize and concentrate in the cell nucleus, where they interact with DNA sequences called hormone response elements (HRE), or with other transcription factors, and regulate the rate of transcription of their target genes.^2^ Progesterone (P4) is a steroid hormone that regulates target cell behavior in many tissues via interaction with the intracellular progesterone receptor (PR). In breast cancer cells, we have described the interaction of ligand-activated PR with progesterone responsive elements (PREs) in chromatin and the subsequent chromatin remodeling steps required for gene regulation (for a recent review see Beato *et al* 2020).^3^ However, in addition to these well-established genomic pathways, steroid hormones are known to act via a non-genomic mechanism.^4^ In breast cancer cells estradiol activates the SRC/RAS/ERK kinase pathway via a small fraction of the nuclear estrogen receptor alpha (ERα) attached to the cell membrane (mbERα).^5^ We found that in T47D breast cancer cells this kinase pathway is also activated by progestins via a small fraction of PR attached to the cell membrane (mbPR) in a complex with mbERα.^6^ Upon P4 exposure, the SRC/RAS/ERK kinase pathway is activated.^7, 8^ It is also known that PR membrane attachment is mediated by the palmitoylation of cysteine 820, located in a short amino acid region of the ligand binding domain conserved in other steroid hormone receptors.^9^ After exposure to 10 nM progestin, the mbPR remains attached to the cell membrane, while the activated ERK kinase phosphorylates intracellular PR at serine 294, leading to its activation, displacement of its associated chaperones and dimerization. This activated phosphoPR (pPR) forms a ternary complex with activated ERK and MSK1 that binds to PREs in the nucleus and initiates the changes in chromatin and transcription factor recruitment that lead to the regulation of P4-responsive genes.^7, 10^ In recent studies we found that progestins are able to activate the response of breast cancer cells at 50 pM concentration, 200-fold lower than used in most previous studies, by activating the kinase signalling pathway and promoting pPR binding to a small subset of accessible enhancers.^11^

An interesting question is whether hormone entry into the cells and binding to the intracellular PR is necessary for activating cell response to P4. It is in principle possible that hormone binding to the mbPR and subsequent kinase activation is sufficient to promote phosphorylation of the ligand-free PR, followed by dissociation of pPR from the chaperones, dimerization, binding to accessible chromatin PREs, and activation of the cell response. In the past, several groups have used steroid hormones attached to serum albumin in an attempt to demonstrate that the hormones can act without crossing the cell membrane.^12, 13^ However, this strategy is not convincing for two main reasons: the attachment of steroids to albumin is non-covalent, and albumin can be endocytosed and degraded by lysosomes leading to release of P4 into the cytosol. Here we report an original and more stringent approach to ensure that the hormone remains firmly attached to the cell membrane during the experimental procedure, allowing the study of those hormonal functions, if any, that only require ligand binding to the mbPR. To this end, we attached a P4 derivative via a linker chain to the polar head group of a phosphatidylcholine analogue, an *Archaea* like phospholipid with two fatty acids linked to glycerol by ether bonds that cannot be cleaved by cellular phospholipases A2.^14^ Therefore, the steroid would be stably anchored to the membrane via the alkyl phospholipid, and the linker chain (4-8 ethylene oxide units) would provide space for the anchored steroid to reach the mbPR binding site.^15^ In addition, the lipid was fluorescently labeled in one of the alkyl chains to allow detection within the cells.^16^ The results show that our approach works in principle, as the membrane-attached P4 derivative activates the cell response to hormone, without entering the cell. This strategy may help in the study of the proteins associated with membrane-bound hormone receptors and their interactions with protein kinases. In addition, this study underscores the physiological importance of low levels of hormone receptors attached to the cell membrane in breast cancer, which opens the possibility to target them for blocking kinase activation as part of the endocrine therapy of cancer.

## RESULTS

### Design and synthesis of the probes

The probes used in this work (CRG033, CRG047, CRG034 and CRG048) consist of a progesterone derivative attached to the polar head group of a chemically modified lipid by means of a polyethylene glycol (PEG) linker of variable length (Figure 1).

**Figure 1.**
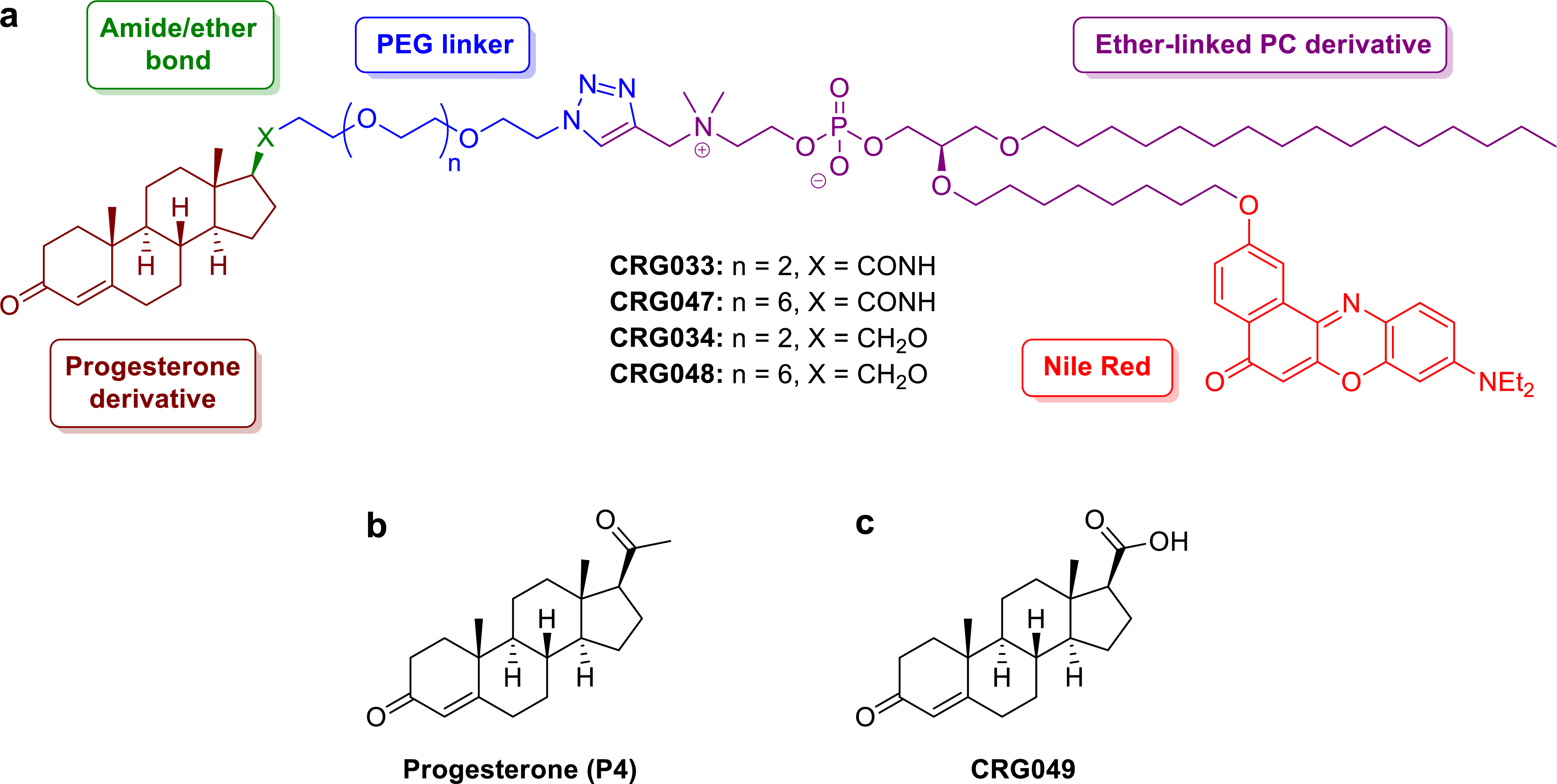
Chemical structure of synthesized compounds. a) Structure of the designed mPR probes CRG033, CRG047, CRG034 and CRG048. b) Structure of progesterone P4. c) Structure of compound CRG049, the hydrolysis byproduct of probes CRG033 and CRG047.

For a full account of the synthetic experimental part see the Supporting Information. Briefly, a Cu(I)-catalyzed azide-alkyne [3 + 2] dipolar cycloaddition (CuAAC) between an azide-terminated moiety bearing the progesterone derivative and an alkyne-containing lipid was envisaged as a suitable and efficient way to synthetically ensemble the targeted probes.^17, 18^ To provide a stable anchor to the cell membrane, an ether-linked phosphatidiylcholine derivative was used as the lipidic moiety of the probes. As opposed to ester-linked lipids, in this *Archaea*-like phospholipid structure the two acyl chains were linked to the glycerol by ether bonds, which conferred an increased metabolic stability against cellular phospholipases A2.^14^

In addition, the lipid was fluorescently labeled in one of the alkyl chains in order to allow probe detection during *in vitro* and *in vivo* experiments.^16^ Nile Red was chosen as a fluorophore. Under physiological conditions, Nile Red is an uncharged, low-polar molecule, which facilitates dye insertion into the cell membrane. Hence, a 2-hydroxy Nile Red derivative was placed at the distal position of one of the lipid alkyl chains via an ether bond (Scheme S4).^19–21^

As previously mentioned, a PEG linker was connecting the steroidal and lipid moieties of the probes. Insertion of a short-chain PEG conferred water solubility to the probes and increased their amphipathic nature^15^. In this case, we reasoned that linkers with either four (CRG033 and CRG034) or eight (CRG047 and CRG048) ethylene oxide units would provide enough space for the progesterone derivative to reach the mbPR binding site and exert its activity (Scheme S1).

Since the C3 ketone functionality is known to be crucial for PR activation,^22^ the progesterone fragment was attached to the probe at the 17β position of the steroidal scaffold. In compounds CRG033 and CRG047, the steroid was anchored to the linker *via* an amide bond, which preserved the carbonyl functionality of the natural ligand but could be susceptible of metabolic degradation by amidases. Thus, despite not bearing the C20 carbonyl group of progesterone (Fig. 1), the chemically and metabolically more stable ether-containing probes CRG034 and CRG048 were also prepared (Schemes S2, S3 and S4).

### Affinity of the synthesized compounds for PR

The chemical structure of several of the synthesized compounds and of a possible product of the hydrolysis of the amide bond of CRG033 or CRG047 (labelled CRG049) are shown in Figure 1. After their chemical and fluorometric characterization (see Supporting Information, Figure S1 and Table S1), we first explored their affinity for purified recombinant full length PR expressed in baculovirus^23^ using microscale thermophoresis (MST, see Experimental Section for details). Among tested compounds, only CRG047 exhibited a K_d_ for PR (≈ 14 nM) at the same low nM range as P4^24^ (Fig. 2a, b and Fig. S2 a-d). A comparable affinity was found using the recombinant ligand binding domain of PR expressed in *E. coli*^25^ (Fig. 2c). We therefore chose CRG047 for subsequent characterization.

**Figure 2.**
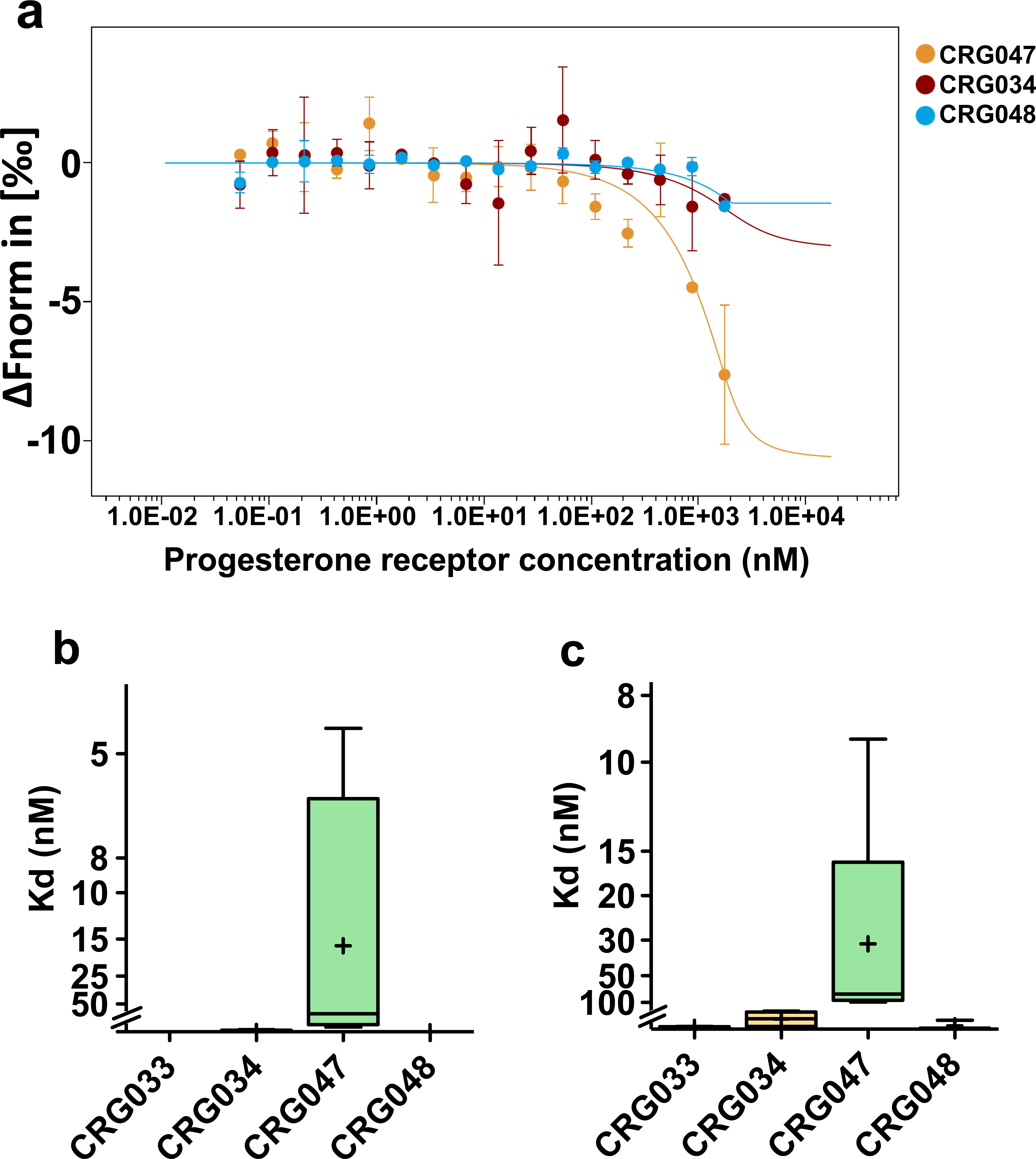
PR affinity with progesterone-like compounds. a) Microscale thermophoresis (MST). The thermophoresis changes as normalized fluorescence (ΔFnorm in [‰]) is plotted against the concentration of the unlabeled protein and fitted according to the law of mass action. Average ± SD of 2 readings from the same experiment is shown. b-c) Microscale thermophoresis (MST) analysis of 2 μM progesterone-like compounds with increasing concentrations of full-length progesterone receptor expressed in baculovirus (b) or GST-progesterone receptor Ligand-binding Domain (LBD) (c). Internal fluorescence of the Nile-red moiety in the compounds was used for the detection of the binding affinities. Due to precipitation, signal saturation was not reached. Boxplots show results from at least 4 independent experiments. ‘+’ indicates average (CRG047 Kd for full PR, 16.1 nM; for PR-LBD, 31.2 nM).

### CRG047 in lipid bilayers

The fluorescence properties of CRG047 in organic solvent (ethanol) are summarized in Figure 3a. For measuring the critical micellar concentration (CMC) of CRG047, increasing concentrations of the compound dissolved in DMSO were added to 25 mM HEPES, 150 mM NaCl, pH 7.4, and the increase in emitted fluorescence was measured to estimate its CMC, that was of ≈ 12 μM (Fig. 3b).

**Figure 3.**
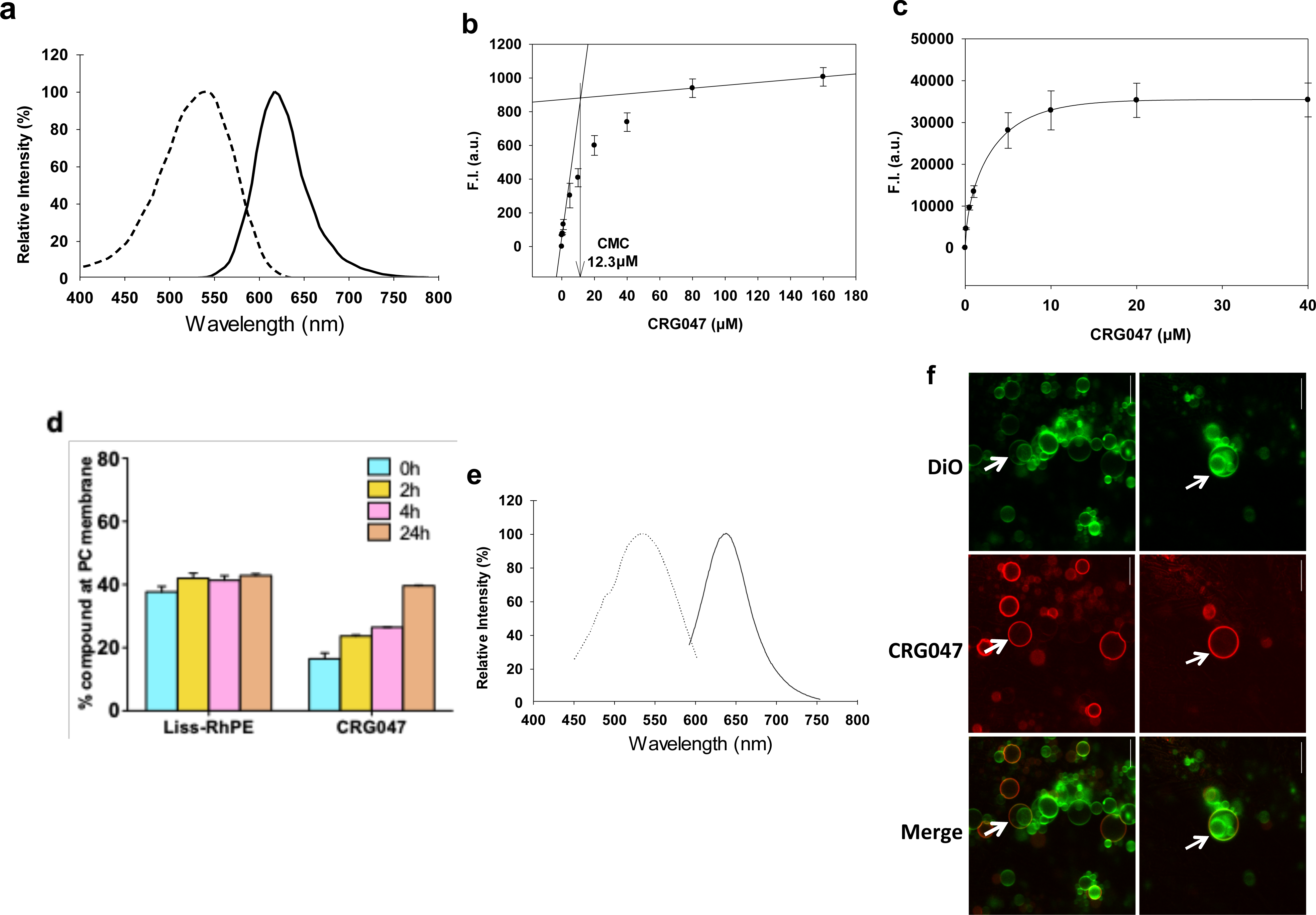
Biophysical properties of the CRG047 probe. a) Excitation and emission spectra of CRG047 in ethanol. b) Critical micellar concentration (CMC) of CRG047, measured by fluorescence spectroscopy. c) Dose-dependent fluorescence emission of CRG047 in small unilamellar vesicles (SUV) composed of egg PC. d) CRG047 incorporation into egg PC bilayers (SUV) as a function of time, and a comparison with the incorporation of Liss-RhPE as indicated. e) Excitation and emission spectra of CRG047 in SUV composed of egg PC. f) Incorporation of CRG047 into vesicles found in mostly unilamellar GUV composed of egg PC. Two representative fields are shown. Top: Vesicles stained with DiO during the formation of the vesicles. Middle: CRG047 stain. Bottom: merge. The arrows point to internal vesicles within the larger GUV. CRG047 cannot diffuse through the outer bilayers towards the inner vesicles.

To study the CRG047 affinity for phospholipid membranes, increasing amounts of the compound in DMSO were added to a suspension of small unilamellar vesicles (SUV), and the fluorescence was recorded. Fluorescence intensity increased until saturation was reached (at ≈ 20 µM under our conditions) (Fig. 3c). The lower this concentration, the higher is the affinity of the compound for the vesicles. The time increase of bound CRG047 after mixing (Fig. 3d) indicated that the compound did not leave the membranes for several hours, and that additional compound was gradually transferred from micelles to membrane bilayers (Fig. 3d). In contrast with the amphipathic CRG047, the less polar Liss-RhPE reached equilibrium with the bilayers in the first minutes (Fig. 3d). The excitation and emission spectra of CRG047 (in SUVs) are shown in Figure 3e.

We also used fluorescence microscopy to explore the incorporation and retention of CRG047 (Fig. 3f, red) into preformed giant vesicles, some of them oligolamellar, previously stained with DiO in the first step of GUV formation (Fig. 3f, green). In the case of giant vesicles containing smaller vesicles labeled by DiO, CRG047 remained in the outermost vesicle membrane and was unable to diffuse across the bilayers (white arrows) to reach inner vesicles. This shows the suitability of the phospholipid-bound progestin to be used in live-cell experiments.

### CRG047 interaction with breast cancer cells membrane and activation of the kinase cascade

To further assess CRG047 incorporation to the cell membrane of living breast cancer cells, we incubated T47D cells with a physiologically relevant 10 nM concentration of CRG047 in PBS for 30 min. We visualized the incorporation of CRG047 by means of single molecule imaging of the fluorescent organic dye Nile Red under total internal reflection fluorescence microscopy (TIRF). We observed that the incorporation of CRG047 at such low concentrations yielded single molecules on the cell membrane of living T47D cells (see attached Video 1, Supplementary Figure Fig. S3). In order to efficiently visualize the incorporation of CRG047, we performed high-density single molecule maps (Fig. 4a) of Nile Red reflecting not only that CRG047 is incorporated on the cell membrane but that it also diffuses laterally as expected from a lipid bilayer intercalating dye.

**Figure 4.**
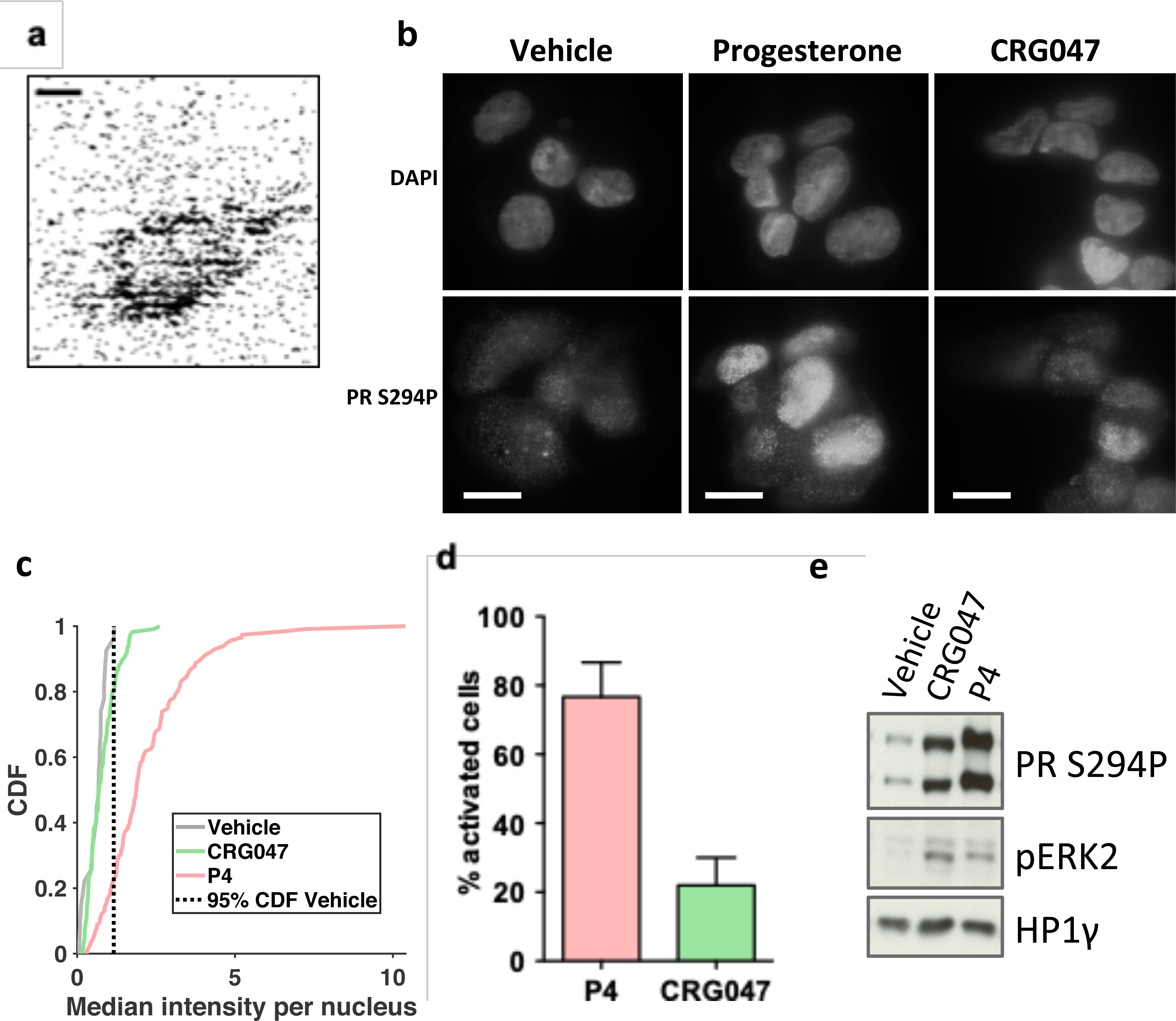
CRG047 interacts with and diffuses along the plasma cell membrane in T47D cells and activates MAPK pathway. a) High density map of the Nile-Red signals found in the movie (Fig. S3), showing the accumulation of fluorescence around the cell membrane in contact with the glass surface. Scale bar: 20 μm. b) Immunostaining. T47D cells imaged under HILO illumination in hormone-free conditions were exposed to vehicle, P4 or CRG047 as indicated. Upper panel: staining with DAPI for nuclei identification. Lower panel: PR Ser294 phosphorylation (PR S294P) signal after 30 min of exposure for each of the conditions. Scale bar: 10 μm. c) Cumulative density function (CDF) of the median intensity per nucleus for vehicle (grey), CRG047 (green) and P4 (magenta). The dashed line depicts 95% CDF for the vehicle condition. The nucleus with a median intensity above this value are therefore considered as activated by the respective compound. The intensity is in arbitrary units. A total of 27 cells were analyzed for vehicle, 117 cells for P4 condition and 116 cells for CRG047 condition. d) Histogram showing the percentage of cells activated upon exposure to progesterone or CRG047 (77% for P4; 22% for CRG047). Error bars indicate SD. e) Western blot on T47D extracts. T47D cells were treated with either vehicle, P4 or CRG047 (in the presence of 0.001% Tween20). Upper panel: blot against PR Ser 294 phosphorylation. Middle panel: blot against ERK2 p42/p44 phosphorylation. Lower panel: blot against HP1g as loading control. Source data: Middle panel (Figure 4 source data 1 and 2); upper and lower panels (Figure 4 source data 3 and 4).

We proceeded to assess the activation of the MAPK signalling cascade exposing T47D cells to P4 or CRG047 compared to vehicle. We first tested the activation of ERK-kinase activated phosphorylated PRS294P by means of a specific antibody using highly inclined and laminated optical sheet (HILO) fluorescence microscopy of an AlexaFluor 488 labeled secondary antibody. A significant fraction of cells showed a punctuated fluorescence pattern characteristic of PR activation (Fig. 4b). By calculating the cumulative density function (CDF) of the fluorescence intensity per nucleus of the antibody PRS294P (Fig. 4c), we determined that 77% or 22% of the cells were activated when exposed to 10 nM P4 or CRG047, respectively (Fig. 4d). We confirmed in cell extracts both ERK1/2 activation and PR phosphorylation when the cells were exposed to P4 or CRG047 (Fig. 4e). Thus, these experiments show that the compound CRG047 binds to the cellular membrane of the breast cancer cells and is able to activate the ERK-dependent PR signalling cascade.

### Functional response of the cells: gene regulation and cell proliferation through PR-chromatin interaction

To evaluate the nuclear response to the membrane-attached progestin, we used qRT-PCR to measure mRNA levels for genes known to be activated by P4, including *cMyc*, *Snail1*, *TiPARP* and *EGFR,* as well as the chromosomally integrated reporter *MMTV-luc*. All these genes were significantly activated by 10 nM P4 and also by 10 nM CRG047 (Fig. 5a).

**Figure 5.**
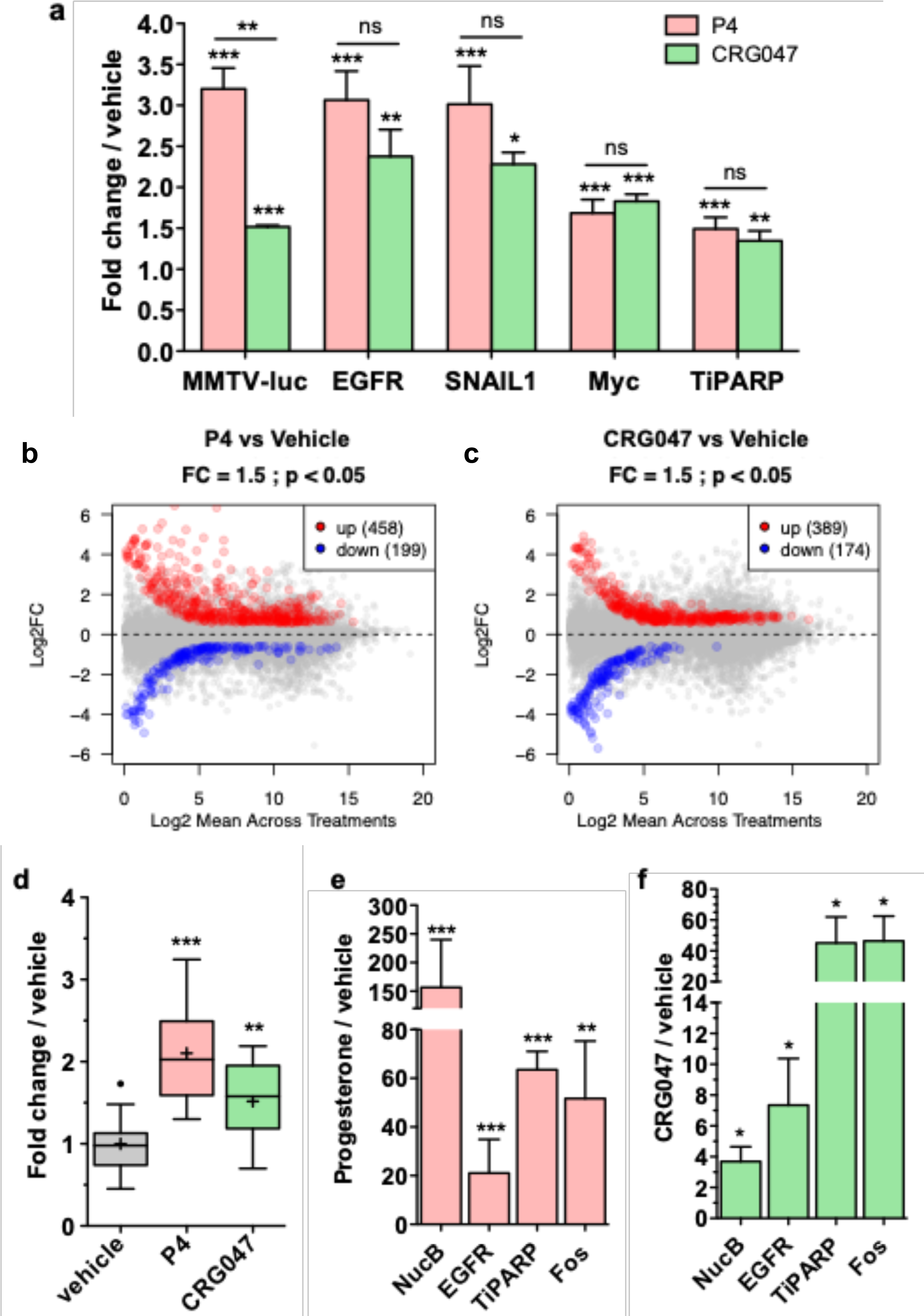
CRG047 activates transcription and proliferation through targeting PR to its HRE. For all graphs in this figure: All statistical analyses in this panel were done by 1 tail T-test comparing the indicated bar with vehicle treated samples or as indicated. *: p-value < 0.05; **: p-value < 0.01; ***: p-value < 0.001; ns: p-value > 0.05. a) RNA induction in T47D cells. Expression of progesterone target genes was analyzed by qRT-PCR. T47D cells in hormone-free conditions were exposed to vehicle, P4 or CRG047, RNA was extracted and qPCR analysis was performed for the indicated genes. For each gene, vehicle average levels were set to 1 and numbers indicate fold change over vehicle average. Values represent average ± SD of 3 independent experiments. b) Scatter plot of Pol II gene expression in T47D cells exposed to P4 and vehicle. The number of genes up- or downregulated in the P4 condition (−1.5 < FC > 1.5; adjusted pvalue < 0.05) is indicated. c) Scatter plot of Pol II gene expression in T47D cells exposed to CRG047 and vehicle. The number of genes up- or downregulated in the CRG047 condition (−1.5 < FC > 1.5; adjusted p-value < 0.05) is indicated. d) Cell proliferation was measured by BrdU incorporation in T47D cells following the addition of vehicle, P4 or CRG047. Boxplots represent the average of 12 independent experiments. Dot indicates outlayer. ‘+’ indicates average. e-f) ChIP against Progesterone receptor upon P4 (e) or CRG047 (f) induction of T47D cells. Values represent averages of 3 independent experiments ± SD.

We therefore performed a whole gene expression analysis of poly(A)-RNA by RNA-seq in cells exposed to 10 nM of either P4 or CRG047. After exposure to 10 nM P4, 458 genes were up-regulated and 199 were down-regulated, while upon exposure to 10 nM CRG047, 389 genes were up-regulated and 174 were down-regulated (Fig. 5 b,c). This indicates that CRG047 regulates a large number of genes comparable to those regulated by P4, though the extent of the regulation by CRG047 was less pronounced (Fig. S4 b, right panel).

As a further indicator of biological response to progestin, we measured the incorporation of BrdU into DNA, an indicator of cells entering the S phase of the cell cycle. We shortly exposed T47D cells to 10 nM of either P4, CRG047 or vehicle, followed by incubation in hormone-free medium. We found that both P4 and CRG047 significantly activated DNA synthesis compared to vehicle exposure (Fig. 5d).

To explore whether this gene regulation by CRG047 was mediated by binding of PR to its known HREs near regulated genes, we performed ChIP-qPCR at P4 regulated genes after exposure to 10 nM P4 or CRG047. We found a significant binding of PR to the regulatory PREs in five P4 responsive genes including the promoter region of the MMTV-luc reporter (NucB). The effects of CRG047 were in the same direction than those of P4 (Fig. 5e for P4 and f for CRG047). Therefore, we conclude that progestin-like compound can activate the cell response in a similar manner as progesterone.

### Neither CRG047 nor its putative hydrolysis product are detected within cell nuclei

To explore whether CRG047 did enter the cell nuclei, T47D cells were exposed shortly to 10 or 100 nM P4 or CRG047 and total cell extracts and nuclear extracts were subjected to quantitative mass spectrometry (MS). Analyses were performed in parallel for CRG047 (Fig. S5 a-d) or P4 (Fig. S5 e-h). Although the limit of detection was below 1 pmol (not shown), neither CRG047 nor P4 could be detected in nuclear extracts (compare panels a, b for CRG047 and panels e, f for P4). Similar analyses were performed with the progesterone analogue R5020 (promegestone) (panels i-l).

To exclude that CRG049 – the steroid metabolite of CRG047 hydrolysis – could be responsible for some of the observed effects, the levels of CRG047 and CRG049 in total extracts from cells exposed to 10 nM and 100 nM CRG047 were measured by MS. The results are shown in Figure S6. While CRG047 was clearly detected in the cell extracts (Fig. S6a), CRG049 was indistinguishable from noise at the same concentrations both in nuclear and total cell extracts (Fig. S6a). Thus, it seems unlikely that the hydrolysis of CRG047 was responsible for the observed effects in breast cancer cell function.

### Palmitoylated PR at Cys 820 is required for the response to CRG047

Given that other P4 binding proteins are known to inhabit the cell membrane,^26^ in order to ensure that the mbPR was responsible for the action of CRG047, we inhibited palmitoylation with 10 µM 2-bromopalmitate (2-Br).^9^ We found that this treatment effectively blocks the response of T47D to both P4 and CRG047 (Fig. 6a). In addition, we used a PR mutant in which the palmitoylation site (Cys-820) was mutated to Ala (PR C820A). This mutant PR does not integrate into the cell membrane.^9^ We found that, when expressed in T47D cells deprived of wtPR that express a single MMTV promoter copy (TYML),^27^ PR C820A could not mediate the cell response to 10 nM P4 or CRG047 in terms of activation of P4-responsive genes (Fig. 6b and Fig. S7a). Similar results were obtained when wtPR or PR C820A mutant were co-expressed along with the MMTV promoter plasmid in another cancer cell line defective in progesterone receptor (U2OS cells) and exposed to P4 or CRG047 (Fig. S7b). In addition, with U2OS cells expressing either the wtPR or the mutant PR C820A, only the wtPR-expressing cells activated the ERK kinase and led to PR phosphorylation (Fig. 6c and d). We conclude that CRG047 action on T47D cells require the anchoring of PR to the cell membrane via palmitoylation.

**Figure 6.**
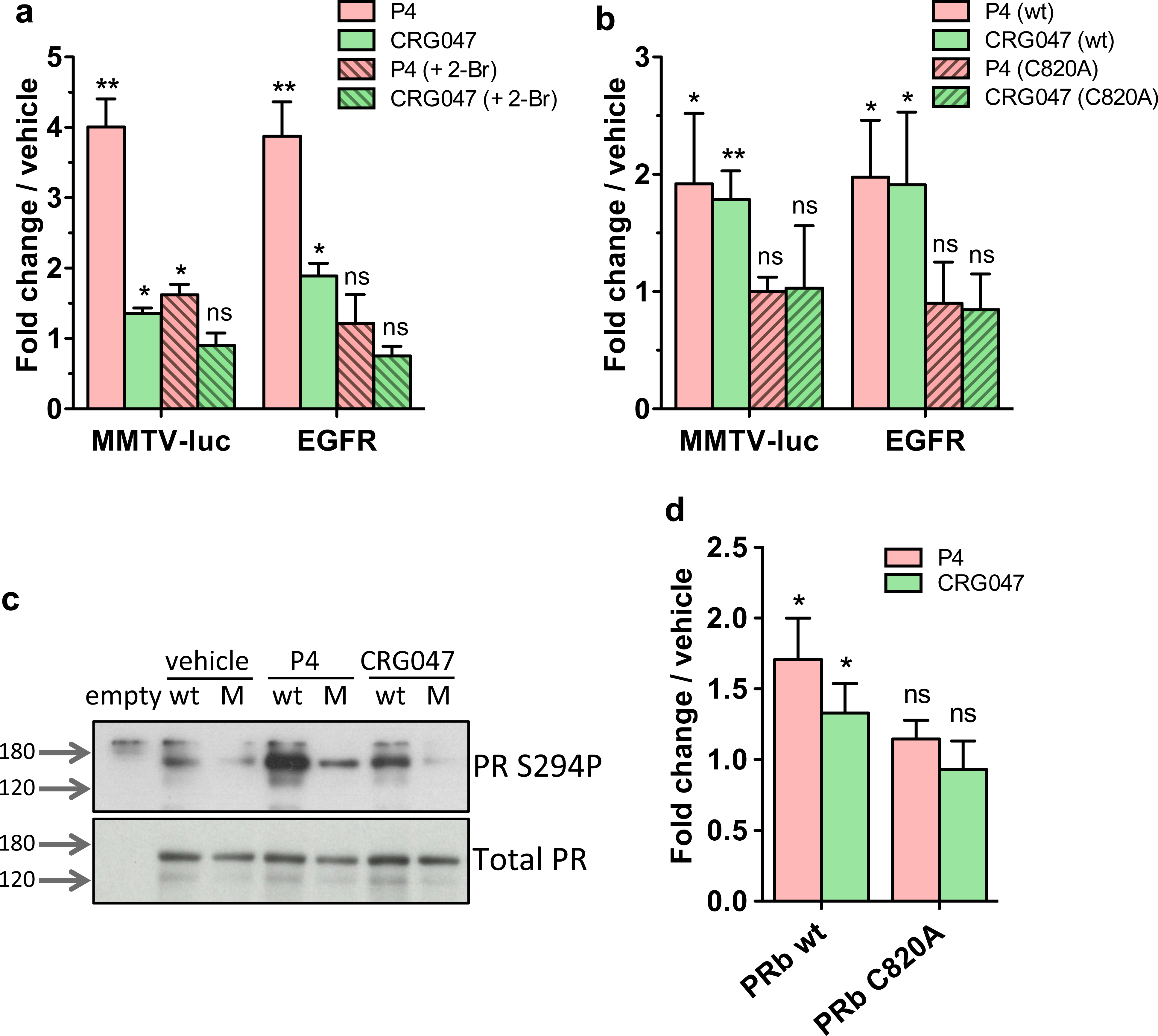
CRG047 action is dependent on PR palmitoylation. For all graphs in this figure: Statistical analyses in this panel were done by 1 tail T-test comparing the indicated induction (P4 or CRG047) with vehicle in triplicates. *: p-value < 0.05; **: p-value < 0.01; ns: p-value > 0.05. a) Palmitoylation inhibition. T47D cells were incubated for 6h with 2-bromopalmitate 10 μM (2-Br, dashed bars) prior to induction with the indicated compounds for 3 h. RNA signal for indicated genes is shown relative to vehicle induction. Bars represent the average ± SD of 3 independent experiments. b) PR-negative breast cancer cell line T47D-Y stably expressing a single copy of the MMTV promoter (TYML) were transfected with wild type PRb (solid bars) or the palmitoylation mutant PRbC820A (dashed bars) plasmids and incubated with P4 or CRG047 as indicated. Average ± SD of 3 independent experiments is shown. Fold changes are relative to the average response of cells incubated with vehicle. c) Western-blot against PR Ser294 phosphorylation (upper panel) or total progesterone receptor as loading control (lower panel). U2OS cells were transfected with empty plasmid, or with plasmids for the expression of either wild type PRb (wt) or the palmitoylation mutant PRb C820A (M) and incubated with vehicle, P4 or CRG047. Fold changes are relative to the average response of cells incubated with vehicle. d) Histogram of 3 independent experiments (average ± SD) performed as in panel c. Signal was captured by pixel density and normalized against total PR signal as loading control. Source data: upper panel (Figure 6 source data 1 and 2); lower panel (Figure 6 source data 3 and 4).

## DISCUSSION

The purpose of this work was to explore whether a membrane-attached progestin derivative could activate the cellular response of breast cancer cells to progestins without the requirement of the hormone penetrating into the cell and binding the intracellular PR. The results are compatible with this hypothesis, although we do not exclude other possible explanations.

In support of our hypothesis, the above results show that the progestin linked to a stable phospholipid (CRG047) binds the mbPR with high affinity and remains attached to the cell membrane during the time of exposure of the cells (30 min). Chemically, the amide bond linking progestin to the phospholipid is not hydrolyzed during exposure of the cells to CRG047 as the resulting compound (CRG049) cannot be detected in the cell extracts. Our designed compound CRG047 induces the activation of the ERK kinase pathway leading to phosphorylation of the intracellular PR at S294, which results in PR binding to genomic PREs, regulation of the associated progesterone sensitive genes, and entry into the S phase of the cell cycle. Finally, our data show that anchoring of PR to the cell membrane via palmitoylation at C820 is essential for CRG047 action.

In fact, it has never been demonstrated that the PR that binds to chromatin PREs upon exposure of breast cancer cells to 10 nM or to 50 pM P4 is actually carrying a bound hormone ligand,^11, 28^ and we cannot demonstrate the presence of P4 in nuclear extracts of cells exposed to 10 nM P4. Thus, the main function of the hormone ligand could be to activate the kinase pathway via mbPR, leading to phosphorylation of the intracellular PR, detachment from chaperones, dimerization and transfer to the cell nucleus.

One limitation of our study is that our assays are focused on the initial response of the cells to hormone, as we did not dare to expose the cells to CRG047 for longer than 30 min. The reason was that we wanted to exclude the possibility that the mbPR could be reaching the cell interior during the metabolic processing of the membrane. This short exposure time may also be responsible for the low percentage of cells (∼20%) that responded to CRG047, since judging from the experiments with phospholipid vesicles, binding of CRG047 increased after longer incubation times, most likely due to the fraction of compound forming micelles not available for binding to the cell membranes being released. Nevertheless, even this short exposure time was sufficient to initiate the cell cycle in the next 16 hours.

A further limitation of our study is that we have not explored the binding of the compound CRG047 to other known membrane-attached progestin binding proteins.^26^ PAQRs are seven-transmembrane-helix proteins that bind progesterone and adiponectin Q, and are expressed in many tissues including the nervous system. To our knowledge, these PAQRs receptors are not expressed in the T47D cell lines we have used for this study. Progesterone Membrane Components 1 and 2 (PGRMC1/2) are single-pass transmembrane receptors that bind progesterone, cortisol, androgens, cholesterol and heme.^29^ We have not explored the possible function of PGRMCs in response to progestins, but we have shown through the palmitoylation of PR mutants that the effects of CRG047 require the binding of PR to the membrane.

Finally, the strategy used in this study has not been tested to experiments in living animals, as the phospholipid-containing compounds we used may bind many components of the blood and other fluids, making difficult their interaction with the target cells at the required low concentrations.

In any case, the relevance of the mbPR for the hormone response suggests that it could be a novel target for endocrine therapy of breast cancers. For instance, developing selective inhibitors of its palmitoylation, or blocking its interaction with the SRC kinase. It is very likely that ERα is bound to the cell membrane by a similar mechanism, since it shares the conserved palmitoylation signal.^9^ Thus, a selective palmitoylation inhibitor would block the action of both estrogens and progestins. This strategy would require a better understanding of the complex formed by mbPR with ERα, the SRC kinase, and other associated proteins.

## EXPERIMENTAL SECTION

### Materials

Progesterone **and R5020 were** obtained from Perkin-Elmer. Lipofectamine 2000 Transfection reagent was from Thermo Fischer. 2-Bromopalmitate was acquired from Sigma. pcDNA3.1 (+) SNAP-GFP-PRb wt and pcDNA3.1 (+) SNAP-GFP-PRb C820A plasmids were generated from pTET-Splice SNAP-GFP-PRb plasmid kindly given by Gordon Hager, subcloned into pcDNA3.1(+) backbone. Mutagenesis of the palmitoylation site (aa Cys 820 into Ala) was done by GenScript Biotech (Netherlands) B.V. pAGEMMTVLu(MMTV-luc) was used to assess expression of the progesterone responsive promoter MMTV in U2OS cells.

### Compounds synthesis

For a full account of the synthesis of all new compounds used in this work see Supporting Information.

### Cell culture

T47D cells encompassing a single copy of the MMTV promoter were grown in RPMI 1640 (Gibco) supplemented with 10% fetal bovine serum (FBS), 100U/ml penicillin, 100ug/ml streptomycin, 2mM L-glutamine and 10 µg/ml insulin. TYML cells and U2OS cells were grown in DMEM (Gibco) supplemented with 10% fetal bovine serum (FBS), 100U/ml penicillin and 100ug/ml streptomycin.

### Hormone induction

For hormone induction experiments, cells were plated in DMEM/RPMI media without phenol red, supplemented with 10% dextran-coated charcoal-treated FBS and incubated for 48 h. Cells were synchronized 16 h prior to hormone induction by starvation in serum-free medium.

For inductions, cells were incubated with 10 nM P4, 10 nM CRG047 or vehicle (ethanol) as indicated. Progesterone or progesterone-like compounds were resuspended in ethanol to 2 mM concentrations, and diluted with PBS to the indicated concentrations for cellular induction. Vehicle exposed cells were treated in the same manner relative to ethanol final concentrations.

For longer time points, hormone dilutions were replaced by fresh hormone-free media after 30 min of induction.

### Transient transfections

Wild type and mutant PR constructs were transfected into TYML or U2OS cells using Lipofectamine 2000 Transfection reagent (Thermo Fischer) following manufacturer’s instructions. 300.000 cells were plated in 35 mm wells and transfected with 0.5 µg of plasmid/well. U2OS cells were co-transfected with MMTV-luc plasmid for expression inducing experiments. Hormone induction experiments were carried 48 h post-transfection.

### Palmitoylation inhibitor

T47D cells were exposed to 10 μM 2-Bromopalmitate (Sigma) for 6 h prior to hormone induction.

### Microscale thermophoresis (MST)

Histidine-tagged Progesterone Receptor (PR) B was purified as previously described^23^ with a modification. Activation with a progesterone analog was skipped in order to avoid competition with the compounds to be analyzed. Progesterone-like compounds affinity for Progesterone Receptor was measured by Microscale Thermophoresis (MST). Non-labeled His-PRb recombinant protein or GST-PR Ligand Binding Domain (LBD), purified as described^25^ (starting concentration 1750nM), were titrated to a fixed concentration of Nile-Red-containing progesterone derivative compounds (2 µM) in a buffer containing 20mM tris pH 7.4, 90mM NaCl, 1mM DTT, 10% glycerol and 0.01% Tween 20. MST data was recorded at 20°C using the blue LED at 10% and IR-Laser at 40% according to manufacturer’s instructions. The thermophoresis change as normalized fluorescence (ΔFnorm in [‰]) was plotted against the concentration of the unlabeled protein and fitted according to the law of mass action.

### Fluorescence spectroscopy measurements

Lipids were mixed in chloroform:methanol (2:1), and the solvent was evaporated to dryness under a stream of N_2_. Then the sample was kept under vacuum for 2 h to remove solvent traces, and the lipids were swollen in the appropriate buffer, 25 mM HEPES, 150 mM NaCl, pH 7.4. Small unilamellar vesicles (SUV) were obtained by sonicating the swollen lipid suspensions with a probe-type Soniprep 150 sonicator (MSK, London, U.K.).^30^ The lipid concentration was determined by phosphate analysis.^31^ SUV were mixed with CRG047, and the mixture was left to equilibrate (usually 1 h). Fluorescence was measured in a QuantaMaster 40 spectrofluorometer (Photon Technology International, Lawrenceville, NJ) (λex=550 nm, λem=625 nm). The measurements were performed at a constant temperature of 22°C. The extinction coefficient and quantum yield of CRG047 were obtained as described.^16^

### Confocal microscopy

Giant unilamellar vesicles (GUV) were prepared by electroformation on a pair of platinum (Pt) wires by a method first developed by Angelova and Dimitrov,^32^ modified as described previously.^33^ Lipid stock solutions were prepared in 2:1 (v/v) chloroform/methanol at 0.2 mg/ml, and appropriate volumes of each one were mixed. Labelling was carried out by pre-mixing the desired fluorescent probes with the lipids in organic solvent. 3,3’-Dioctadecyloxacarbocyanine Perchlorate (DiO) was used as a general marker for lipid membrane. The average concentration of individual fluorescent probes in each sample was 0.5 mol%. 2.5 μl organic solutions of lipid mixtures containing the fluorescent probes were deposited on Pt wires. The Pt wires were placed under vacuum for 2 h to completely remove the organic solvent.

One side of the chamber was then sealed with a coverslip. 500 µl assay buffer, prepared with high-purity water (Millipore SuperQ) heated at 37°C was added to the chamber until it covered the Pt wires, and connected to a TG330 function generator (Thurlby Thandar Instruments, Huntingdon, UK). AC field was applied in three steps, all of them performed at 37°C: 1) frequency 500 Hz, amplitude 220 mV (35 V/m) for 5 min; 2) frequency 500 Hz, amplitude 1900 mV (313 V/m) for 20 min; 3) frequency 500 Hz, amplitude 5.3 V (870 V/m) for 90 min. The temperatures used for GUV formation correspond to those at which the different membranes display a single fluid phase.

GUVs were imaged with confocal microscopy using a Nikon D-eclipse C1 confocal system (Nikon corporation, Tokyo, Japan). The excitation wavelengths used for excitation were 488 nm for DiO and 561 nm for CRG047. Fluorescence emission was retrieved at 500-530 nm for DiO and at 573-613 nm for CRG047. When required, 2 µM CRG047 in DMSO was added to study its effect on the GUVs. All these experiments were carried out at 22° C. Image treatment was performed using EZ-C1 3.20 software (Nikon Inc., Melville, NY).

### Immunostaining

Cells were plated in chambered cover glass. After induction, cells were fixed with 3% PFA in PBS for 10 min at RT, rinsed 3 times with PBS and blocked with 3% BSA, 0.2% Triton X-100 in PBS for 1h. Primary antibody incubation was done in blocking buffer for 1 h at rt (anti PR Ser294Phospho, ab61785, Abcam). Primary antibody was washed 3 times in wash buffer (0.2% BSA, 0.05% Triton X-100 in PBS) and secondary (Alexa fluor 488 antibody, Thermo Fisher #A-11008) was added in blocking buffer for 40 min. Cells were washed again 3 times and DAPI was added for 1 min in a 10.000x dilution in PBS and washed 3 times.

### Single molecule sensitive fluorescence microscopy

HILO or TIRF imaging were performed in a Nikon N-STORM 4.0 microscope system equipped with a TIRF 100x, 1.49 numerical aperture objective (Nikon, CFI SR HP Apochromat TIRF 100XC Oil). The dyes Nile Red or Alexa Fluor 488 were excited with a continuous 561 nm or 488 nm laser line, respectively. The emission fluorescence was collected and projected into an EM-CCD Andor Ixon Ultra Camera at a frame rate of 15 ms. The pixel size of the camera is 160 nm. High density maps were produced by collapsing (≈ 7700) single molecule localizations of Nile-Red labeled CRG047, incorporated on the cell membrane of a living T47D cell, into 1 single frame.

### Immunostaining quantification methodology. Quantification of activated cells using MATLAB

To quantify the percentage of activated cells under different conditions, we measured the median intensity of the fluorescence emission of each nucleus (Fig. S8a). In order to obtain such fluorescence intensity values, we first took the DAPI images (Fig. S8b) and used the MATLAB tool Assisted Free Hand to obtain the nucleus masks for each cell (Fig. S8c) by defining the contour of each nucleus. Second, we applied the masks for all the S294P images (Fig. S8d) and computed the median intensity of the fluorescence intensity for each nucleus. Finally, from the histogram distributions of median intensity we obtained the cumulative density function (CDF) of ethanol-exposed samples and took the 95% median intensity, which we use to define the inactive/active threshold. Thus, those cells with a median intensity per pixel larger than the threshold are considered to be active. For the analysis, we took 27 cells exposed to ethanol as a negative control, 117 cells exposed to P4 and 116 cells exposed to CRG047.

### Expression analysis

Cells were induced for 30 min as described and after 3 h RNA was isolated using Qiagen RNAeasy kit following manufacturer’s instructions. RNA concentration was measured with a Qubit fluorometer and RNA was subjected to Bioanalyzer for quality control.

For gene expression analysis, RNA (250 ng) was subjected to cDNA synthesis using the qScript cDNA Synthesis kit (Quanta Biosciences). qPCR was carried out using the LightCycler FastStart DNA Master SYBR Green I kit (Roche), and specific primers selected from the list of available designed primers at Primer Bank (https://pga.mgh.harvard.edu/primerbank).^34^ As a reference gene, *GAPDH* was used. Data shown represents the mean of at least 3 independent experiments. Fold change was calculated using Δ-Δct method. Specific primer sequences are listed in the table below.

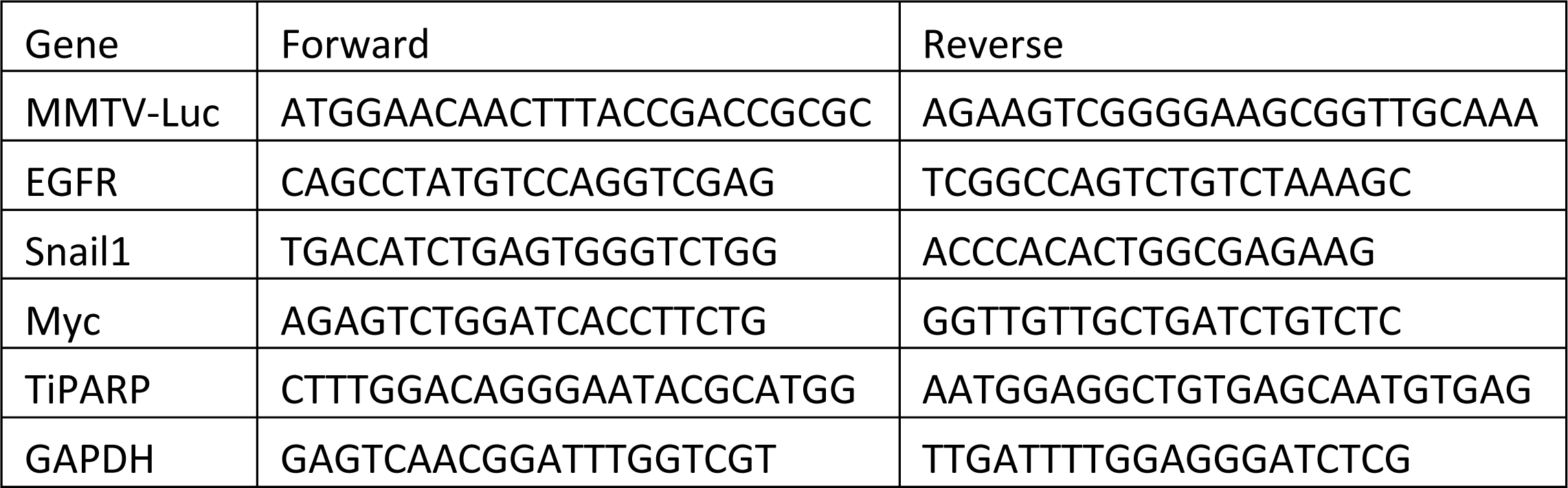

### RNA-seq, Pipeline and Differential Gene Expression Analysis

RNA-seq preparation and analysis was done as described by Ferrari.^35^ Libraries were prepared using 1 μg polyA+ RNA by PCR amplification of cDNA with bar-coded primers using the Illumina TruSeq kit at the CRG Genomic Facility. Libraries were sequenced using Illumina HIseq-2500 to obtain pair-ended (PE) 100-base-long reads.

Sequencing adapters and low-quality ends were trimmed from the reads with Trimmomatic, using the parameters values recommended by Bolger^36^ and elsewhere (https://goo.gl/VzoqQq) (trimmomatic PE raw_fastq trimmed_fastq ILLUMINACLIP:TruSeq3-PE.fa:2:30:12:1:true LEADING:3 TRAILING:3 MAXINFO:50:0.999 MINLEN:36). The trimmed reads were aligned to GRCh38^37^ using STAR.^38^

First, the genome index files for STAR were generated with: star–runMode genomeGenerate–genomeDir GENOME_DIR–genomeFastaFiles genome_fasta– runThreadN slots–sjdbOverhang read_length–sjdbGTFfile sjdb–outFileNamePrefix GENOME_DIR/ where genomeFasta is the FASTA file containing the GRCh38 sequence downloaded from the University of California Santa Cruz (UCSC) Genome Browser, excluding the random scaffolds and the alternative haplotypes; and sjdb is the GTF file with the GENCODE’s V24 annotation.

Second, trimmed reads were aligned to the indexed genome with: star–genomeDir GENOME_DIR/–genomeLoad NoSharedMemory–runThreadN slots–outFilterType “BySJout”–outFilterMultimapNmax 20–alignSJoverhangMin 8–alignSJDBoverhangMin 1–outFilterMismatchNmax 999–outFilterMismatchNoverLmax 0.04–alignIntronMin 20– alignIntronMax 1000000–alignMatesGapMax 1000000–readFilesIn read1 read2– outSAMtype BAM SortedByCoordinate–outTmpDir TMP_DIR/–outFileNamePrefix ODIR1/$sample_id.–outWigType bedGraph–readFilesCommand zcat (https://docs.google.com/document/d/1yRZevDdjxkEmda9WF5-qoaRjIROZmicndPl3xetFftY/edit?usp=sharing).

To avoid mis-alignment, we only accepted uniquely mapped reads and we focused on AE placed within intergenic regions (at least 5 kb away from any human TSS) or on the opposite strand of a known annotated transcript.

Differences in gene expression were calculated using a DESeq2. Genes with fold change (FC) ± 1.5 (p value < 0.05; FDR < 0.01) were considered as significantly regulated.

### Cell proliferation

T47D cells were plated for hormone induction in a 96-well plate at 1×10^4^/well density and induced with either vehicle, P4 or CRG047 for 30 min, followed by 16 h of culture in hormone-free medium, and measured BrdU incorporation into DNA. ELISA BrdU Colorimetric assay (Roche) was performed according to manufacturer’s instructions. Average results from 12 independent experiments are shown.

### Mass spectrometry

#### Chemicals, reagents and lipid standards

Optima® LC/MS grade water, methanol, acetonitrile, 2-propanol and formic acid were from Fisher Scientific (Fair Lawn, NJ, USA). Ammonium formate for mass spectrometry and toluene of analytical standard quality were from Sigma-Aldrich (Sigma Chemical Co., St Louis, MO, USA). Mass spectrometer calibration solutions, Pierce LTQ Velos ESI Positive Ion Calibration Solution and Pierce LTQ Velos ESI Negative Ion Calibration solutions, were provided by Thermo Fisher Scientific (Waltham, MA).

#### Sample preparation for analysis

Cells were grown and induced as above. After induction cells were collected and total extract was obtained in lysis buffer. The Bligh and Dyer protocol was applied to extract the lipidic fraction.^39^ Dried extracts were reconstituted with 50 µl MeOH:Toluene (9:1,v/v) and vortexed for 30 s before injection.

#### UHPLC-HRMS analysis

CRG047, CRG049, progesterone and promegestone analyses were performed in an ULTIMATE 3000 (Thermo Scientific) UHPLC system coupled to a Qexactive HF-X (Thermo Scientific) quadrupole-orbitrap mass spectrometer. A reverse phase column (Acquity UPLC HSS T3, 100 x 2.1 mm, 1.8 µm) and a precolumn (Acquity UPLC HSS T3 1.8 µm VanGuard^TM^) were used. Mobile phases and UHPLC settings are shown in Table S2.

HRMS was operated using the electrospray source in positive mode. The ionization source parameters are shown Table S2. MS data acquisition was performed in mode t-SIM/dd-MS^2^ with inclusion list containing the parent ion mass of target analytes. Data was processed using FreeStyle^TM^ 1.6 SP1 (Thermo Scientific).

#### Identification and quantification

Identity of each compound was confirmed by MS/MS in all samples and quantification was performed by the external standard method, using integration areas from extracted SIM ion chromatograms of the calibration regression of CRG047, CRG049, progesterone and promegestone internal standards. For this purpose, eight solutions containing the internal standards in concentrations ranging from 0.05 to 1000 nM were analyzed in triplicate. Method sensitivity was studied determining calibration curve slopes and detection (LOD) and quantification (LOQ) limit values. The LOD and LOQ for each internal standard were calculated as the lowest concentration that results in a CV ≤ 20%.

#### Chromatin immunoprecipitation

ChIP assays were performed as described^40^ using Progesterone Receptor antibody [Alpha PR6] (Abcam ab2765).

Quantification of chromatin immunoprecipitation was performed by real time PCR using Roche Lightcycler 2.0, with the specific oligos for the determined regions listed below.

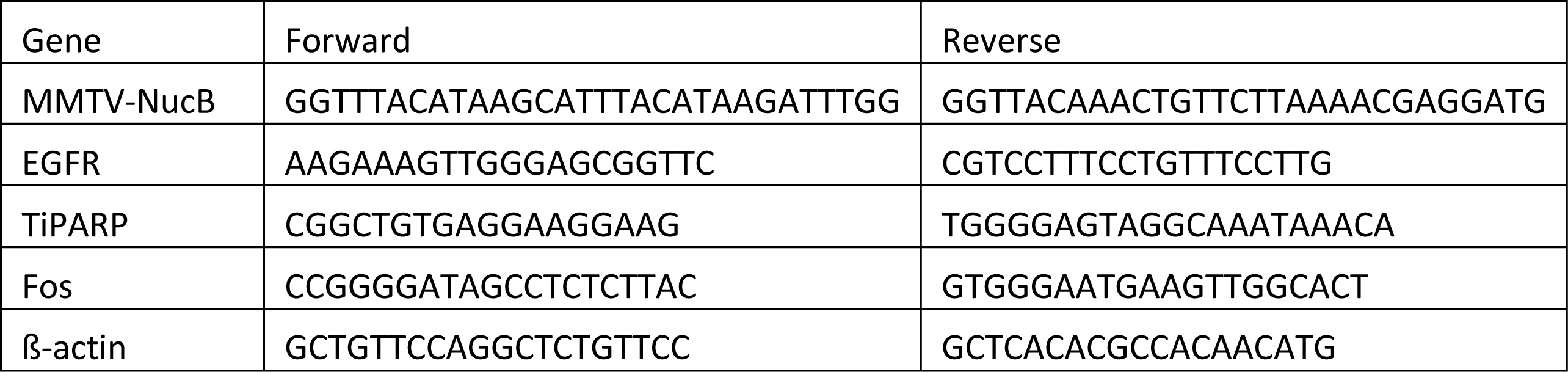

The fold enrichment of target sequence in the immunoprecipitated (IP) compared to input (Ref) fractions was calculated using the comparative Ct method (the number of cycles required to reach a threshold concentration) with the equation 2^Ct(IP)-Ct(Ref)^. Values were corrected by the human β-actin gene and presented as relative abundance over vehicle.

#### Immunoblot

Induced cells were washed twice in cold PBS and collected in lysis buffer (25 mM Tris pH 7.5, 1% SDS, 1 mM EDTA, 1 mM EGTA and protease inhibitors). Protein extracts were quantified and equal amounts were loaded in an 8% SDS-PAGE, transferred into Nitrocellulose membrane, blotted against indicated primary antibodies and HRP-conjugated correspondent secondary antibodies. Primary antibodies: Total Progesterone Receptor antibody [Alpha PR6] (Abcam ab2765), Progesterone Receptor Ser294Phospho, (Abcam ab61785), Phospho-p44/42 MAPK (Erk1/2) (Thr202/Tyr204) (Cell Signaling #9106S), HP1 gamma (Millipore 05-690).

## Supporting Information

Experimental procedures and spectroscopic data for all new compounds (1H NMR, 13C NMR, 31P NMR), including images of NMR spectra.

## ACKNOWLEDGEMENTS

This work was supported in part by the Spanish Ministerio de Ciencia e Innovación (MCI), Agencia Estatal de Investigación (AEI) and Fondo Europeo de Desarrollo Regional (FEDER) (grant No. PGC2018-099857-B-I00), by the Basque Government (grants No. IT1625-22 and IT1270-19), by Fundación Ramón Areces (CIVP20A6619), by Fundación Biofísica Bizkaia, and by the Basque Excellence Research Centre (BERC) program of the Basque Government. This research was supported by European Research Council (Project “4D Genome” 609989), the Ministerio de Economía y Competitividad (Project G62426937) and the Generalitat de Catalunya (Project AGAUR SGR 575 and AGAUR 2019PROD00115/IU68-016733), European Research Council -Proof Of Concept (Project “Impacct” 825176).

## SUPPLEMENTARY FIGURES

**Figure S1 ; Figure 1 - Figure supplement 1.**
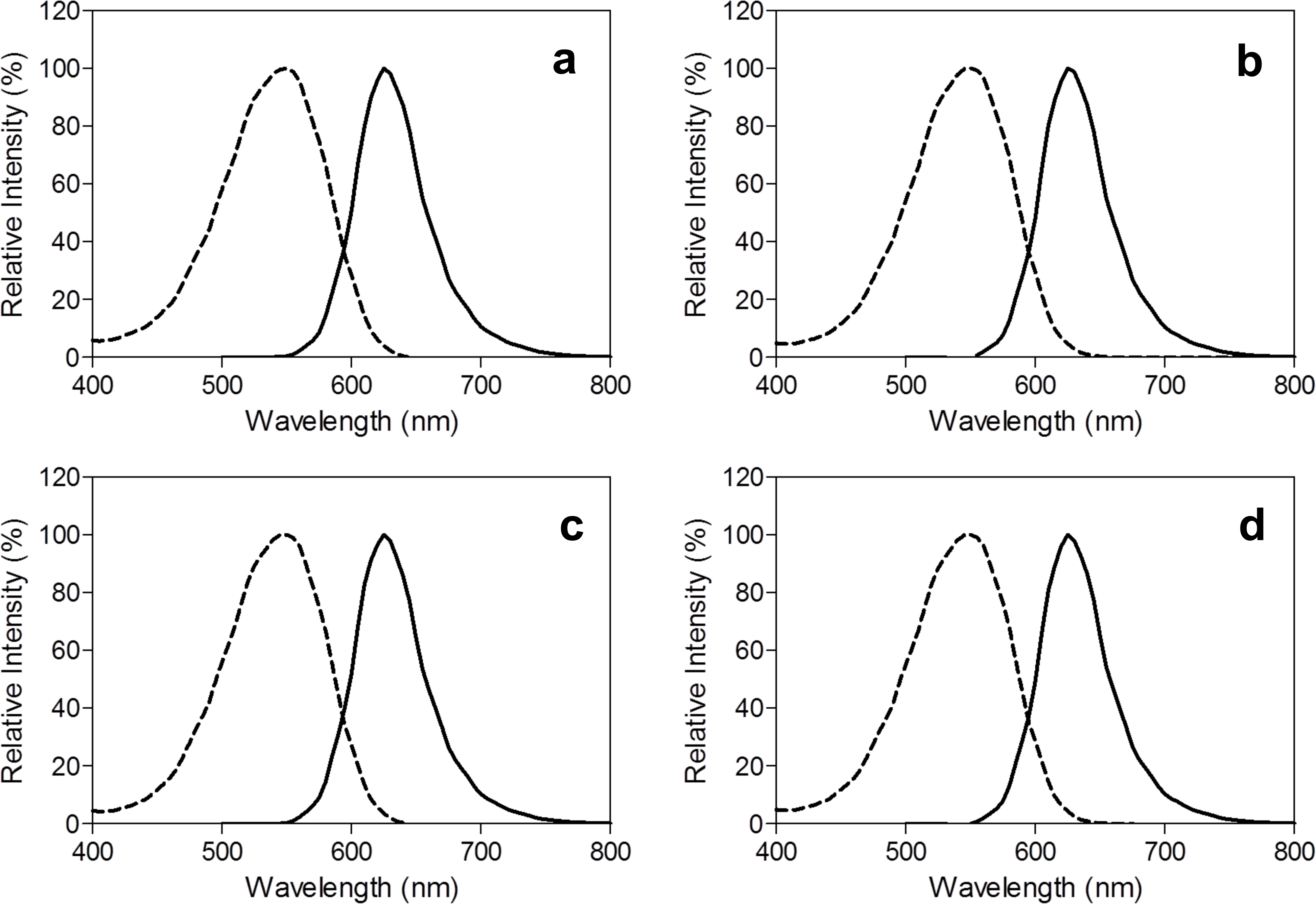
Excitation (dashed line) and emission (bold line) spectra of compounds CRG033 (a), CRG047 (b), CRG034 (c) and CRG048 (d) in EtOH. λ_ex_/λ_em_ were 550/625 nm for all compounds.

**Table S1.** Spectroscopic properties of probes CRG033, CRG034, CRG047 and CRG048 in EtOH.

| Compound | $\lambda_{\text{ex}}/\lambda_{\text{em}}$ (nm) | $\epsilon$ ($\text{M}^{-1}\cdot\text{cm}^{-1}$) | $\Phi_{\text{F}}$ |
| --- | --- | --- | --- |
| **CRG033** | 550/625 | $3.64 \times 10^4$ | 0.33 |
| **CRG034** | 550/625 | $4.17 \times 10^4$ | 0.39 |
| **CRG047** | 550/625 | $4.22 \times 10^4$ | 0.31 |
| **CRG048** | 550/625 | $4.24 \times 10^4$ | 0.31 |

**Figure S2; Figure 2 - Figure supplement 1.**
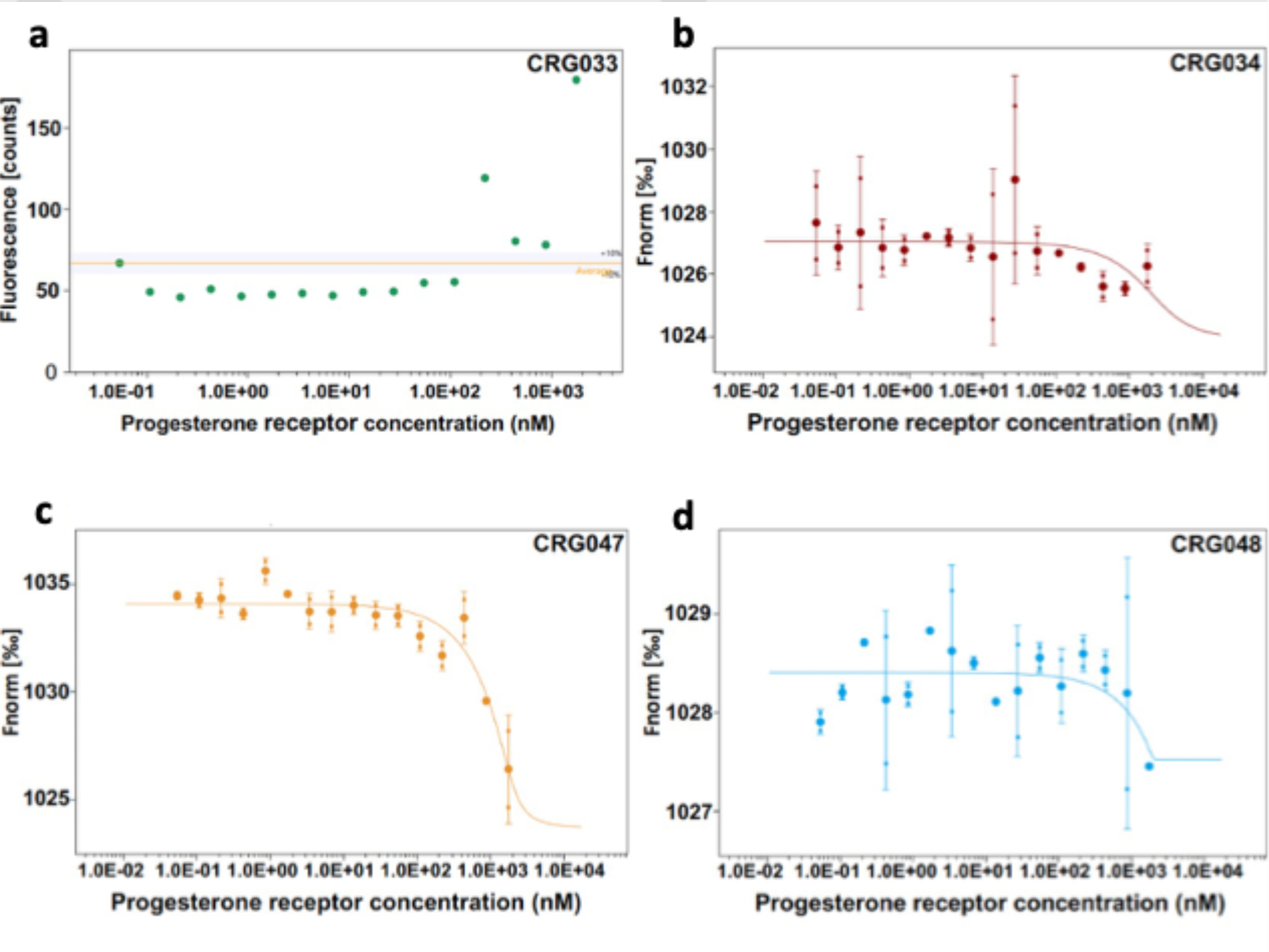
a) Microscale thermophoresis (MST). Initial fluorescence representation of CRG033 interaction with full length PR. b-d). MST dose/response representation of 2 readings of the same experiment (average ± SD is shown) of the indicated compounds. Normalized fluorescence plotted against increasing concentrations of PRb. 2 μM concentrations of the compounds were assessed with increasing concentrations of PRb.

**Figure S3; Figure 4 - Figure supplement 1.**
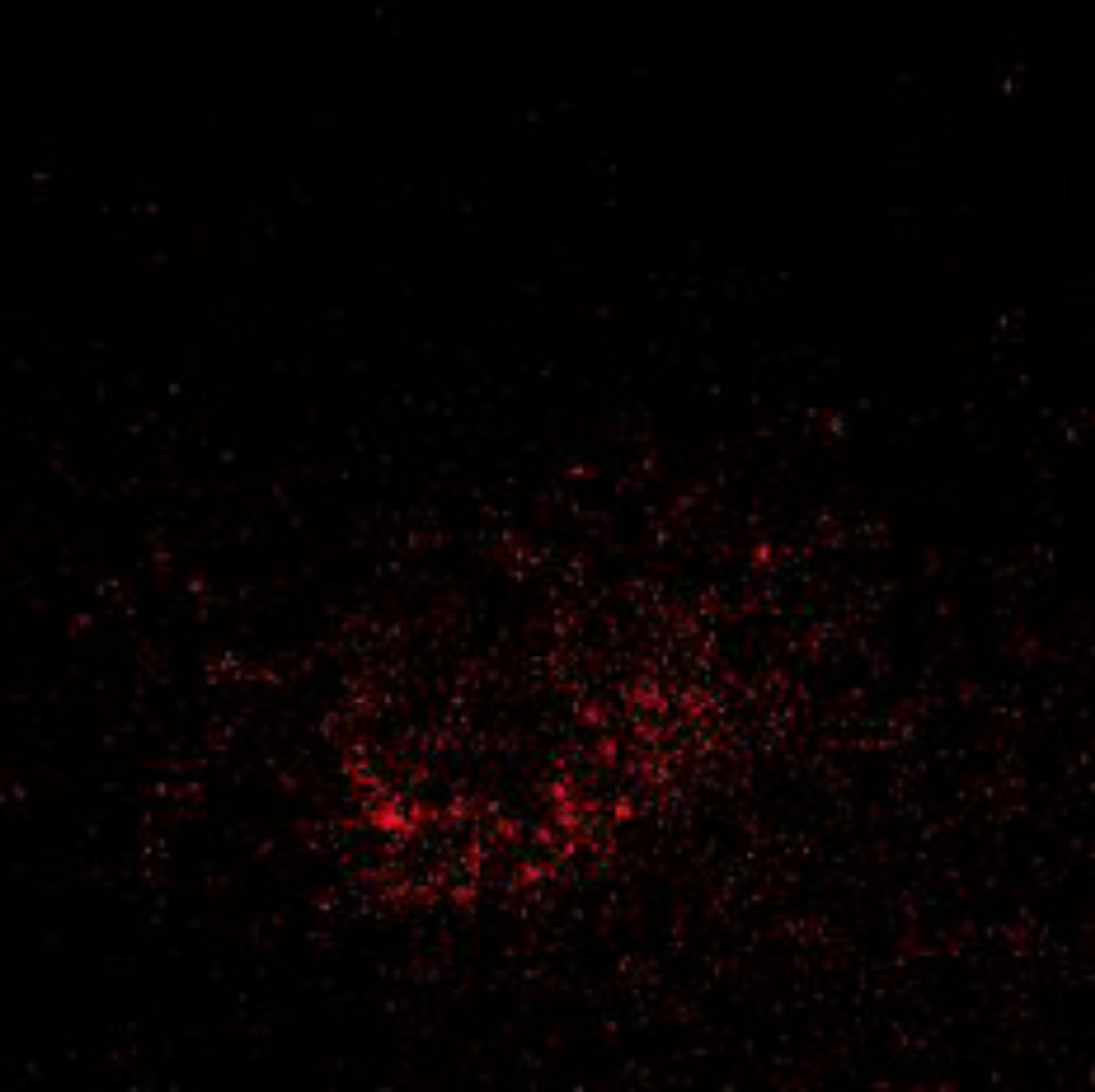
TIRF imaging of single molecules diffusing on the cell membrane of breast cancer cells exposed to 10 nM of the progesterone derivative CRG047 dissolved in PBS. Single molecules are detected with a custom software and projected into 1 single frame to build up cartography maps as shown in Figure 4a. (Note to editor: First frame shown, video attached as Supporting Information (SI) - “Supplementary Figure S3”.

**Figure S4; Figure 5 - Figure supplement 1.**
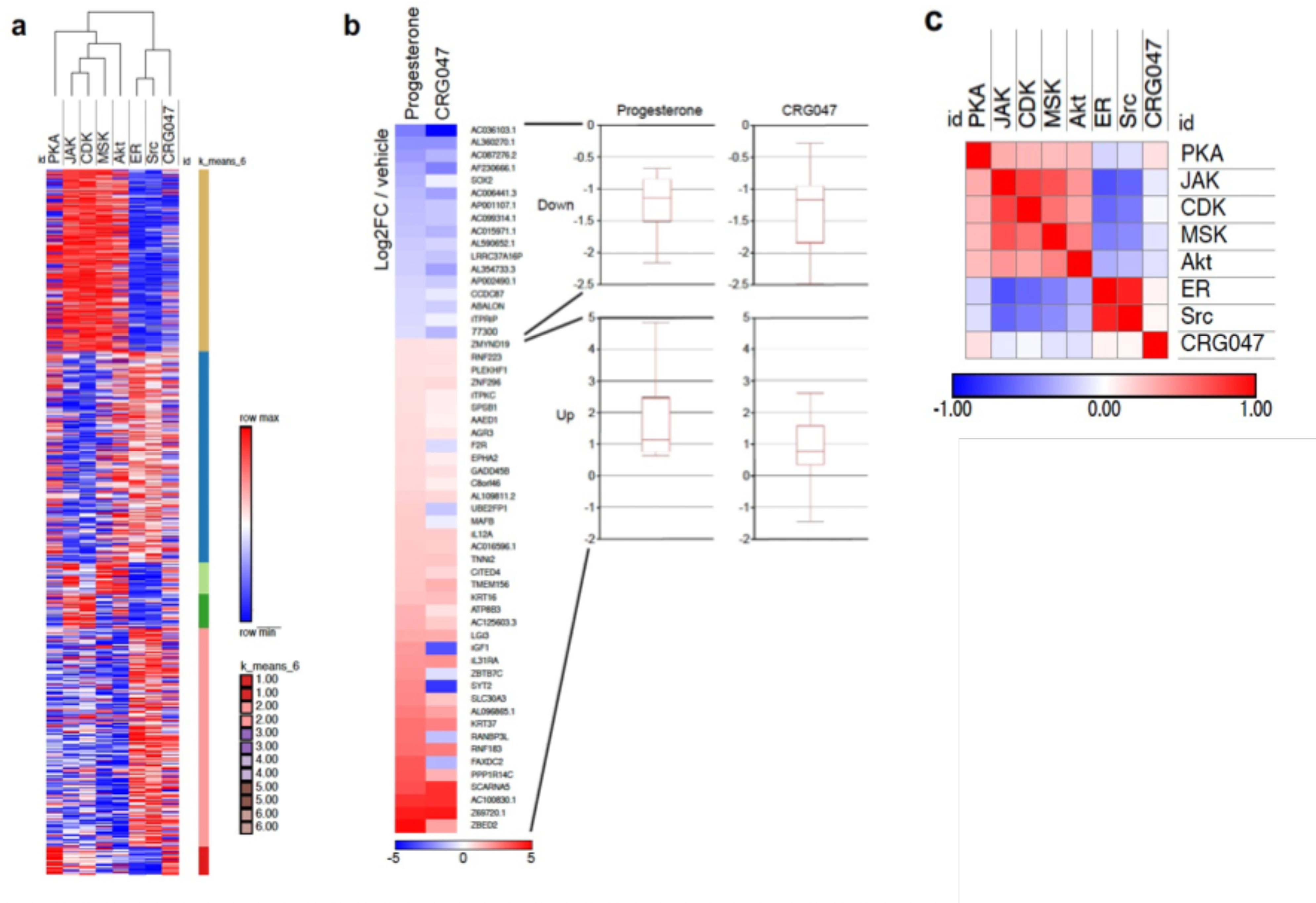
a) Hierarchical clustering of the gene expression signatures of R5020 regulated genes following inhibition of the protein indicated (PKA, JAK, CDK, MSK, Akt, ER or SRC) with the gene expression signature following CRG047 treatment. Analysis was carried out using Morpheus (Broad Institute); K means clustering on rows indicates 6 significantly distinct gene groups. Each row indicates a specific gene and the data was log2 normalized per row before clustering. b) Correlation matrix between significant genes from both treatments, clustered by genes down-regulated (red) and up-regulated (blue). Boxplots depict correlation of genes enclosed in each category, showing most of the genes although having a low FC over vehicle they fall on the same side of activation (positive or negative). c) Similarity matrix comparing the gene expression signatures of R5020 regulated genes following inhibition of the protein indicated (PKA, JAK, CDK, MSK, Akt, ER or SRC) with the gene expression signature following CRG047 treatment. Analysis was carried out using Morpheus (Broad Institute); negative Pearson correlations are shown in blue and positive in red (−1 to 1).

**Figure S5.**
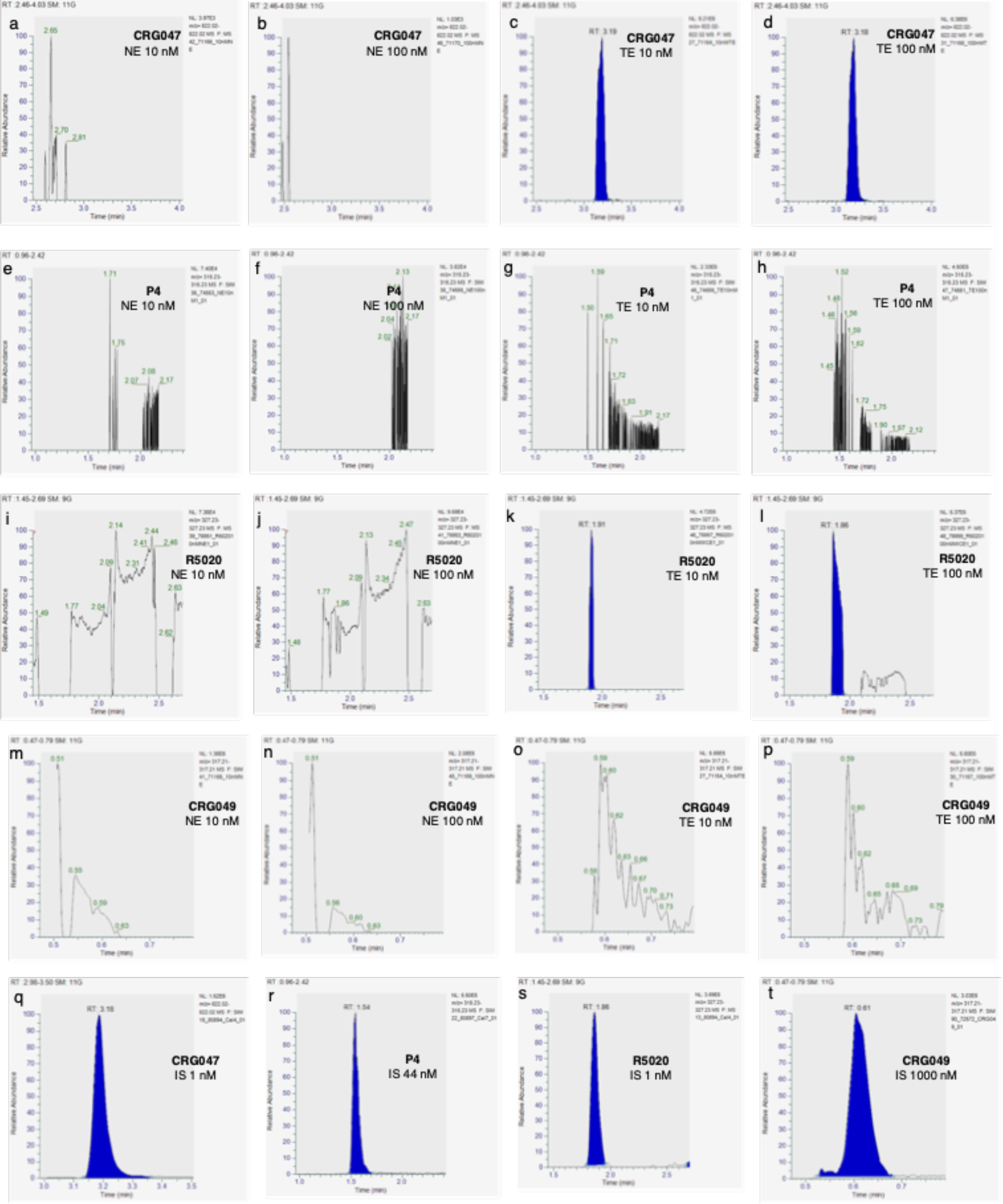
Mass spectrometry data on analyzed compounds. UHPLC-ESI-MS/MS SIM traces of precursor ions of CRG047 (a-d), P4 (e-h), R5020 (i-l) and CRG049 (m-p) compound for exposure to 10 nM and 100 nM as indicated in nuclear extracts (NE) and total extracts (TE). Panels q-t show the signals arising from internal standards (IS) of CRG047, P4, R5020 and CRG049 at indicated concentrations respectively.

**Figure S6.**
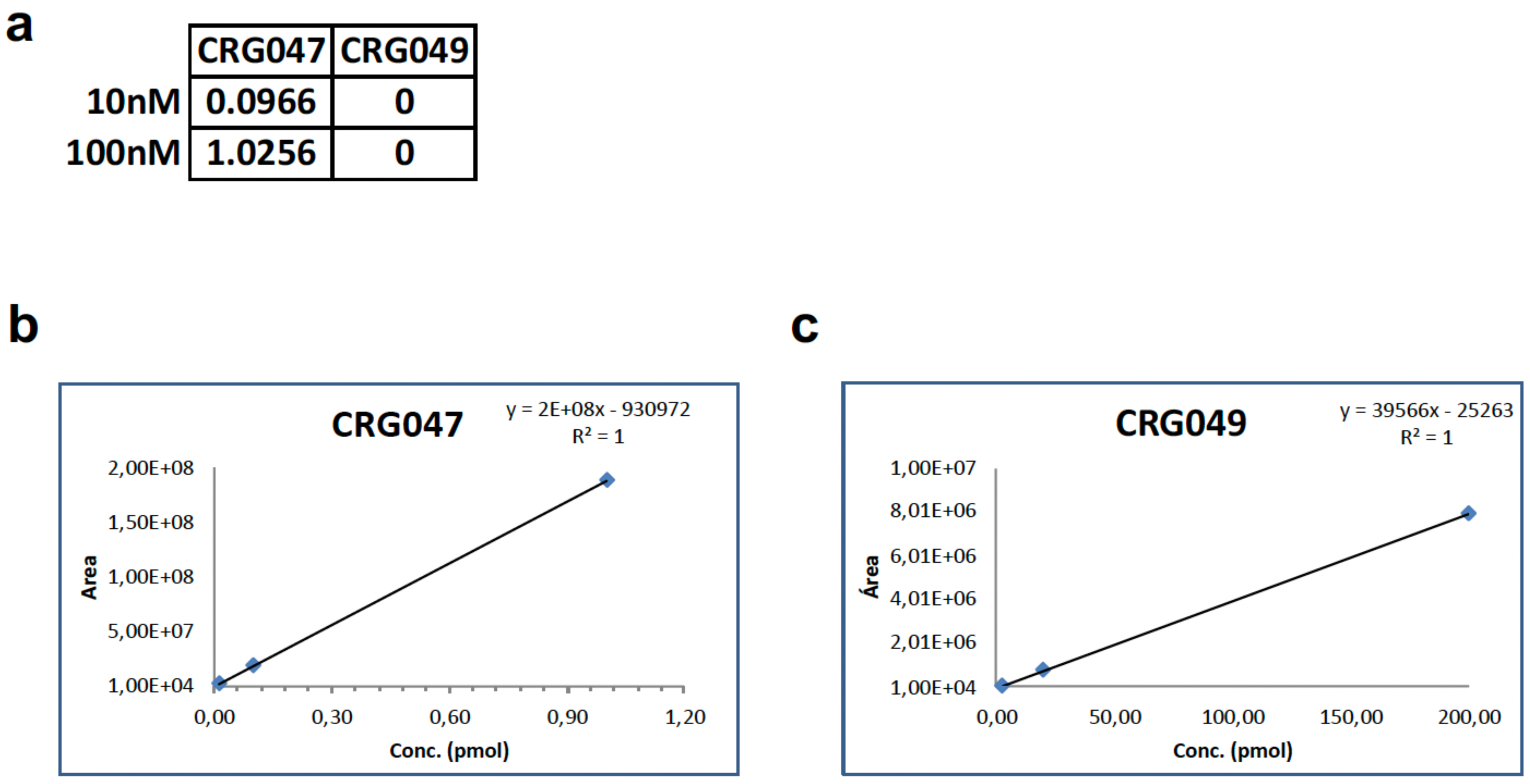
a) Table represents pmol/10^6^ cells of the indicated compounds detected (CRG047 or CRG049) following addition of CRG047 at the concentrations indicated. b-c) Calibration curves of indicated compounds for mass spectrometry analysis derived from internal standards for CRG047 and CRG049 as indicated.

**Figure S7; Figure 6 - Figure supplement 1.**
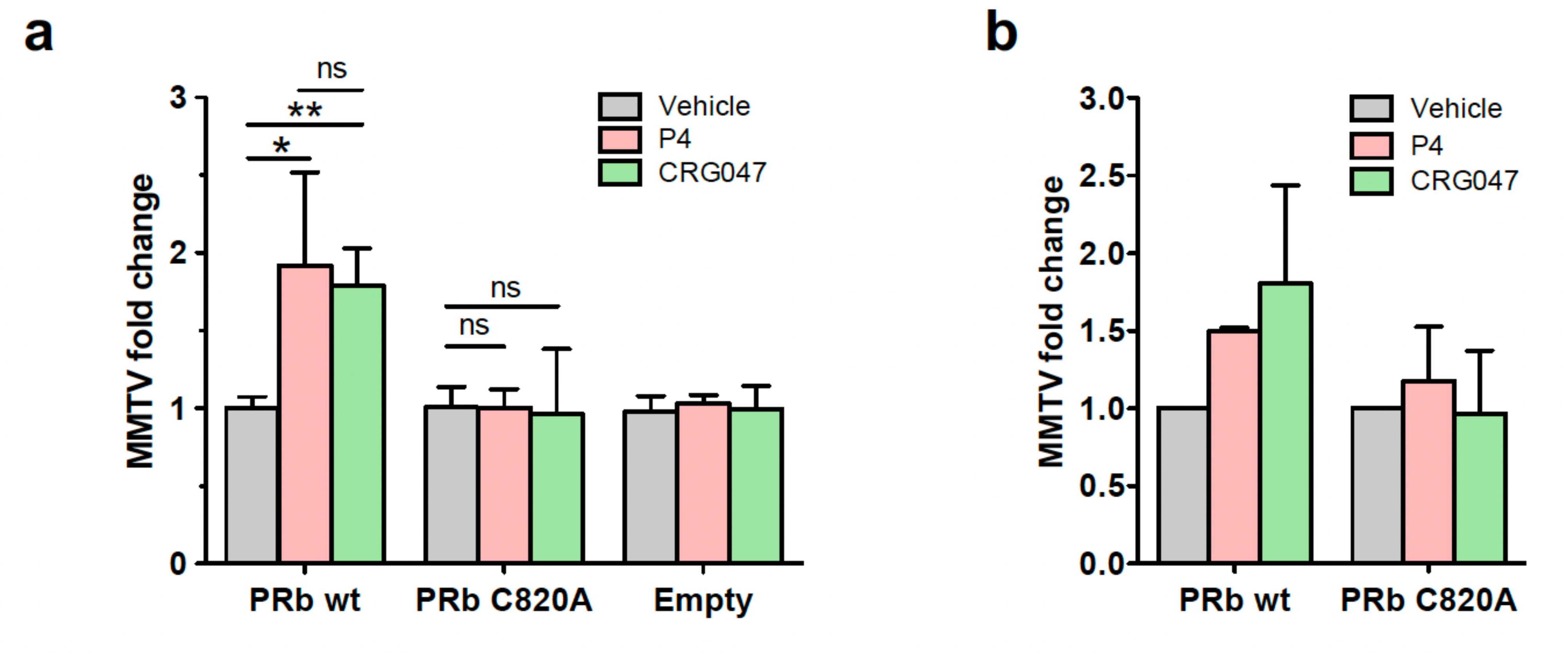
a) PR-negative breast cancer cell line T47D-Y stably expressing a single copy of the MMTV promoter was transfected with plasmids for the expression of either wild type PRb, palmitoylation mutant PRbC820A or with empty plasmids and incubated with either vehicle, progesterone (P4) or CRG047 as indicated. RNA signal for MMTV is shown. Average ± SD of 3 independent experiments is shown. Fold changes are relative to the average response of vehicle-incubated cells. *: p-value < 0.05; **: p-value < 0.01; ns: p-value > 0.05. Statistical analysis was done by 1 tail T-test comparing the indicated conditions. b) U2OS cells were co-transfected with plasmids expressing a MMTV-luc reporter and either wild type PRb or palmitoylation mutant PRbC820A and incubated with either vehicle, progesterone (P4) or CRG047 and qPCR was done against MMTV. Histograms show average ± SD of 2 independent experiments.

**Figure S8.**
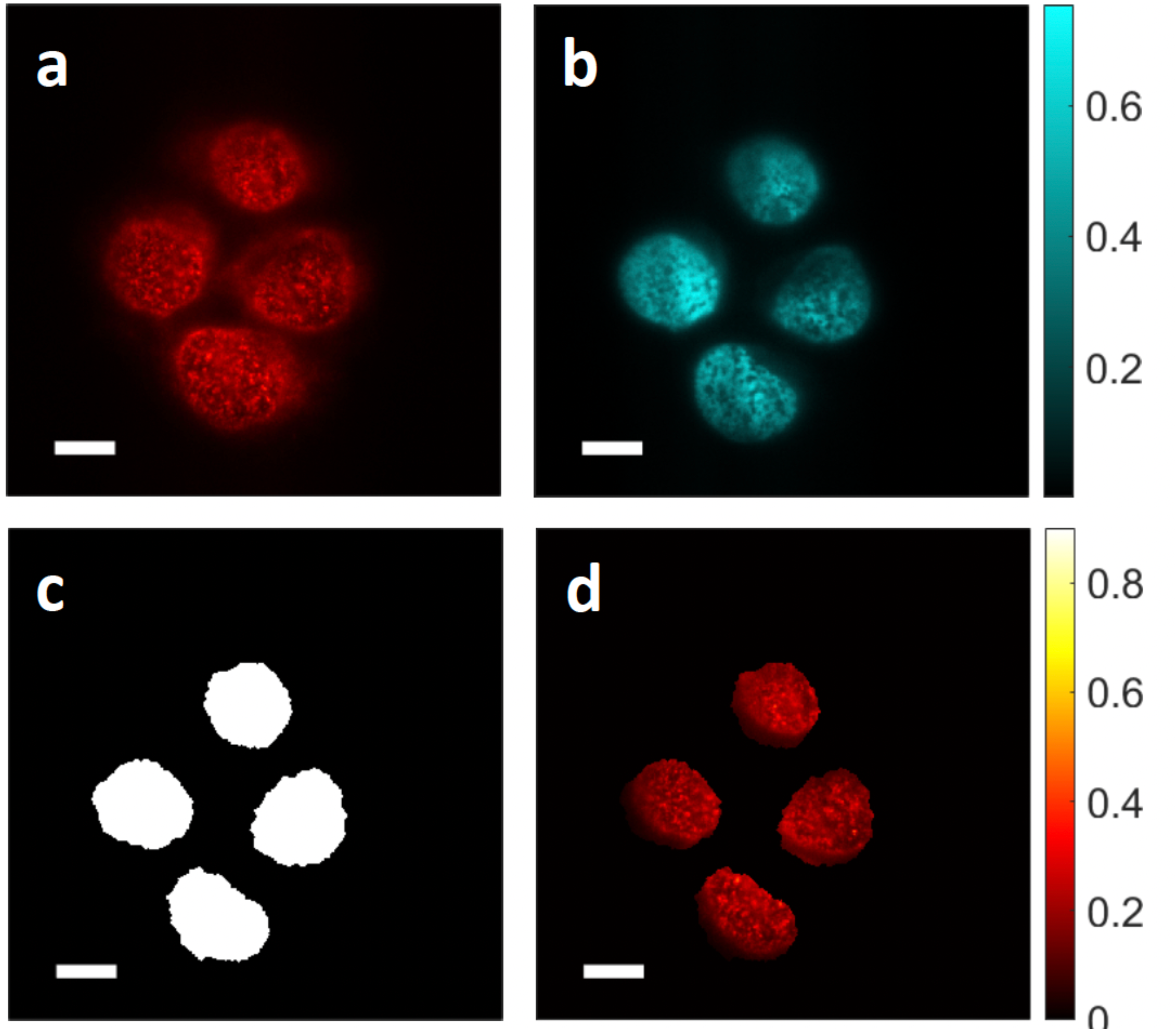
Procedure for the quantification of immunofluorescence. a) PR Ser 294 phosphorylation (S294P) immunofluorescent image. b) DAPI image. c) Masks for each nucleus obtained from the DAPI image (b). d) The result of applying the mask (c) to the S294P image (a). Images (a-d) have been normalized to the maximum intensity value for better visualization. Scale bar for all panels: 10 μm.

### NMR spectra of the synthetized compounds.

**Figure S9.**
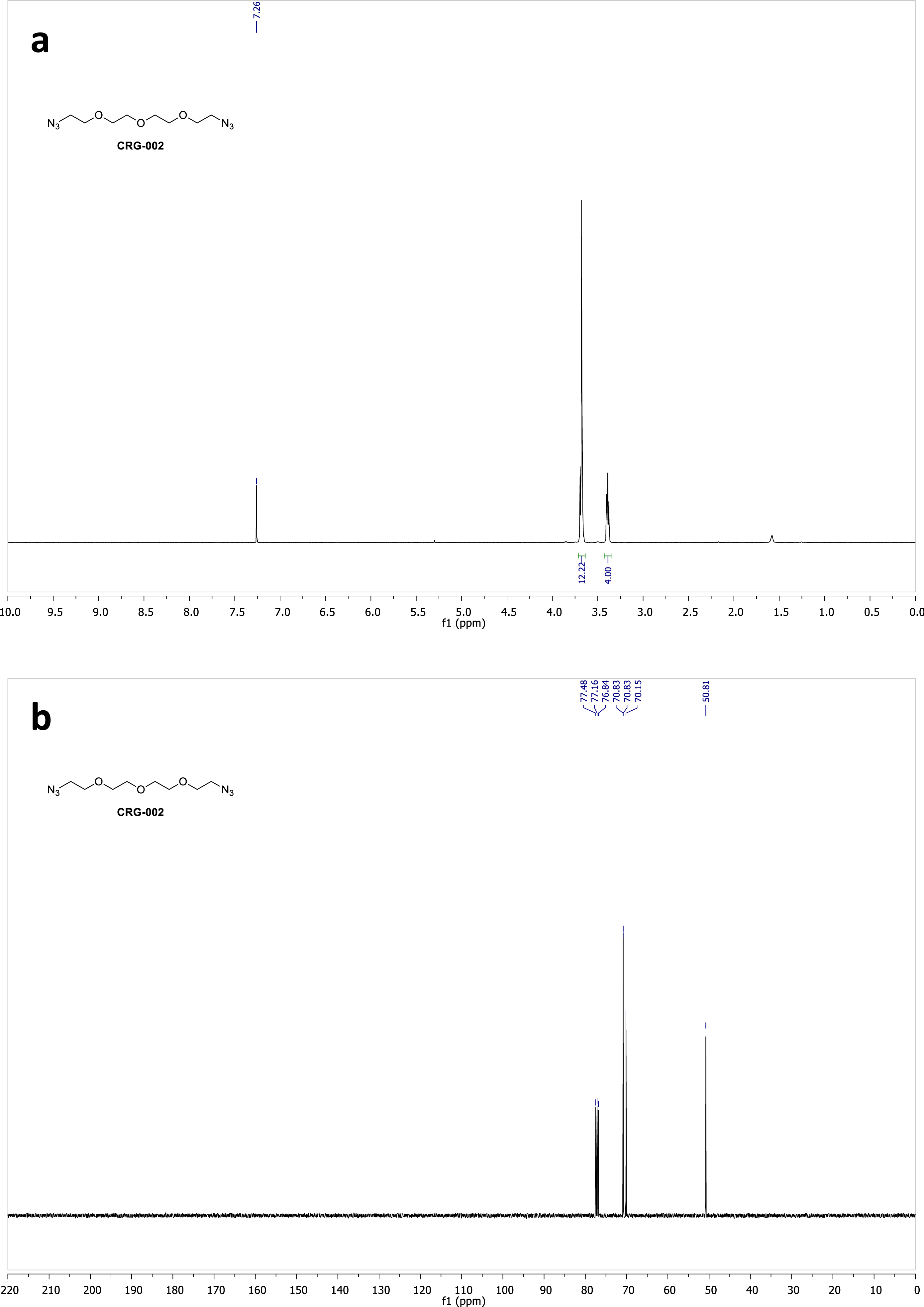
^1^H (a) and ^13^C (b) NMR spectra of compound **CRG002** in CDCl_3_.

**Figure S10.**
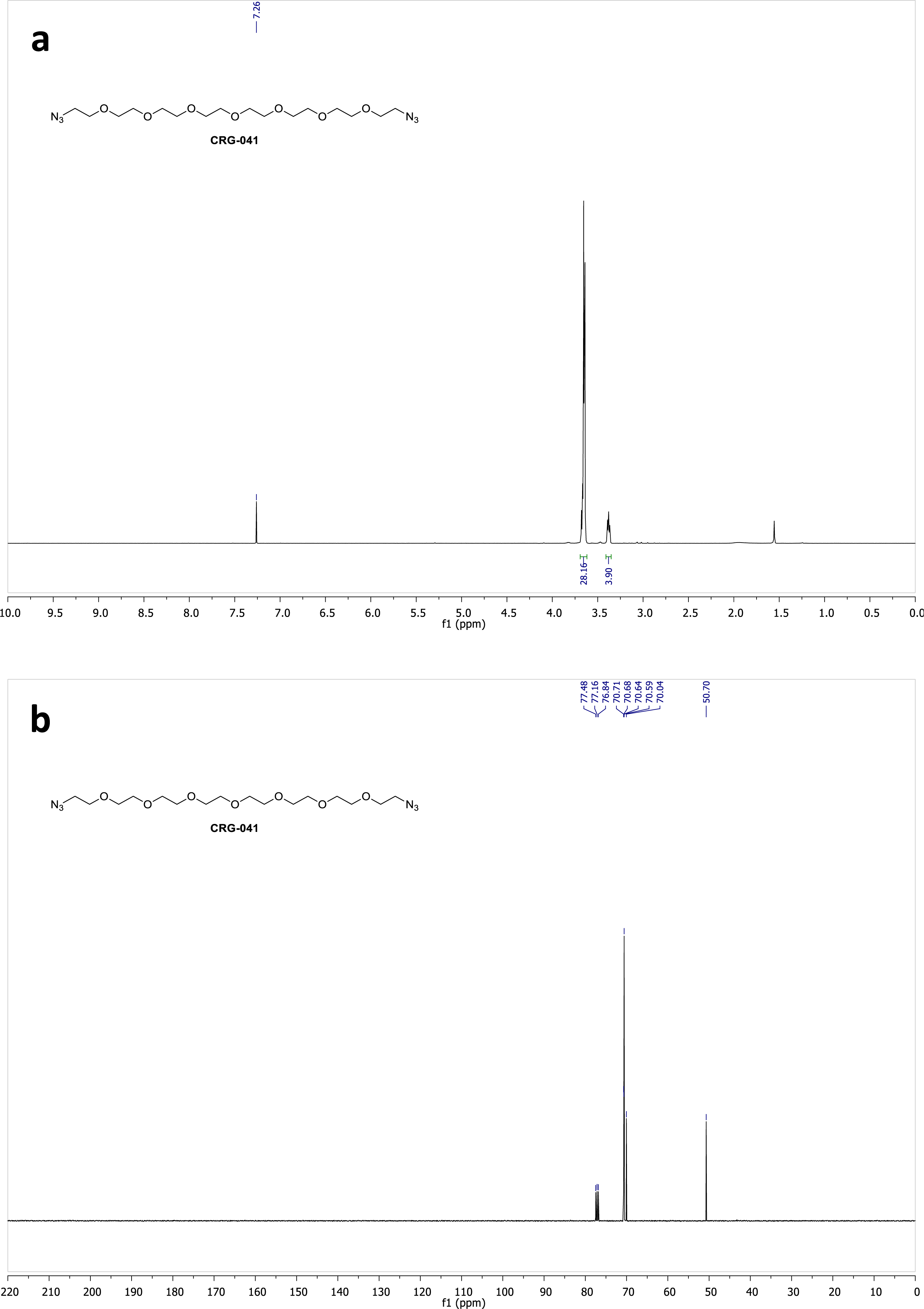
^1^H (a) and ^13^C (b) NMR spectra of compound **CRG041** in CDCl_3_.

**Figure S11.**
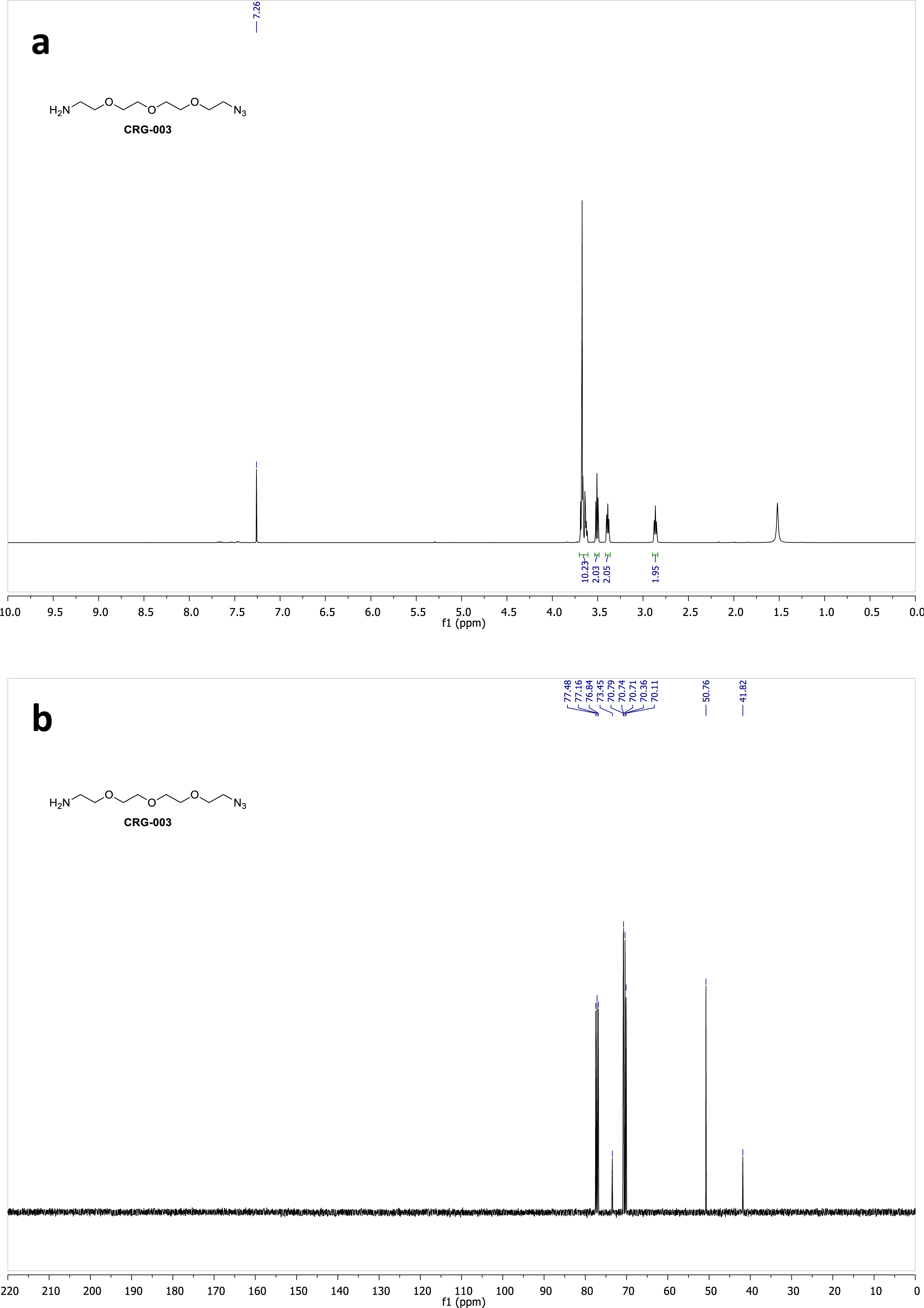
^1^H (a) and ^13^C (b) NMR spectra of compound **CRG003** in CDCl_3_.

**Figure S12.**
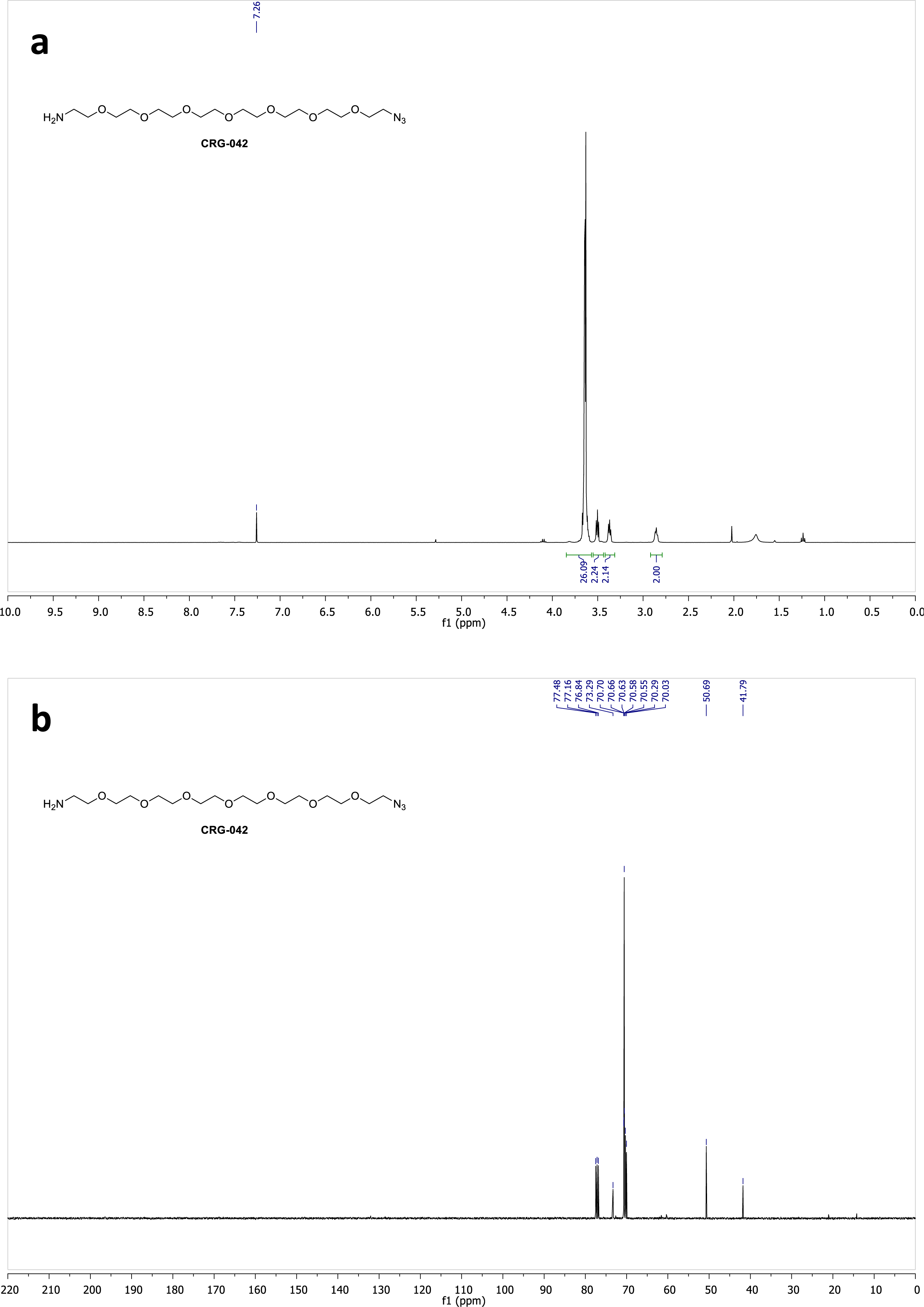
^1^H (a) and ^13^C (b) NMR spectra of compound **CRG042** in CDCl_3_.

**Figure S13.**
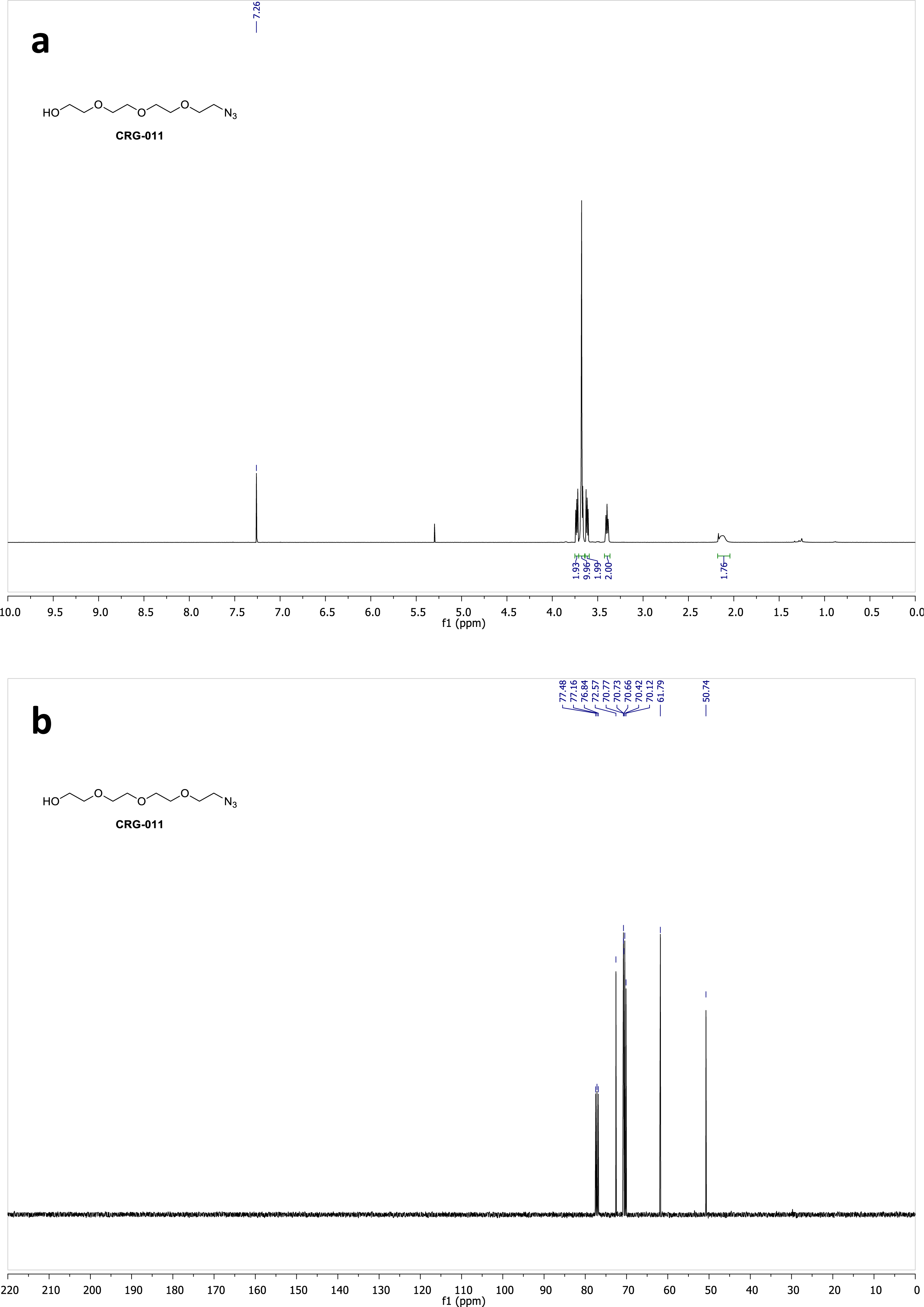
^1^H (a) and ^13^C (b) NMR spectra of compound **CRG011** in CDCl_3_.

**Figure S14.**
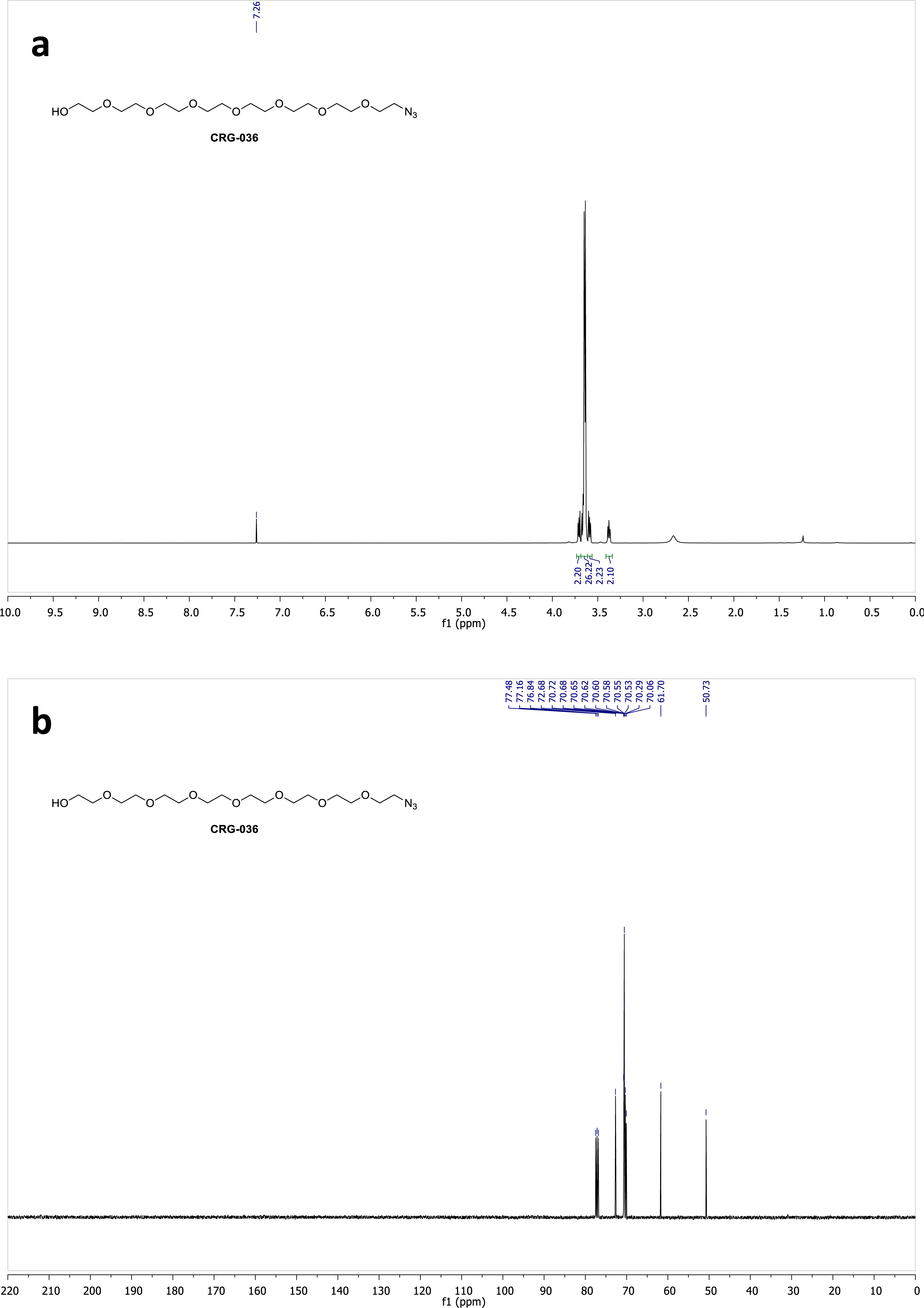
^1^H (a) and ^13^C (b) NMR spectra of compound **CRG036** in CDCl_3_.

**Figure S15.**
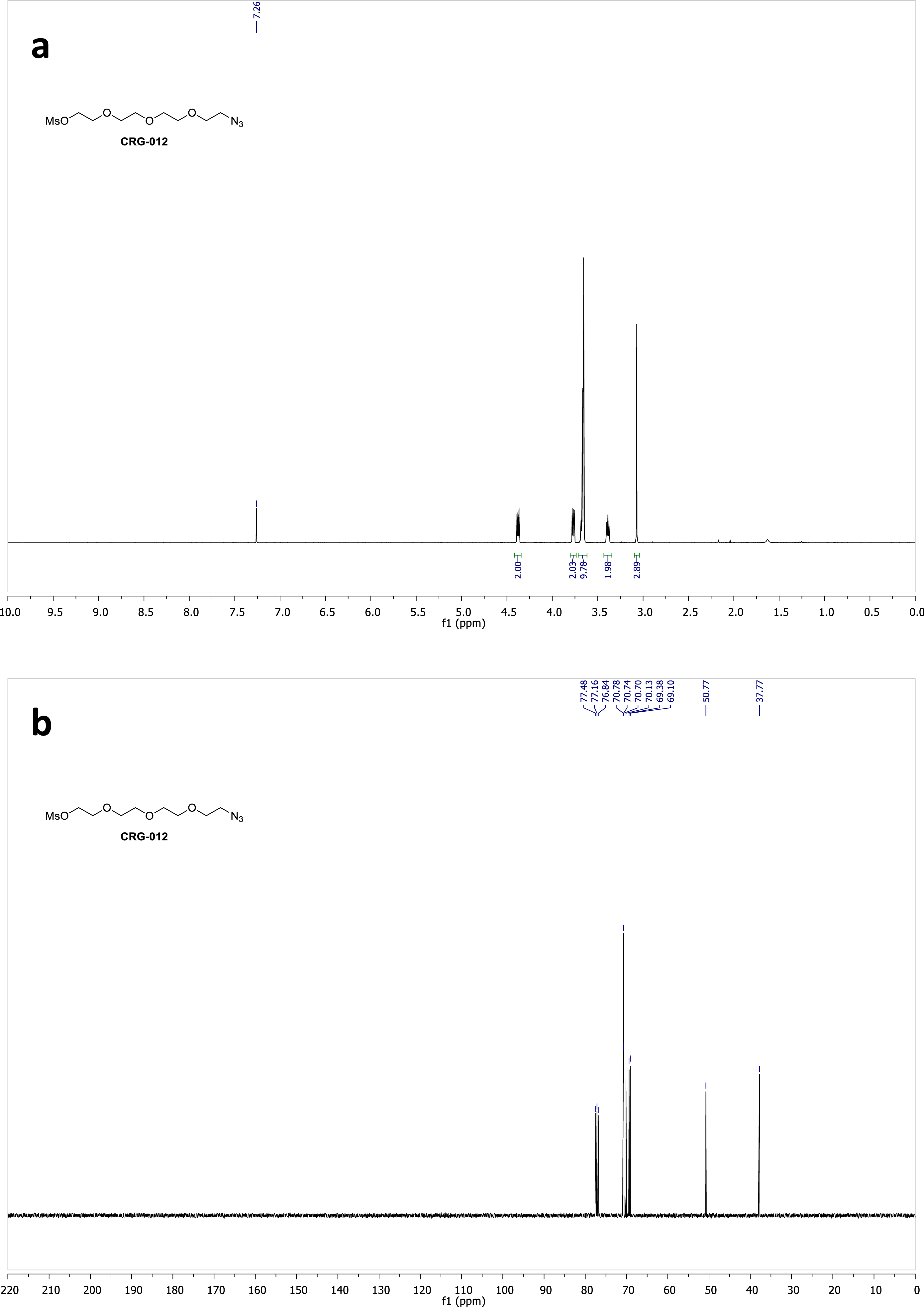
^1^H (a) and ^13^C (b) NMR spectra of compound **CRG012** in CDCl_3_.

**Figure S16.**
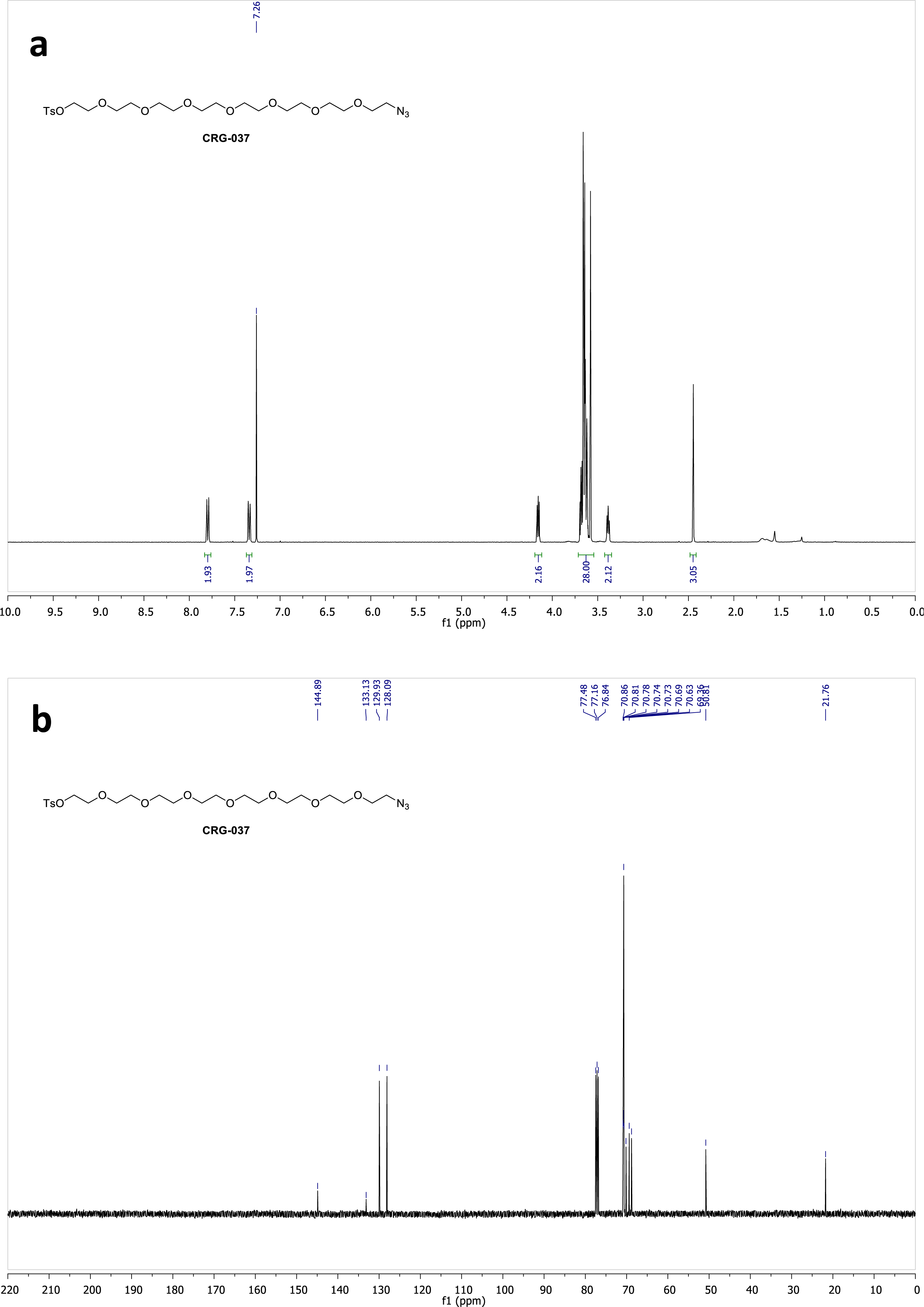
^1^H (a) and ^13^C (b) NMR spectra of compound **CRG037** in CDCl_3_.

**Figure S17.**
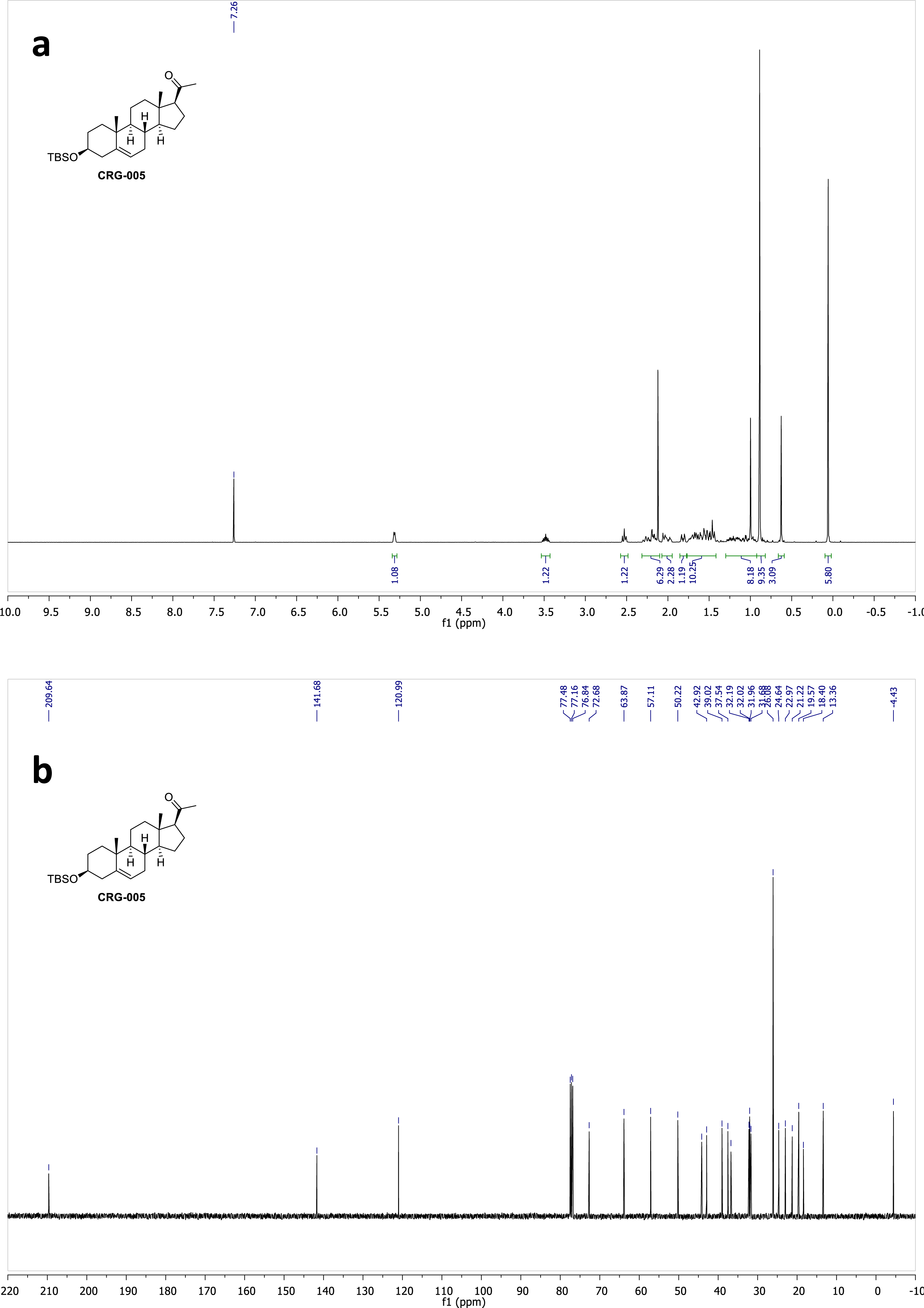
^1^H (a) and ^13^C (b) NMR spectra of compound **CRG005** in CDCl_3_.

**Figure S18.**
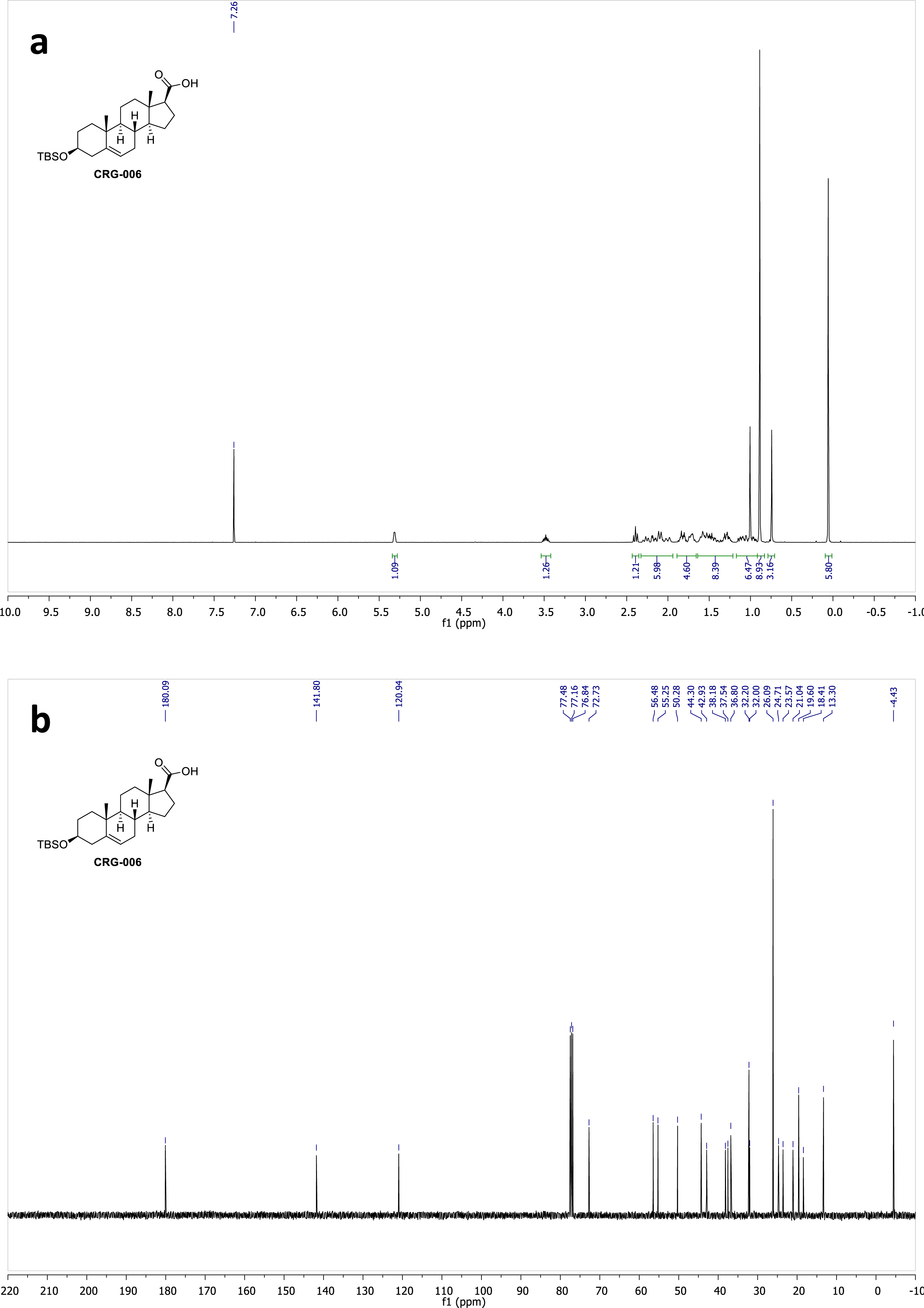
^1^H (a) and ^13^C (b) NMR spectra of compound **CRG006** in CDCl_3_.

**Figure S19.**
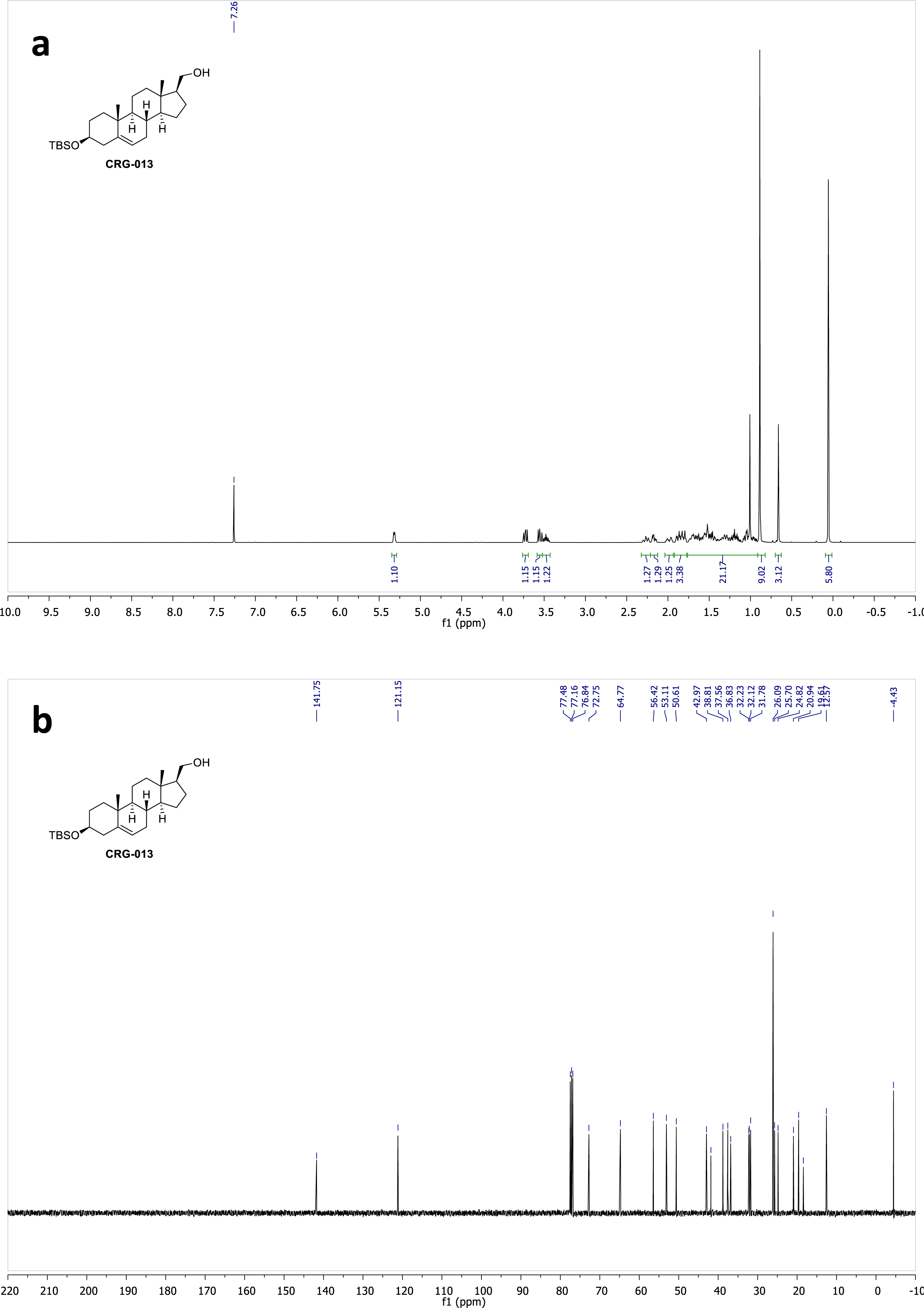
^1^H (a) and ^13^C (b) NMR spectra of compound **CRG013** in CDCl_3_.

**Figure S20.**
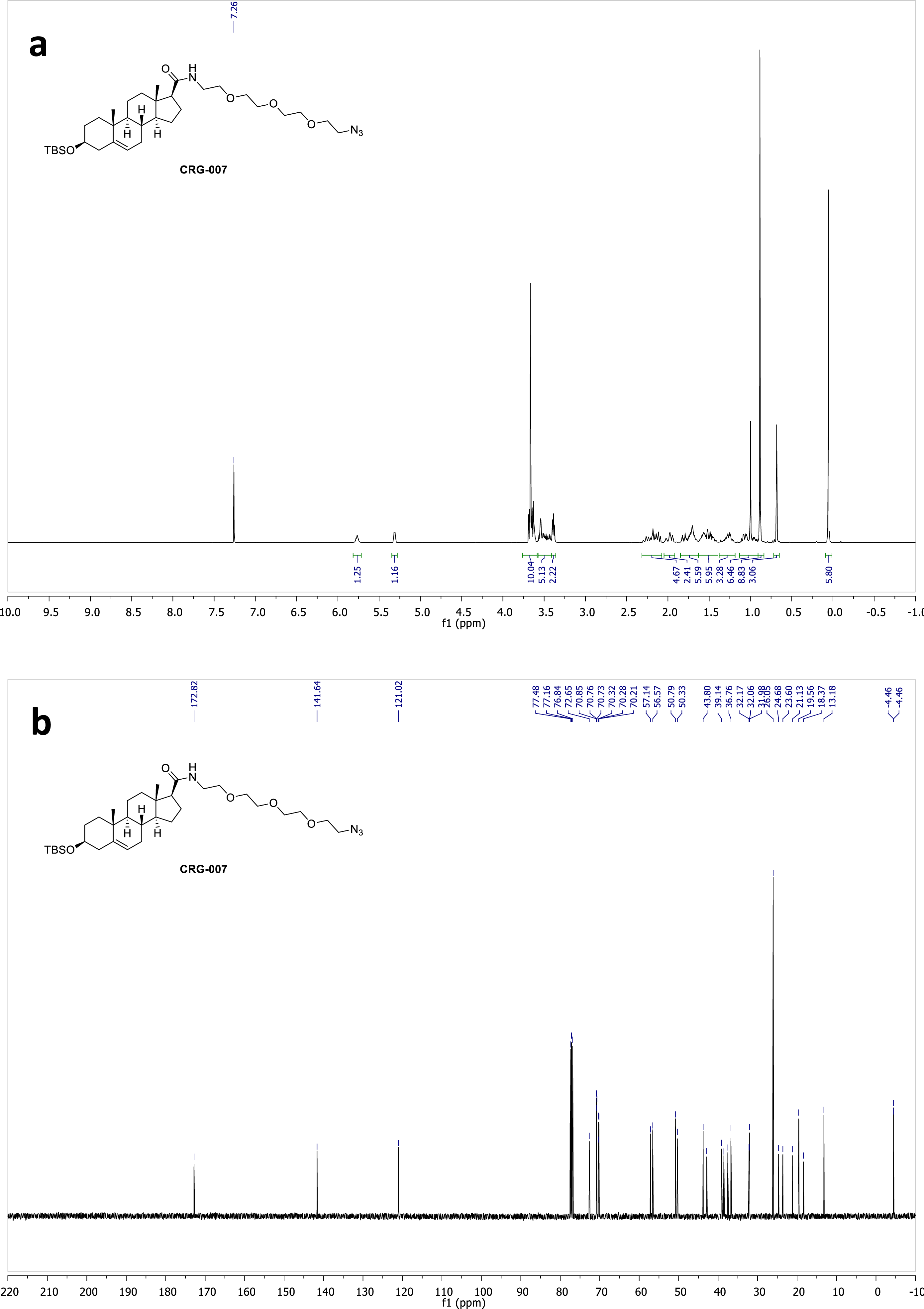
^1^H (a) and ^13^C (b) NMR spectra of compound **CRG007** in CDCl_3_.

**Figure S21.**
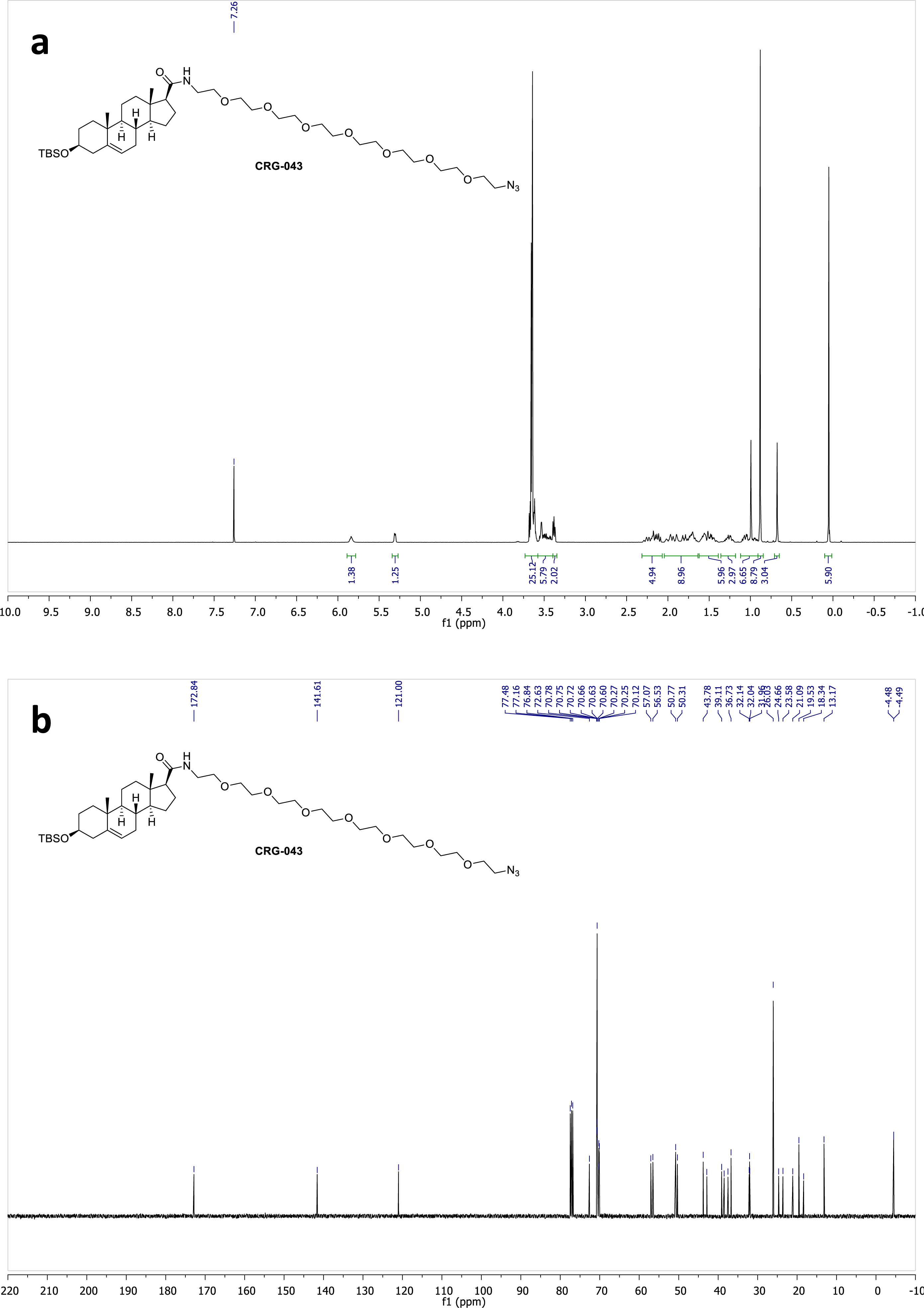
^1^H (a) and ^13^C (b) NMR spectra of compound **CRG043** in CDCl_3_.

**Figure S22.**
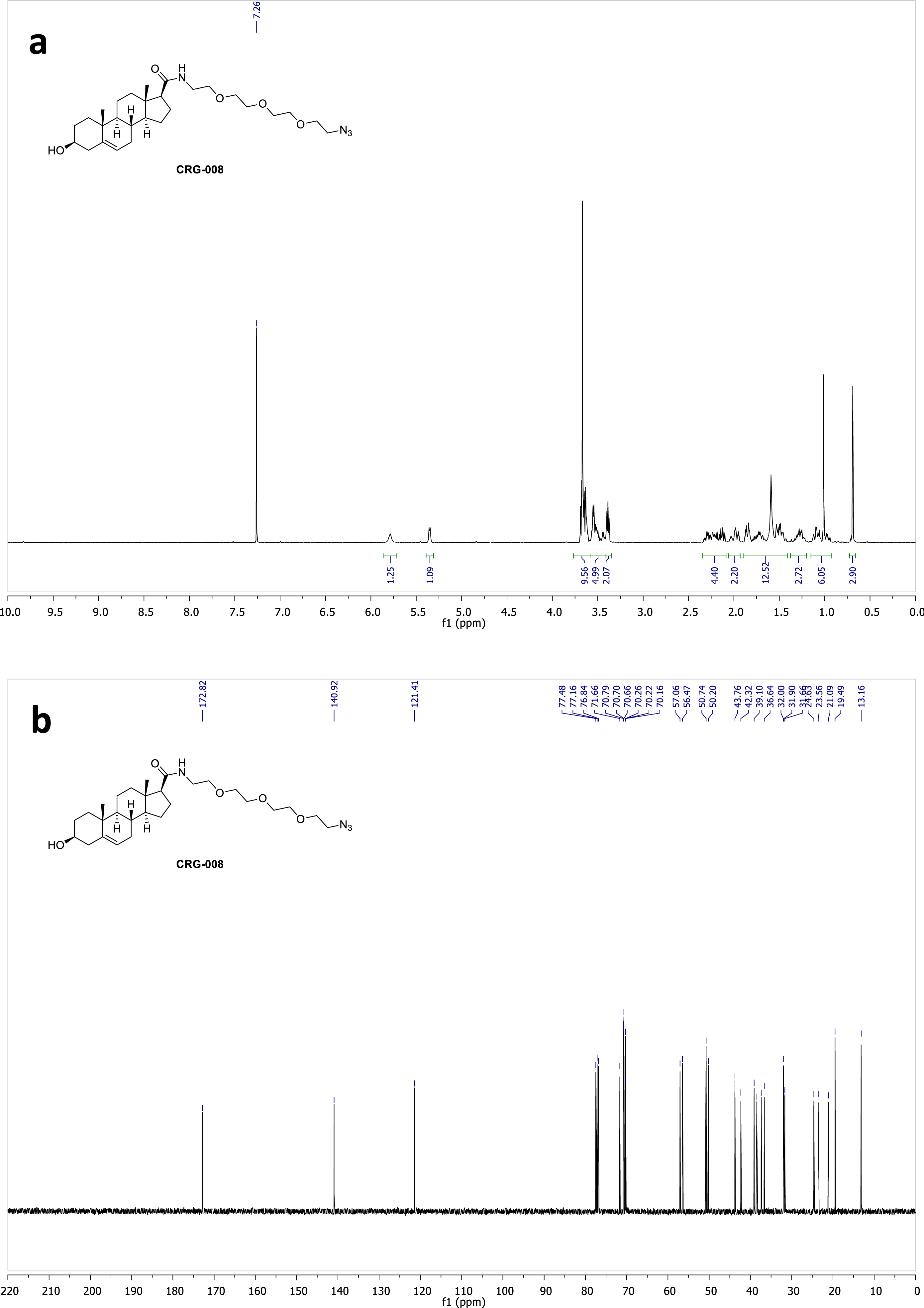
^1^H (a) and ^13^C (b) NMR spectra of compound **CRG008** in CDCl_3_.

**Figure S23.**
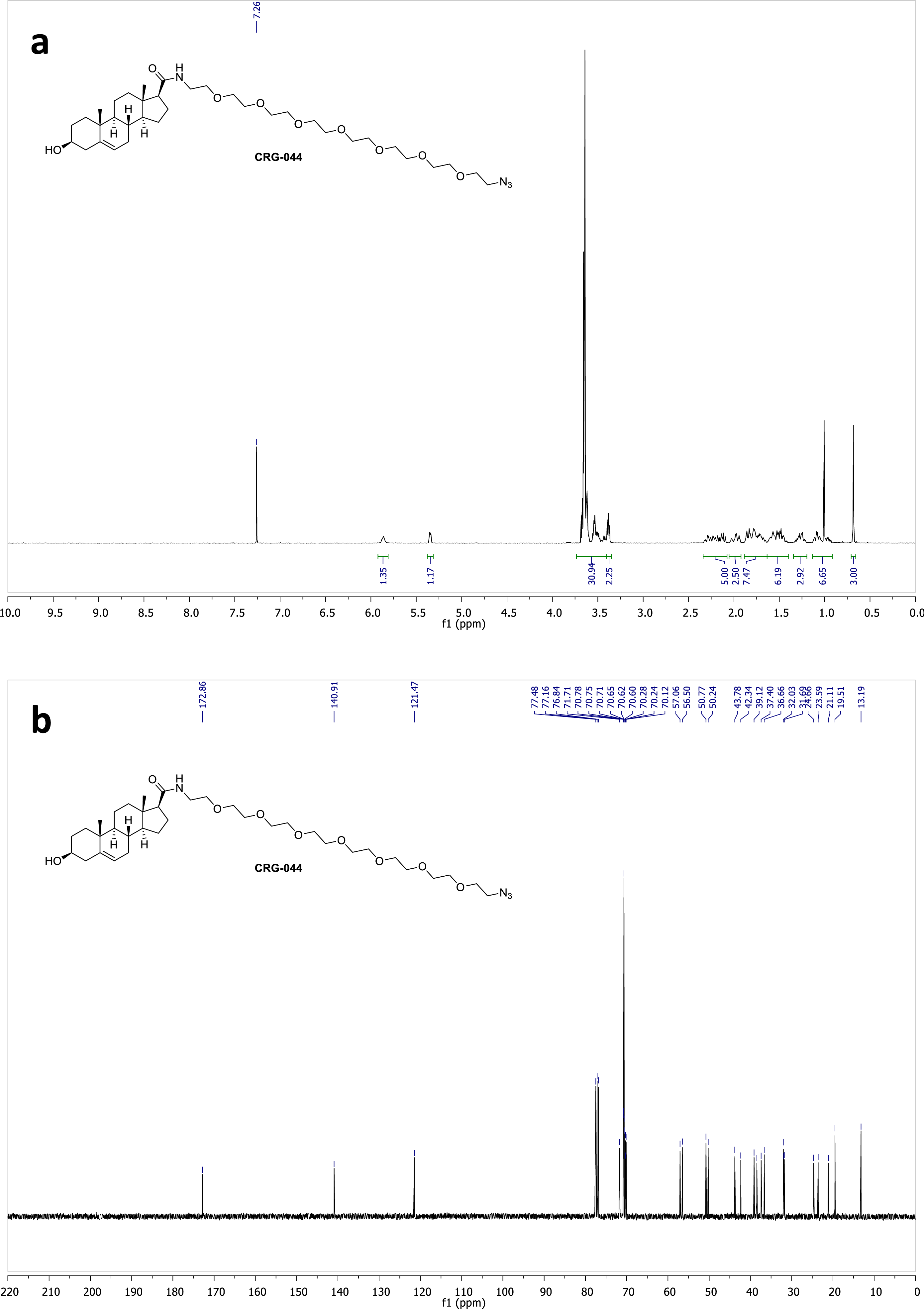
^1^H (a) and ^13^C (b) NMR spectra of compound **CRG044** in CDCl_3_.

**Figure S24.**
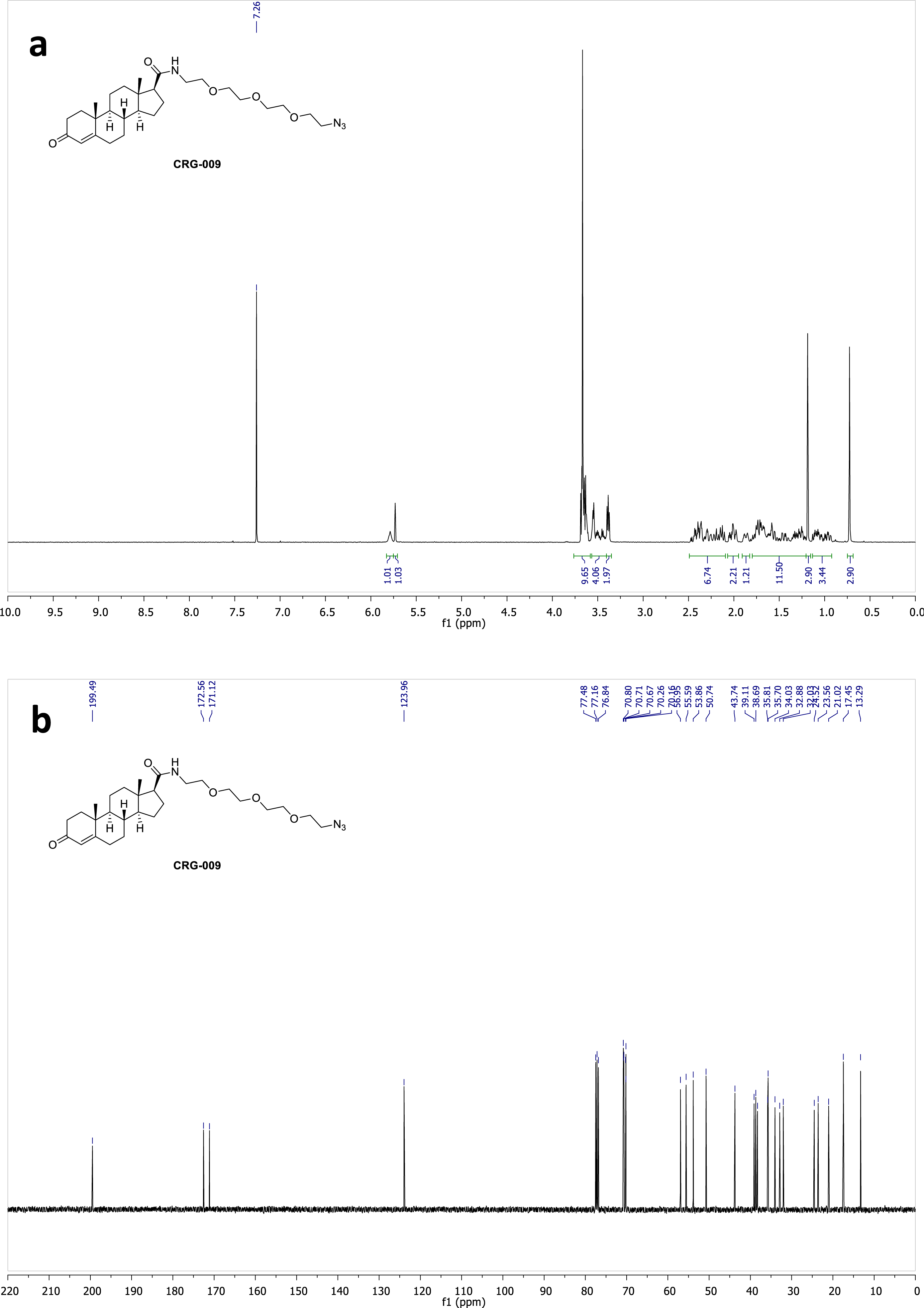
^1^H (a) and ^13^C (b) NMR spectra of compound **CRG009** in CDCl_3_.

**Figure S25.**
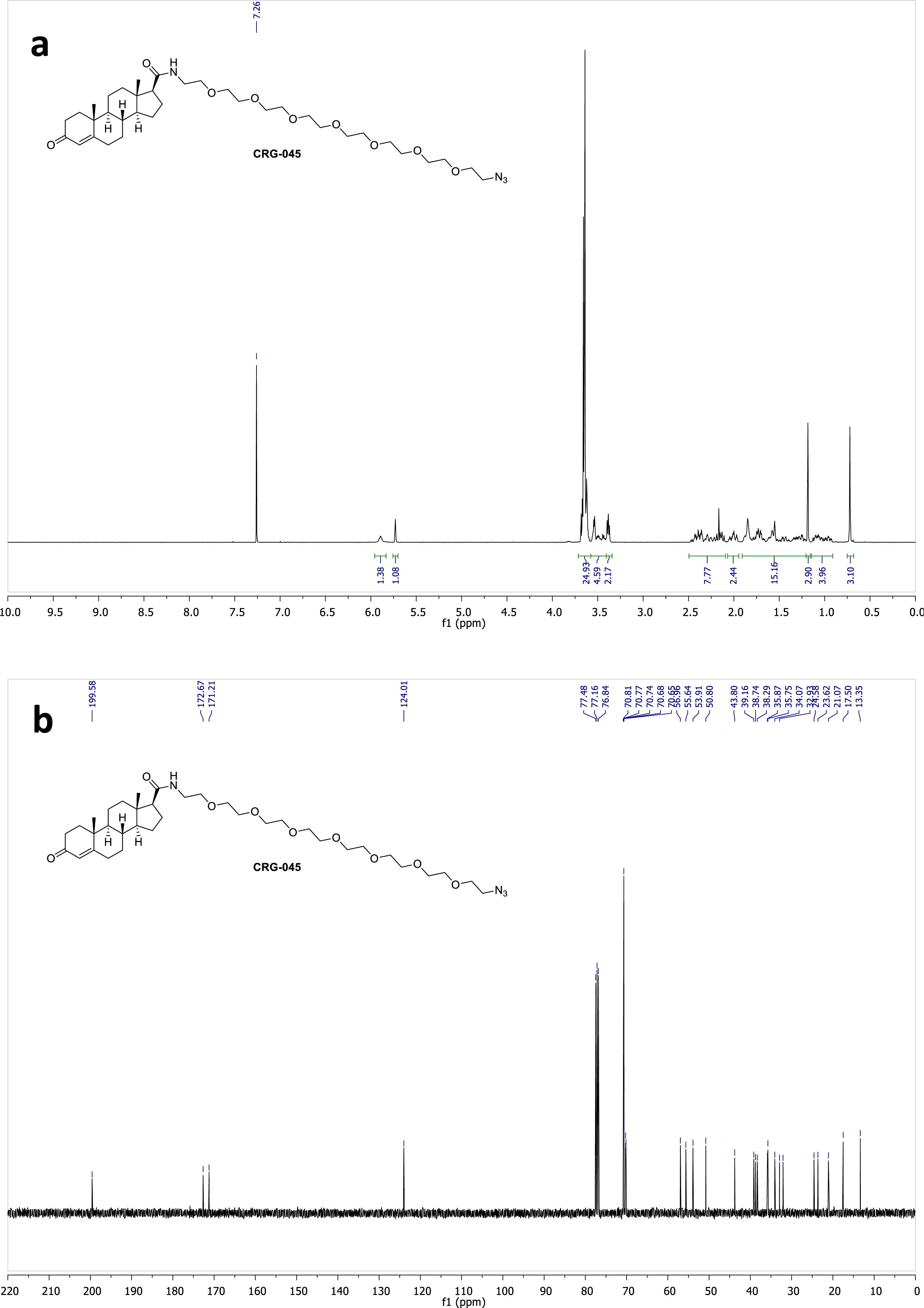
^1^H (a) and ^13^C (b) NMR spectra of compound **CRG045** in CDCl_3_.

**Figure S26.**
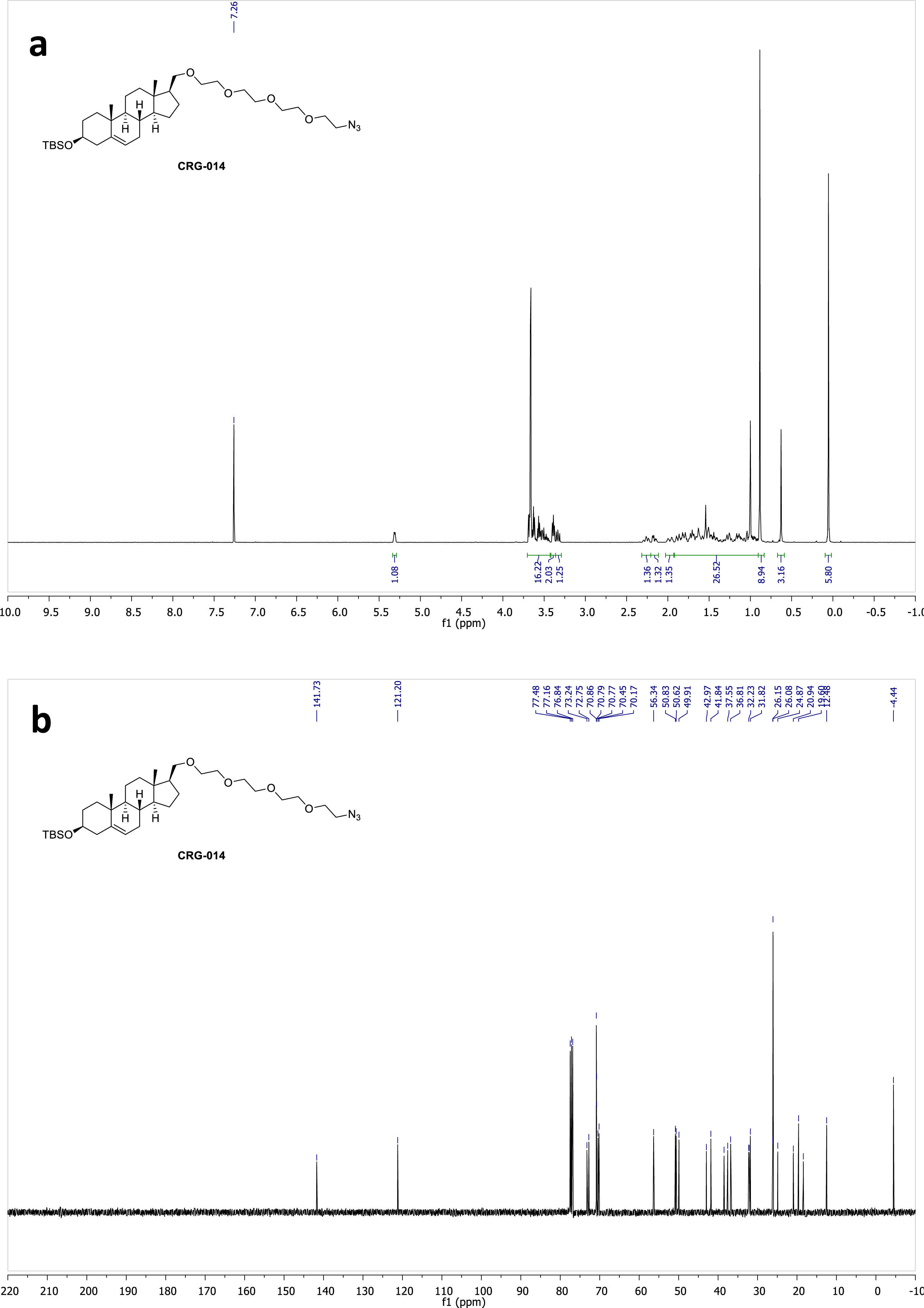
^1^H (a) and ^13^C (b) NMR spectra of compound **CRG014** in CDCl_3_.

**Figure S27.**
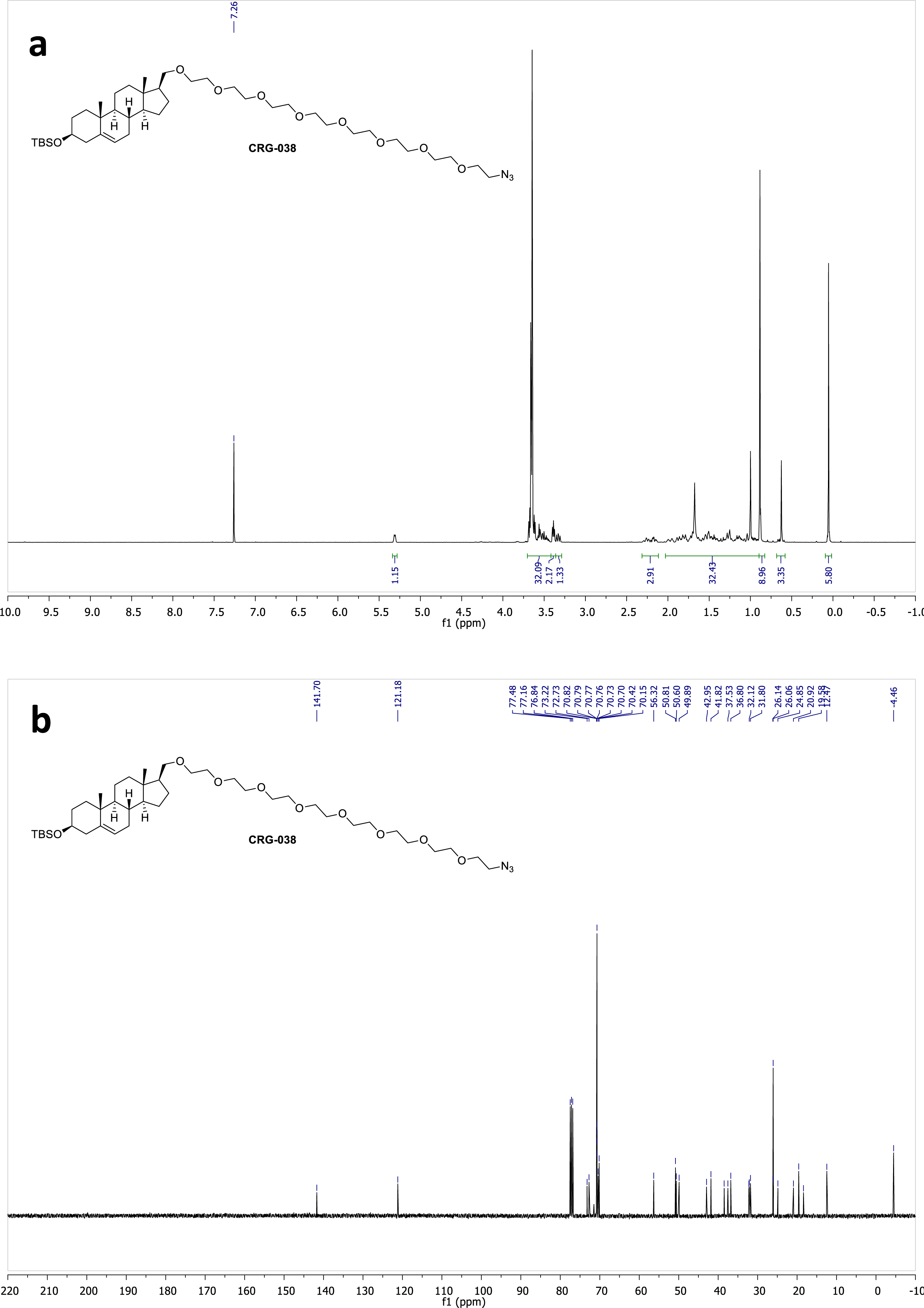
^1^H (a) and ^13^C (b) NMR spectra of compound **CRG038** in CDCl_3_.

**Figure S28.**
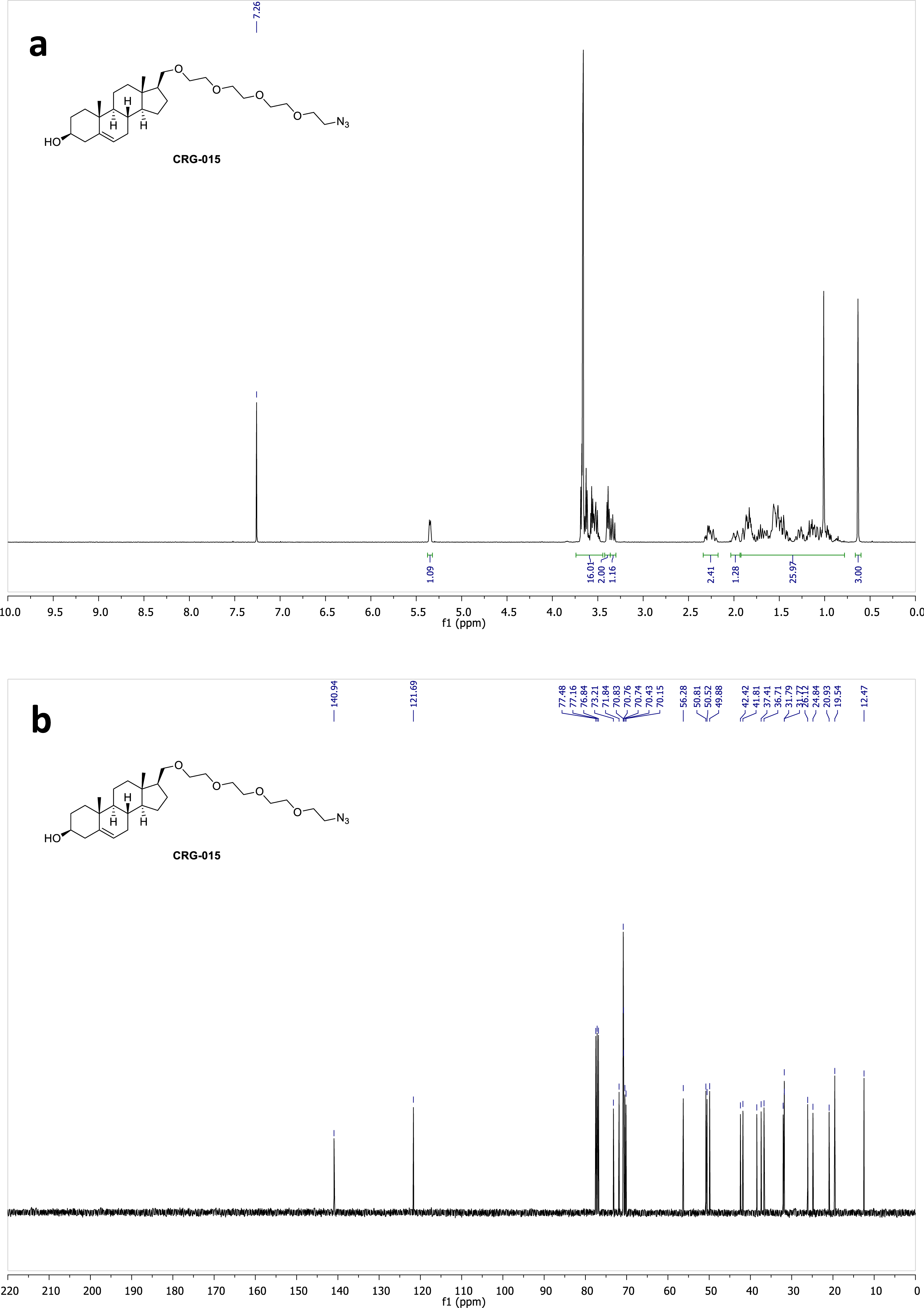
^1^H (a) and ^13^C (b) NMR spectra of compound **CRG015** in CDCl_3_.

**Figure S29.**
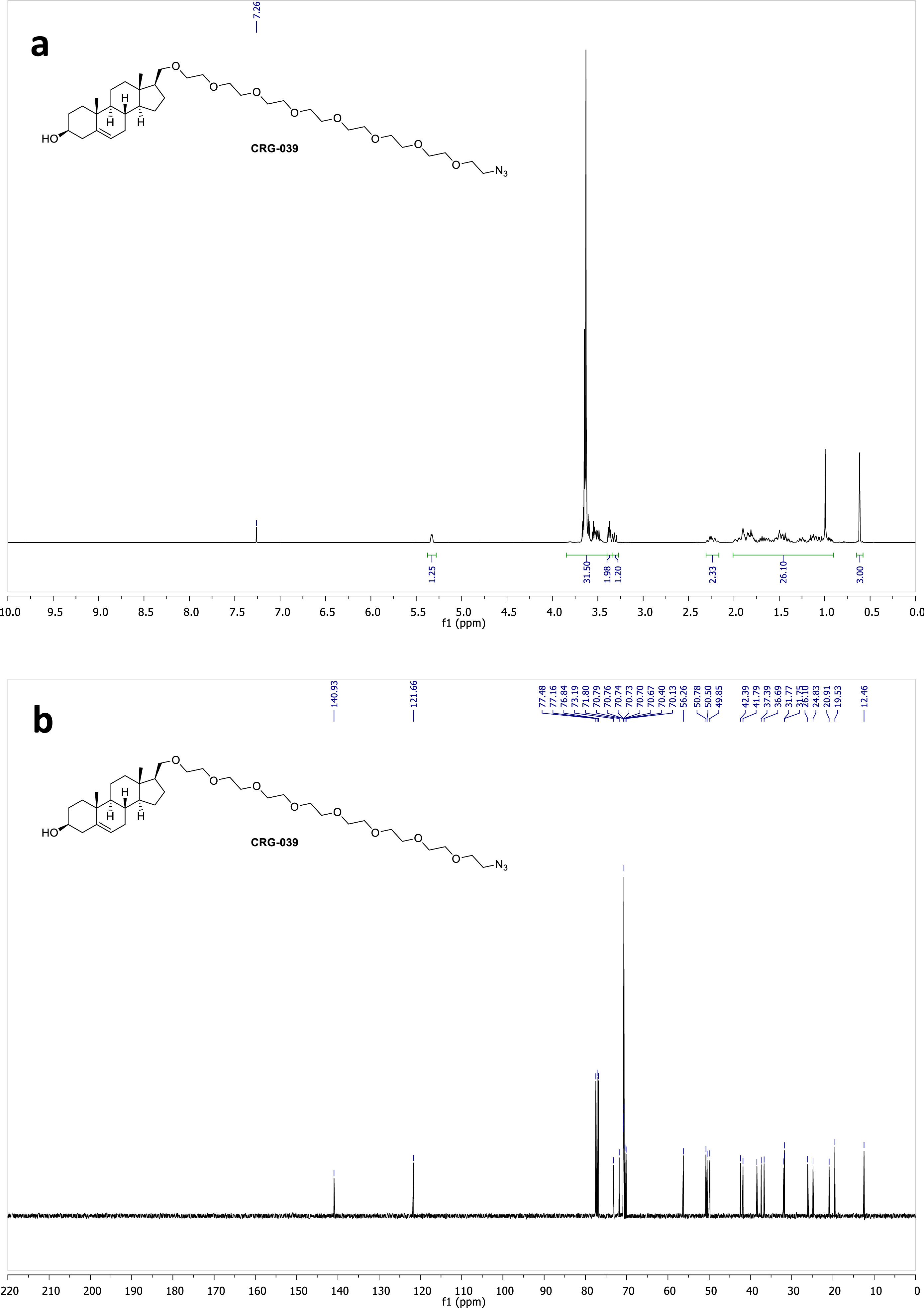
^1^H (a) and ^13^C (b) NMR spectra of compound **CRG039** in CDCl_3_.

**Figure S30.**
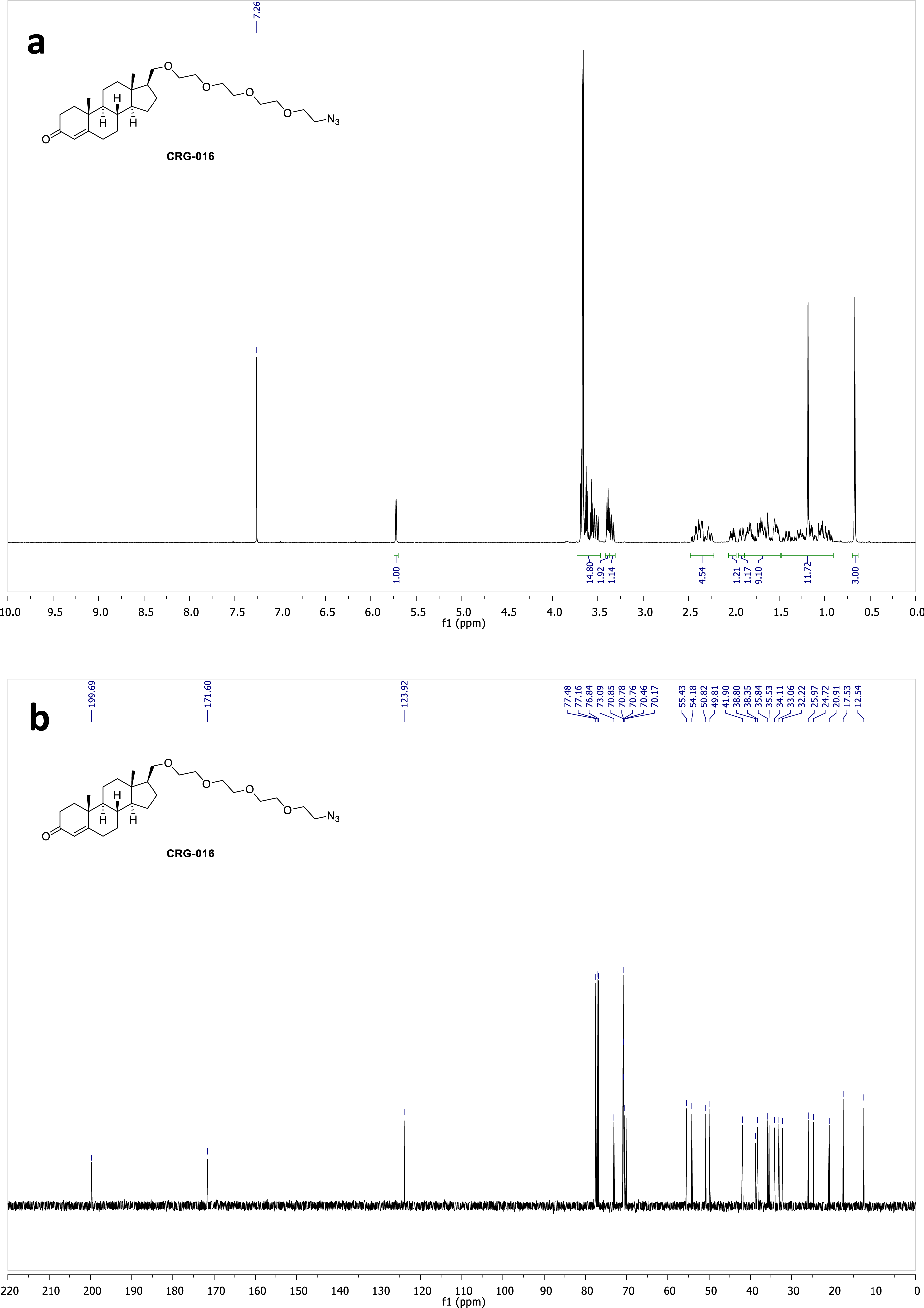
^1^H (a) and ^13^C (b) NMR spectra of compound **CRG016** in CDCl_3_.

**Figure S31.**
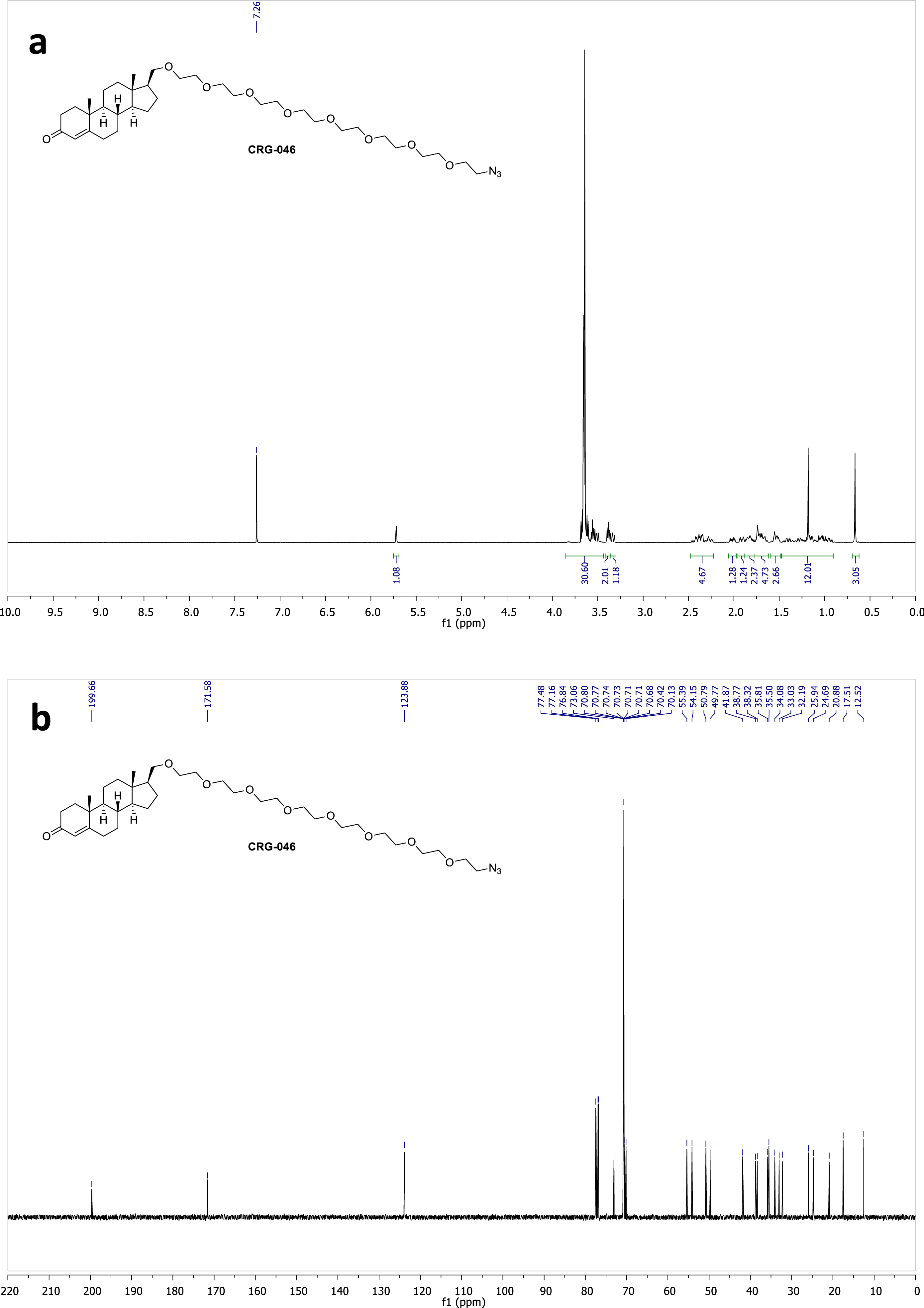
^1^H (a) and ^13^C (b) NMR spectra of compound **CRG046** in CDCl_3_.

**Figure S32.**
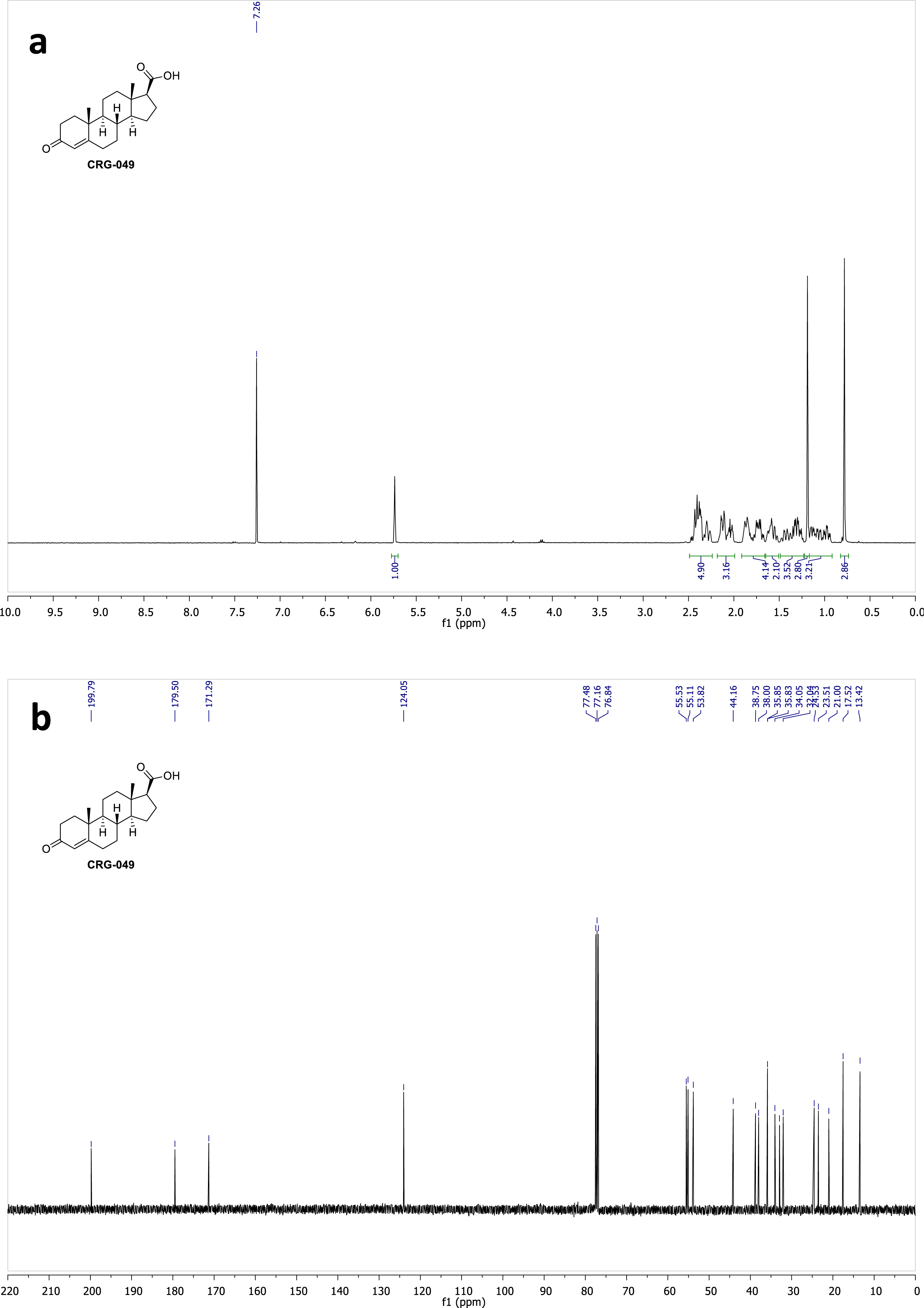
^1^H (a) and ^13^C (b) NMR spectra of compound **CRG049** in CDCl_3_.

**Figure S33.**
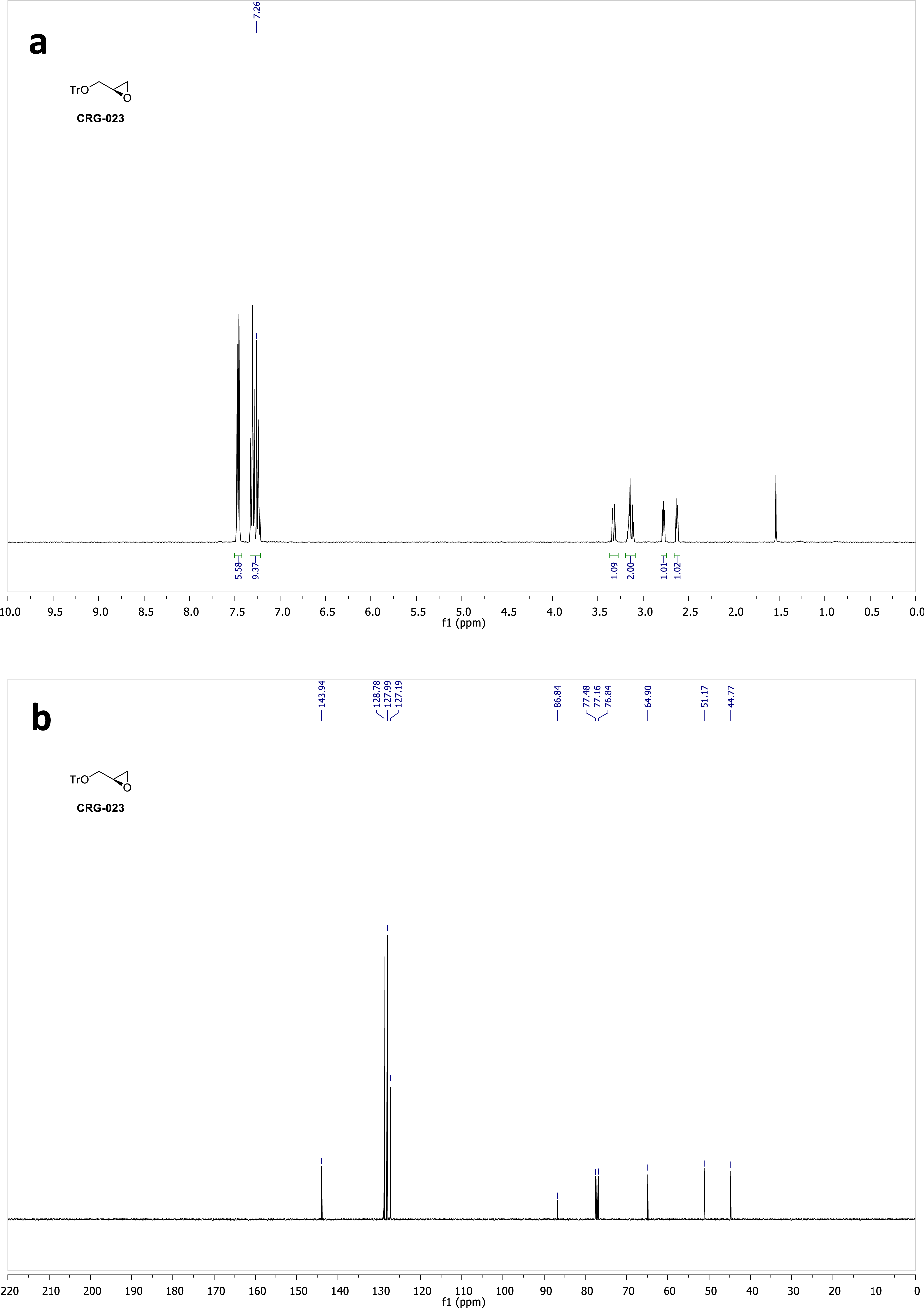
^1^H (a) and ^13^C (b) NMR spectra of compound **CRG023** in CDCl_3_.

**Figure S34.**
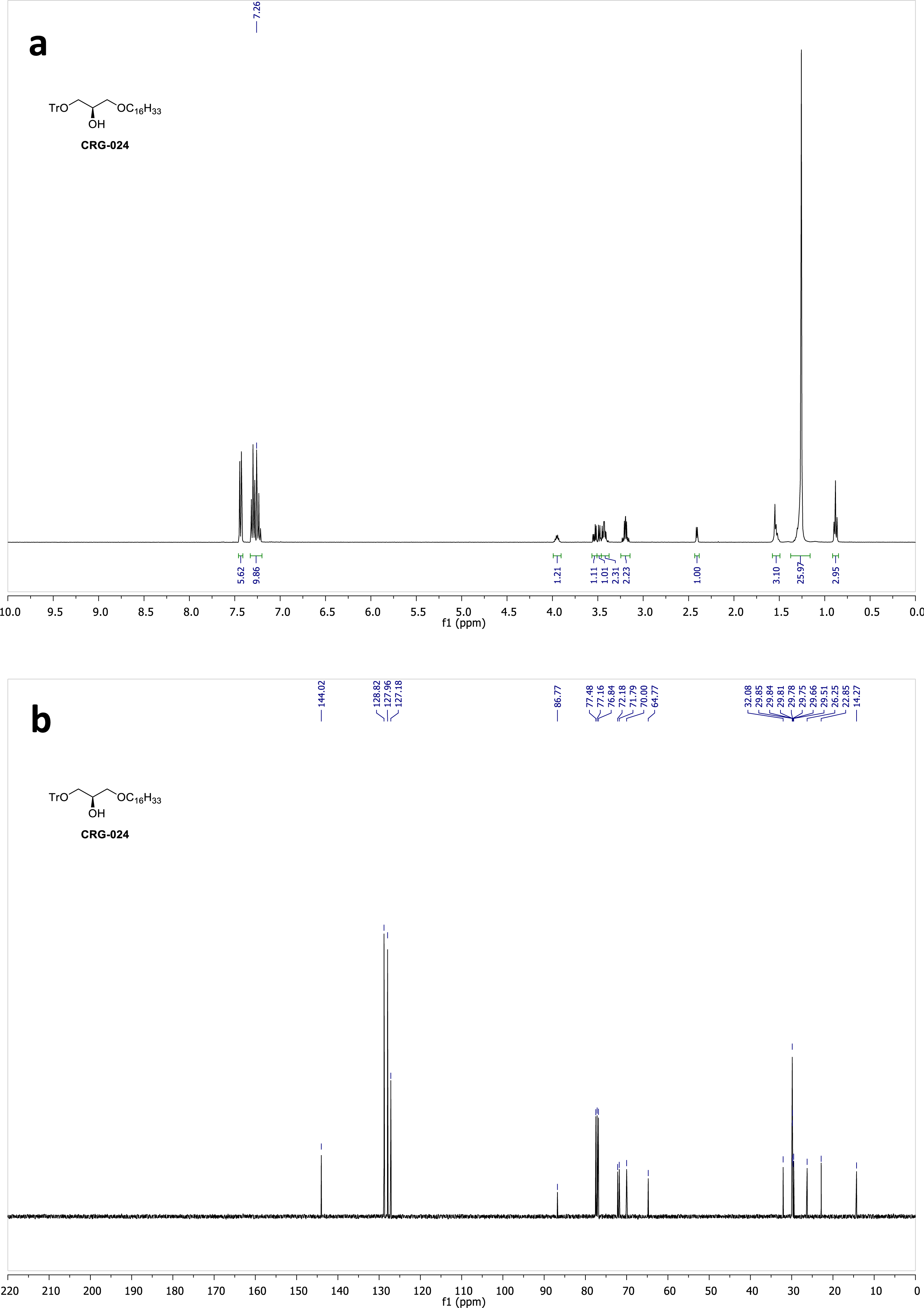
^1^H (a) and ^13^C (b) NMR spectra of compound **CRG024** in CDCl_3_.

**Figure S35.**
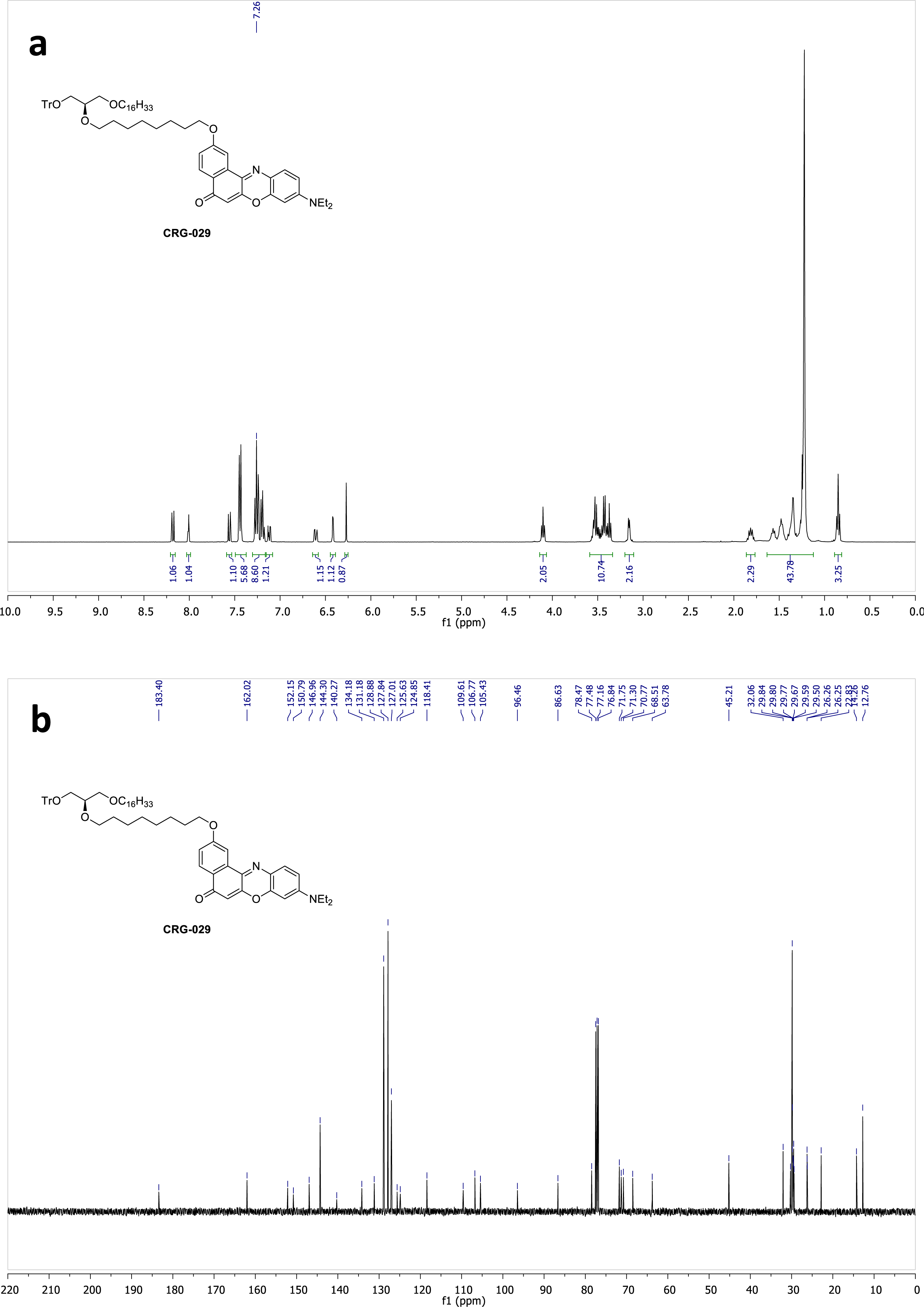
^1^H (a) and ^13^C (b) NMR spectra of compound **CRG029** in CDCl_3_.

**Figure S36.**
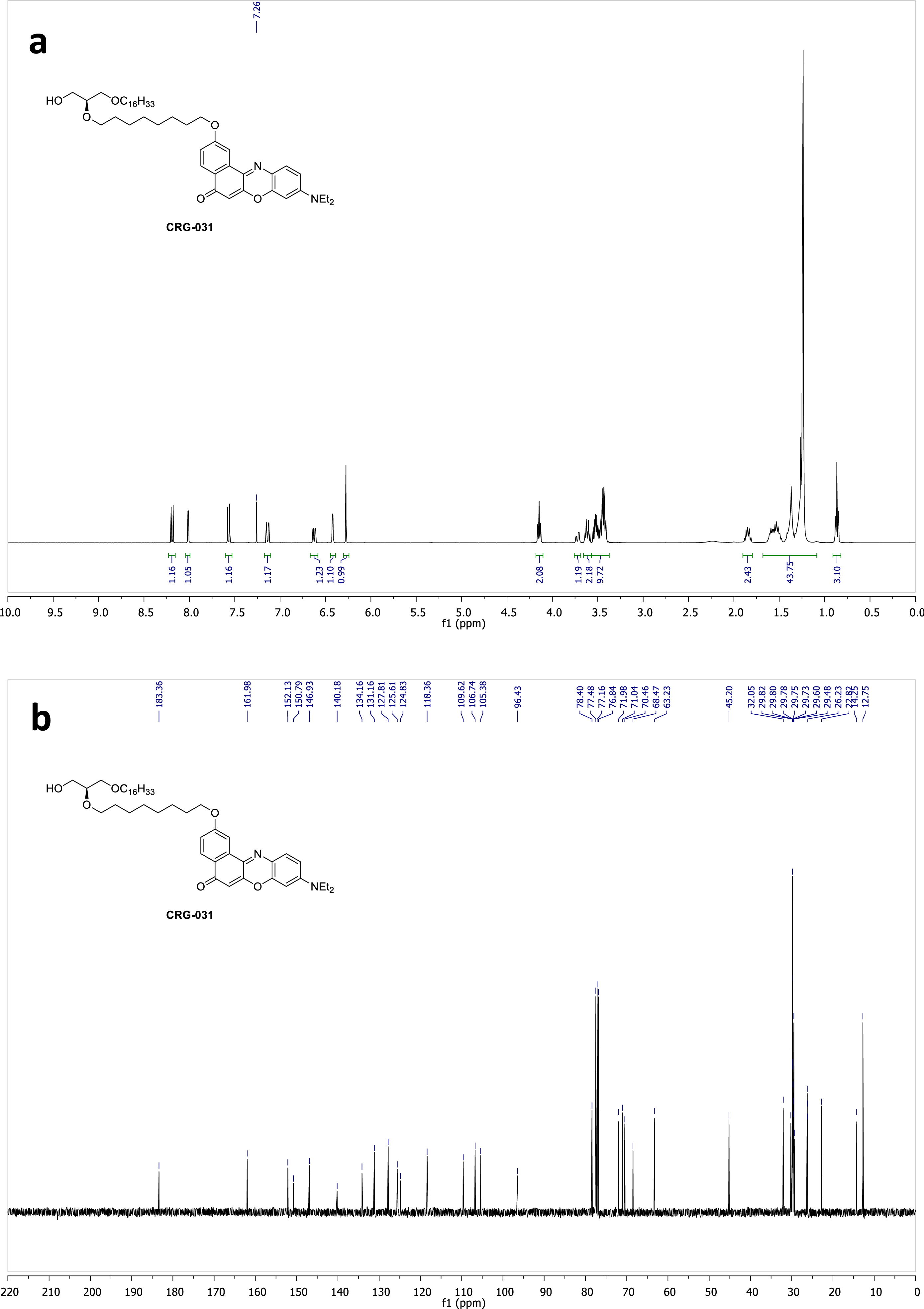
^1^H (a) and ^13^C (b) NMR spectra of compound **CRG031** in CDCl_3_.

**Figure S37.**
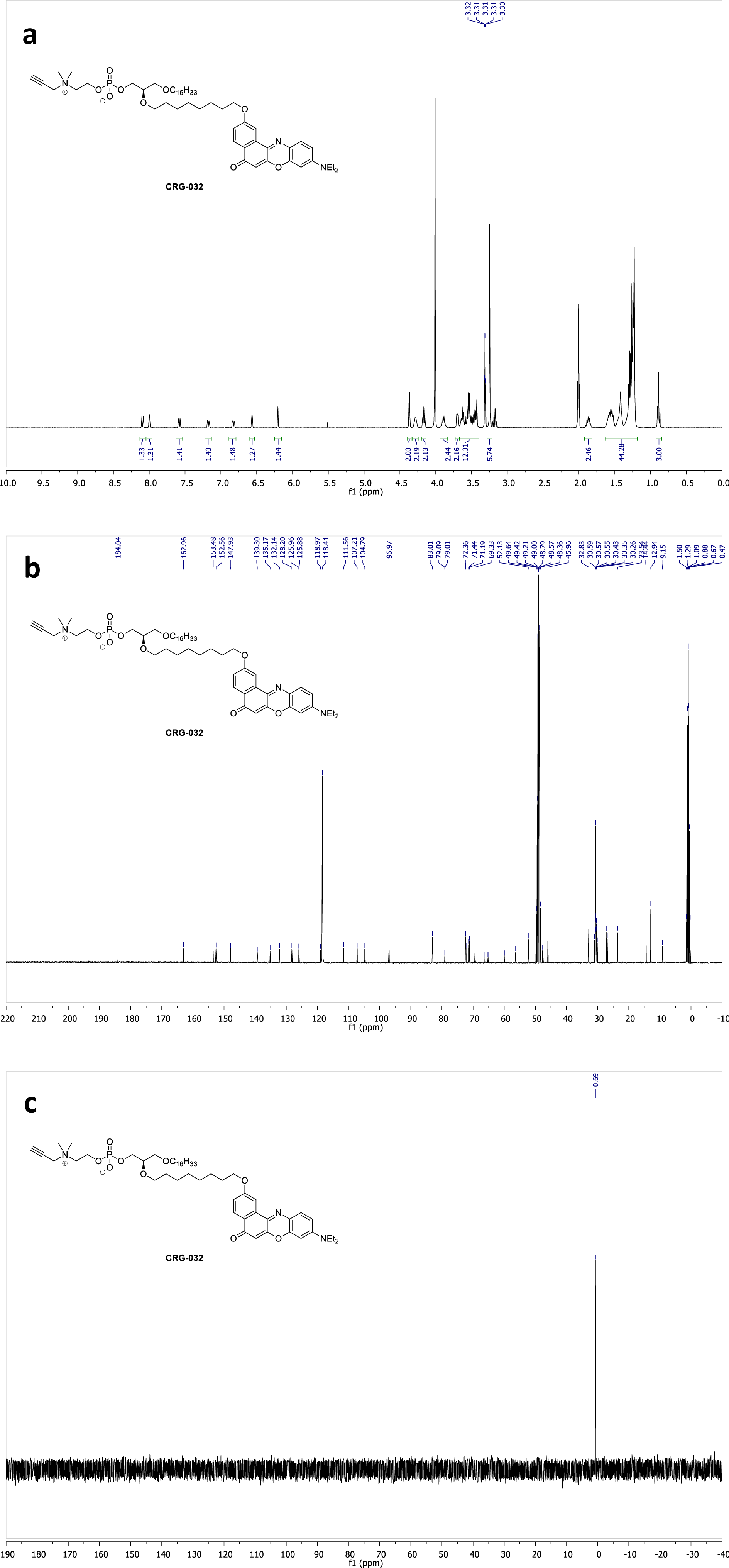
^1^H (a), ^13^C (b) and ^31^P (c) NMR spectra of compound **CRG032** in CD_3_OD/CD_3_CN (1:1).

**Figure S38.**
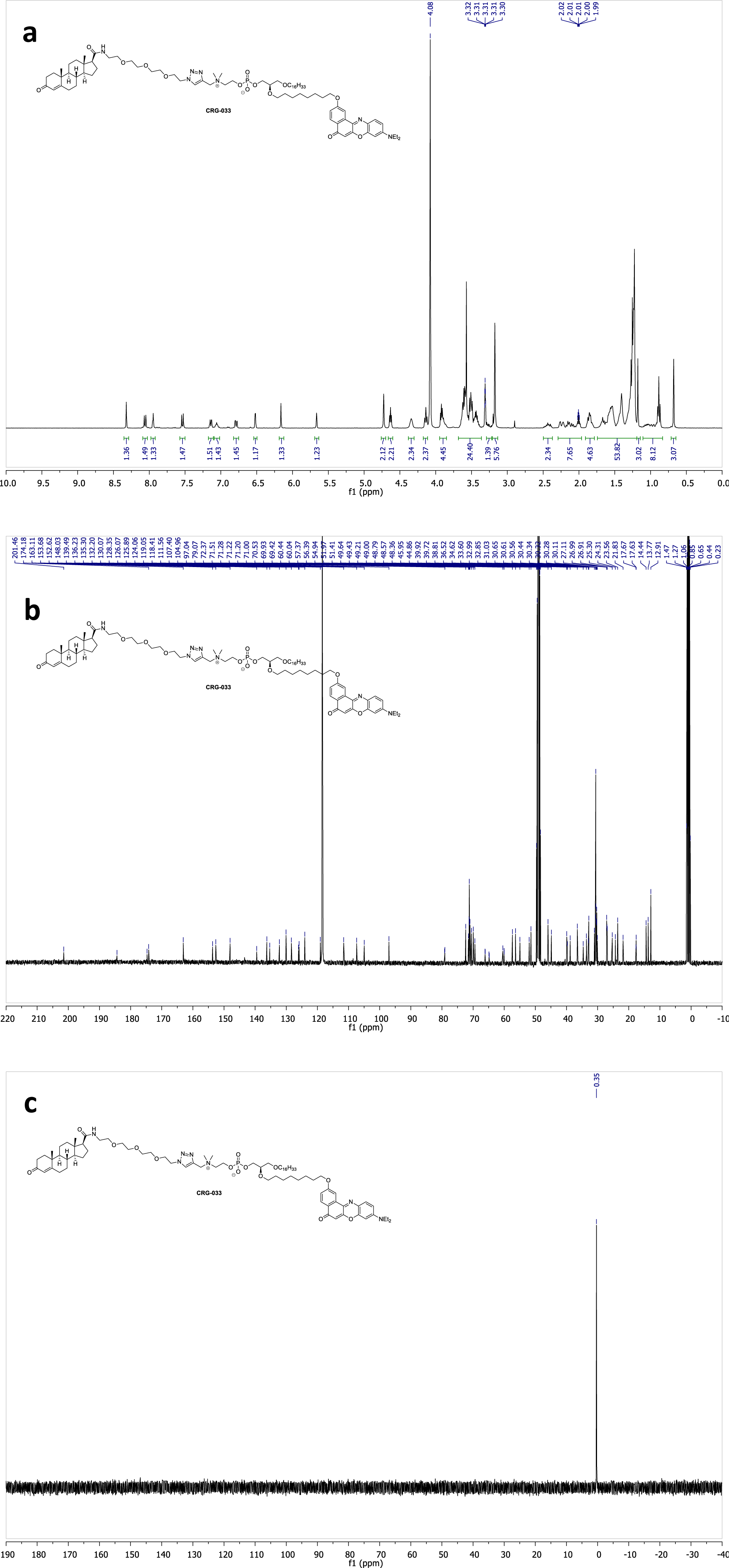
^1^H (a), ^13^C (b) and ^31^P (c) NMR spectra of compound **CRG033** in CD_3_OD/CD_3_CN (1:1).

**Figure S39.**
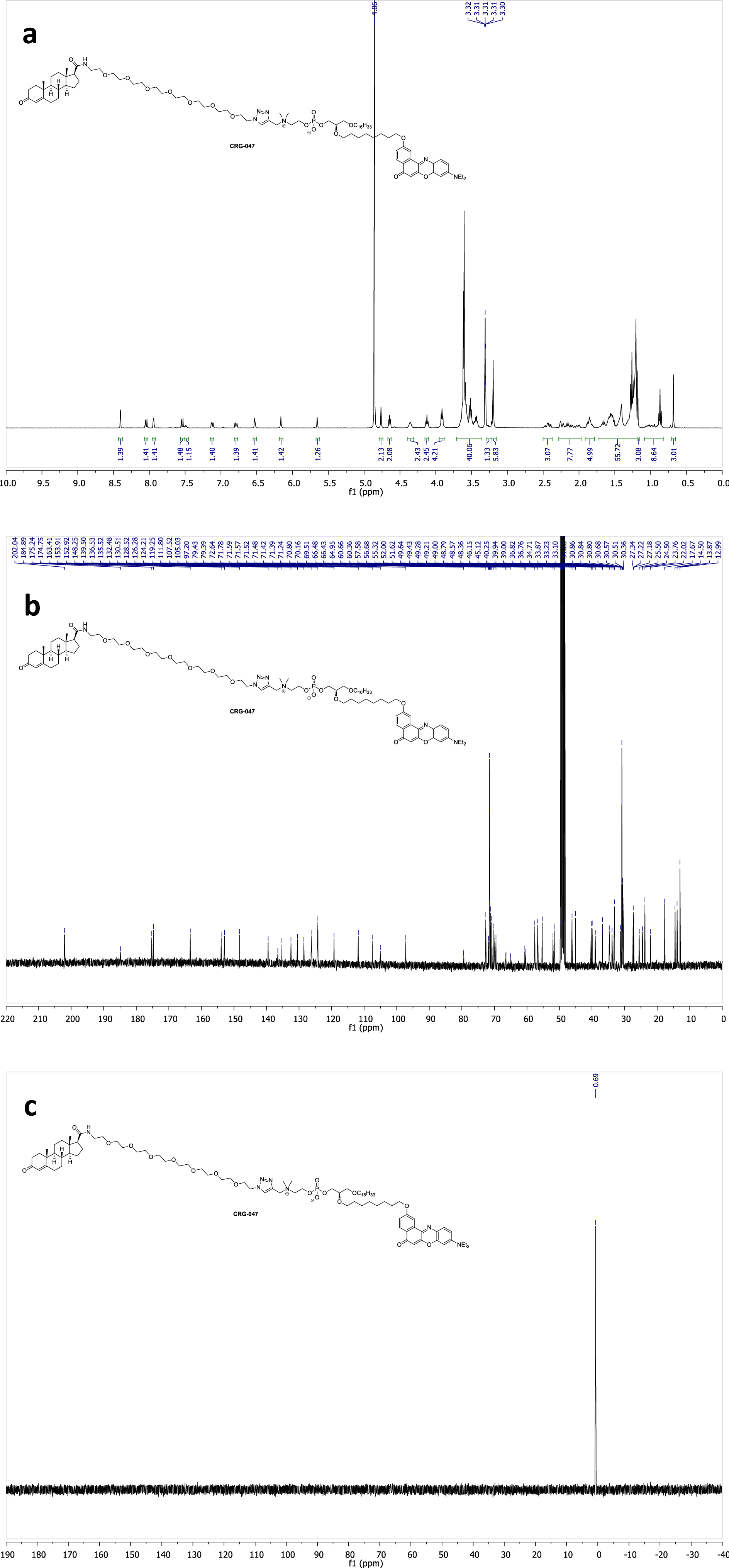
^1^H (a), ^13^C (b) and ^31^P (c) NMR spectra of compound **CRG047** in CD_3_OD.

**Figure S40.**
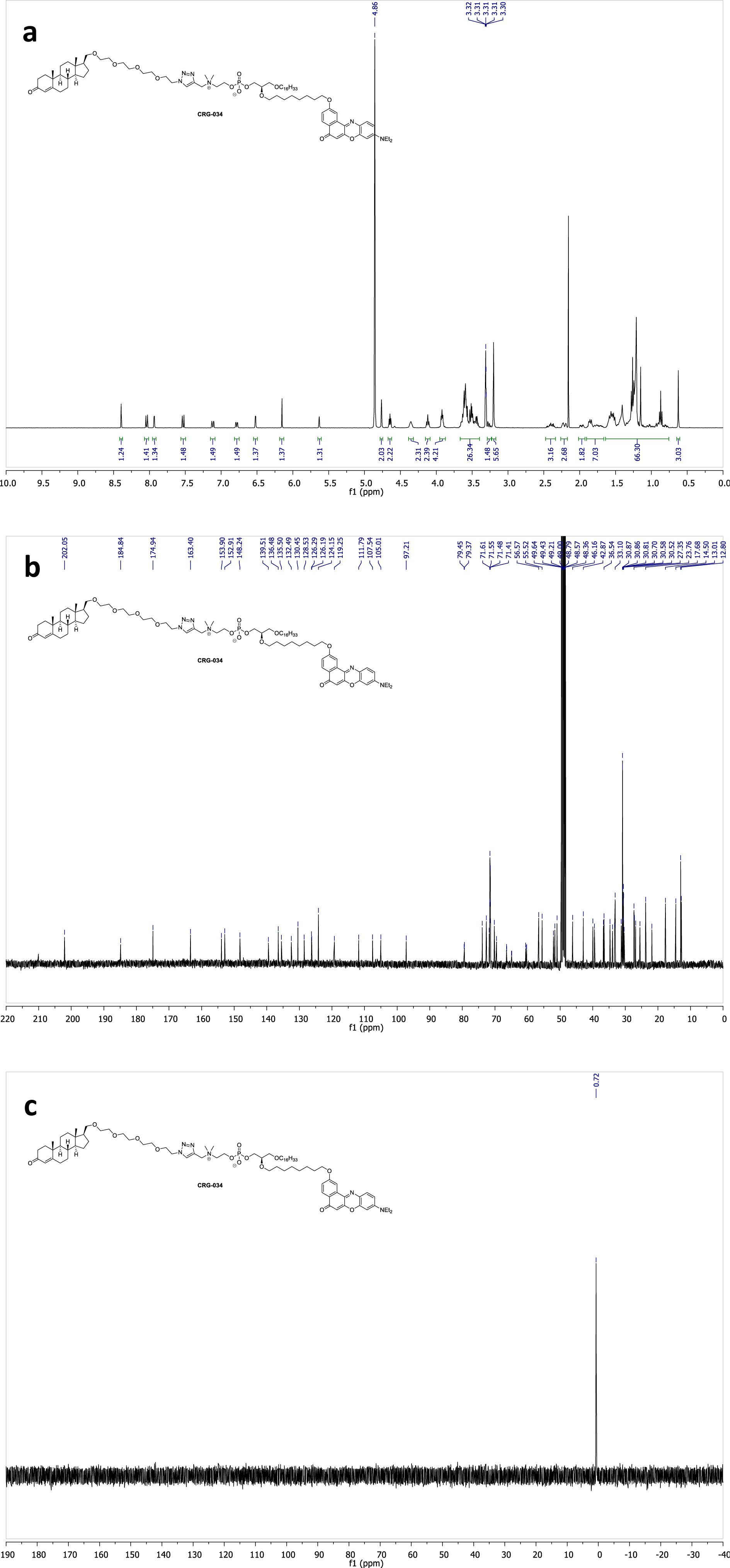
^1^H (a), ^13^C (b) and ^31^P (c) NMR spectra of compound **CRG034** in CD_3_OD.

**Figure S41.**
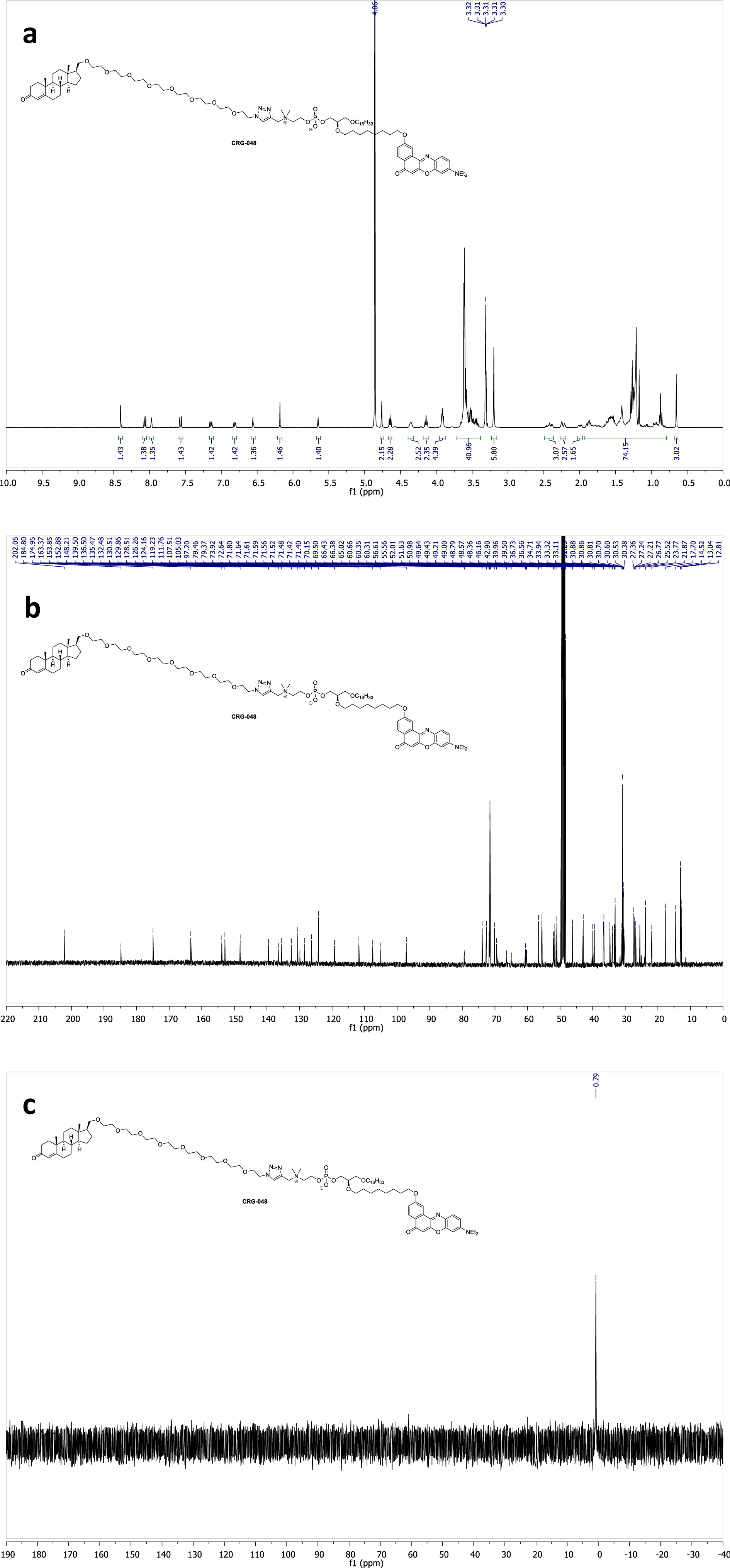
^1^H (a), ^13^C (b) and ^31^P (c) NMR spectra of compound **CRG048** in CD_3_OD.

**Table S2.**
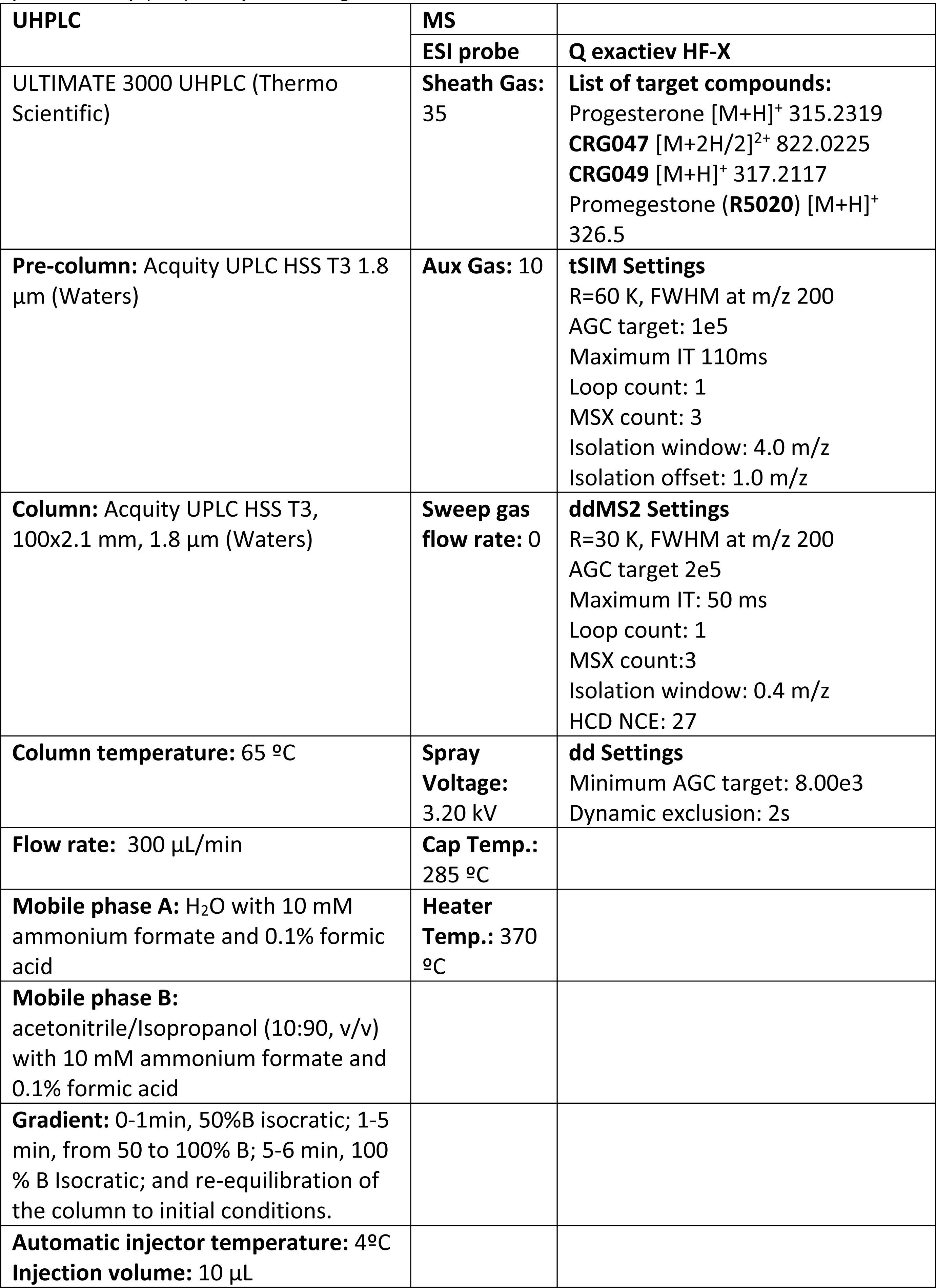
Ultra-High Performance Liquid Chromatography (UHPLC) and Mass Spectrometry (MS) analysis settings.

| UHPLC | MS |  |
| --- | --- | --- |
|  | ESI probe | Q exactiev HF-X |
| ULTIMATE 3000 UHPLC (Thermo Scientific) | **Sheath Gas:**<br>35 | **List of target compounds:**<br>Progesterone [M+H]<sup>+</sup> 315.2319<br>**CRG047** [M+2H/2]<sup>2+</sup> 822.0225<br>**CRG049** [M+H]<sup>+</sup> 317.2117<br>Promegestone (**R5020**) [M+H]<sup>+</sup> 326.5 |
| **Pre-column:** Acquity UPLC HSS T3 1.8 µm (Waters) | **Aux Gas:** 10 | **tSIM Settings**<br>R=60 K, FWHM at m/z 200<br>AGC target: 1e5<br>Maximum IT 110ms<br>Loop count: 1<br>MSX count: 3<br>Isolation window: 4.0 m/z<br>Isolation offset: 1.0 m/z |
| **Column:** Acquity UPLC HSS T3, 100x2.1 mm, 1.8 µm (Waters) | **Sweep gas flow rate:** 0 | **ddMS2 Settings**<br>R=30 K, FWHM at m/z 200<br>AGC target 2e5<br>Maximum IT: 50 ms<br>Loop count: 1<br>MSX count:3<br>Isolation window: 0.4 m/z<br>HCD NCE: 27 |
| **Column temperature:** 65 °C | **Spray Voltage:**<br>3.20 kV | **dd Settings**<br>Minimum AGC target: 8.00e3<br>Dynamic exclusion: 2s |
| **Flow rate:** 300 µL/min | **Cap Temp.:**<br>285 °C |  |
| **Mobile phase A:** H<sub>2</sub>O with 10 mM ammonium formate and 0.1% formic acid | **Heater Temp.:** 370 °C |  |
| **Mobile phase B:**<br>acetonitrile/Isopropanol (10:90, v/v) with 10 mM ammonium formate and 0.1% formic acid |  |  |
| **Gradient:** 0-1min, 50%B isocratic; 1-5 min, from 50 to 100% B; 5-6 min, 100 % B Isocratic; and re-equilibration of the column to initial conditions. |  |  |
| **Automatic injector temperature:** 4°C<br>**Injection volume:** 10 µL |  |  |

## SUPPORTING INFORMATION

### 1. Synthetic route for probes CRG033, CRG047, CRG034 and CRG048

#### 1.1. Synthesis of PEG-linkers CRG003, CRG042, CRG012 and CRG037

The synthesis of the heterobifunctional poly(ethylene glycol) (PEG) linkers CRG003, CRG042, CRG012 and CRG037 is depicted in Scheme S1. For the synthesis of amino-PEG-azides CRG003 and CRG042, both terminal hydroxyl groups of tetra or octaethylene glycol (compounds 1 and 2) were mesylated yielding the corresponding dimesyl intermediates which were subsequently transformed to the di-azido PEGs CRG002 and CRG041. Selective transformation of one of the azido groups to an amine *via* Staundinger reduction gave the desired heterobifunctional linkers CRG003 and CRG042.

On the other hand, desymmetrization of 1 and 2 by conversion of one of their hydroxyl groups to an azido functionality allowed the obtention of monoazide products CRG011 and CRG036 in moderate yields. Finally, treatment of the resulting alcohols with MsCl and TsCl, yielded the sulfonate esters CRG012 and CRG037, respectively.

**Scheme S1.**
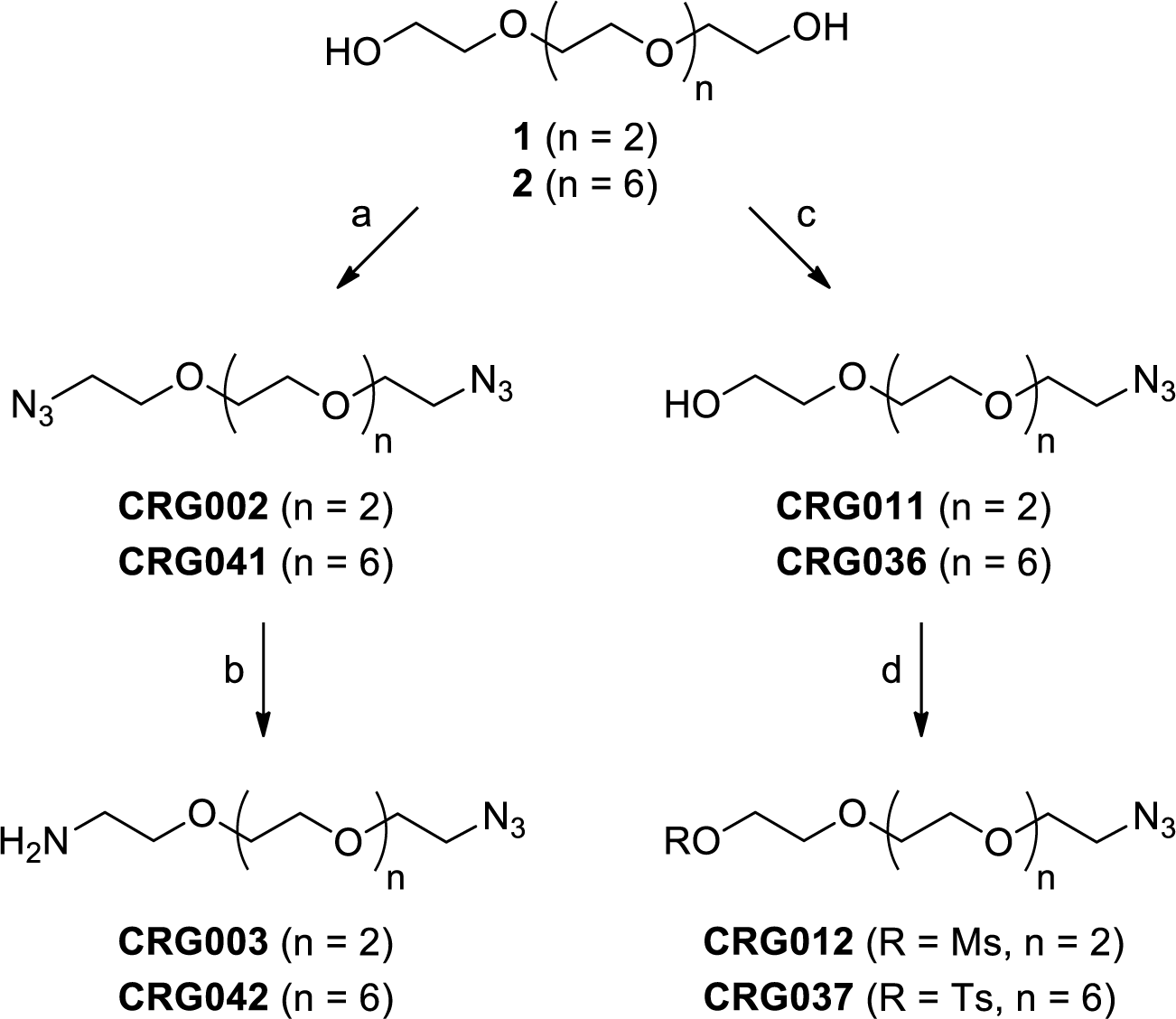
Synthesis of PEG-linkers CRG003, CRG042, CRG012 and CRG037. Reagents and conditions: (**a**) (i) MsCl, TEA, CH_2_Cl_2_, 0 °C to rt. (ii) NaN_3_, DMF, 65 °C, 97% for CRG002, quant. yield for CRG041; (**b**) PPh_3_, 0.5 M HCl /Et_2_O (1:1), rt, 75% for CRG003, 72% for CRG042; (**c**) (i) MsCl, TEA, CH_2_Cl_2_, 0 °C to rt. (ii) NaN_3_, DMF, 65 °C, 36% for CRG011, 41% for CRG036; (**d**) For CRG012: MsCl, TEA, CH_2_Cl_2_, 0 °C to rt, 94%. For CRG037: TsCl, KOH, CH_2_Cl_2_, 0 °C, 93%.

#### 1.2. Synthesis of steroidal moieties CRG006 and CRG013

With the required heterobifunctional linkers in hand, the differently functionalized steroidal scaffolds CRG006 and CRG013 were also prepared (Scheme S2). First, protection of the 3β-hydroxyl group of pregnenolone (**3**) as a *tert*-butyldimethylsilyl ether afforded compound CRG005, which was treated with sodium hypobromide in dioxane/water to give the corresponding 17β-carboxylic acid CRG006. Subsequent reduction of CRG006 using lithium aluminium hydride provided alcohol CRG013 in 61% yield over three steps.

**Scheme S2.**
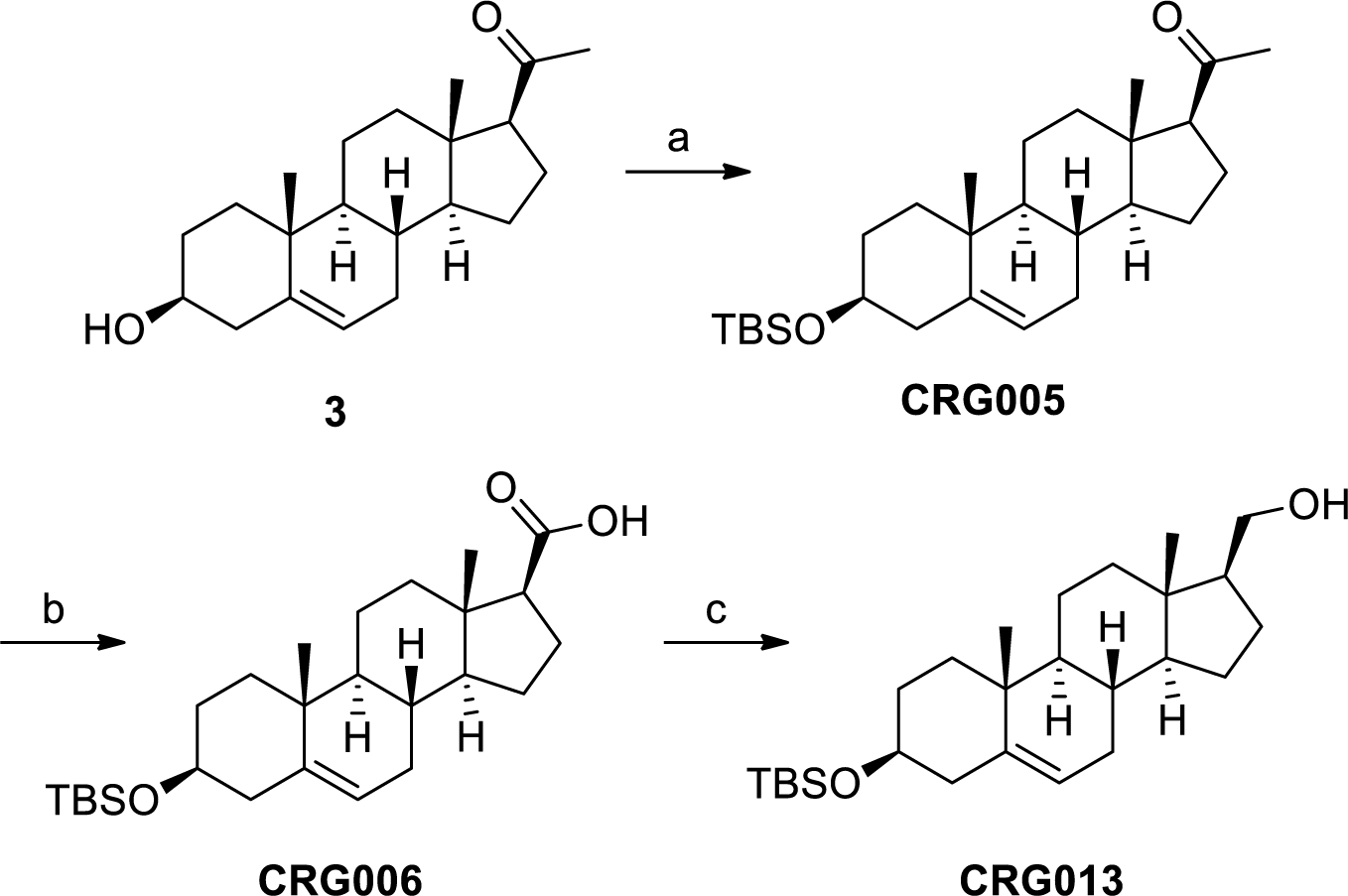
Synthesis of steroidal moieties CRG006 and CRG013. Reagents and conditions: (**a**) TBSCl, imidazole, DMF, 0 °C to rt, 94%; (**b**) NaOBr, H_2_O/dioxane, 0 °C to rt, 83%; (c) LiAlH_4_, THF, 0 °C to rt, 78%.

#### 1.3. Synthesis of azides CRG009, CRG045, CRG016 and CRG046

In order to obtain the series of probes with an amide bond between the steroidal moiety and the azido-terminated linker, amines CRG003 and CRG042 were acylated with carboxylic acid CRG006, yielding amides CRG007 and CRG043, respectively (Scheme S3). Alternatively, treatment of alcohol CRG013 with sulfonate esters CRG012 and CRG037 under basic conditions gave rise to the ether-linked compounds CRG014 and CRG038, respectively.

Although tosylate CRG037 appeared to be more reactive towards the alcoxide generated from CRG013 than its mesylate counterpart (compound CRG012), both reactions occurred in low to moderate yields. Apart from the low reactivity of the starting materials, the observed decrease in yield could also be attributed to the fact the TBS group presented certain lability under the reaction conditions, leading to the formation of considerable amounts of deprotected CRG013 during the course of the reaction (as seen by ^1^H/^13^C-NMR characterization of the crude reaction mixture).

Once both amide and ether-linked derivatives were obtained, the synthetic route continued with the TBAF-mediated removal of the TBS protecting group, which took place efficiently, affording the 3β-alcohols CRG008, CRG044, CRG015 and CRG039. Finally, Oppenauer oxidation of the hydroxy group at C3 (which occurred with concomitant isomerisation of the double bond) afforded the α,β-unsaturated ketones CRG009, CRG045, CRG016 and CRG046, respectively.

**Scheme S3.**
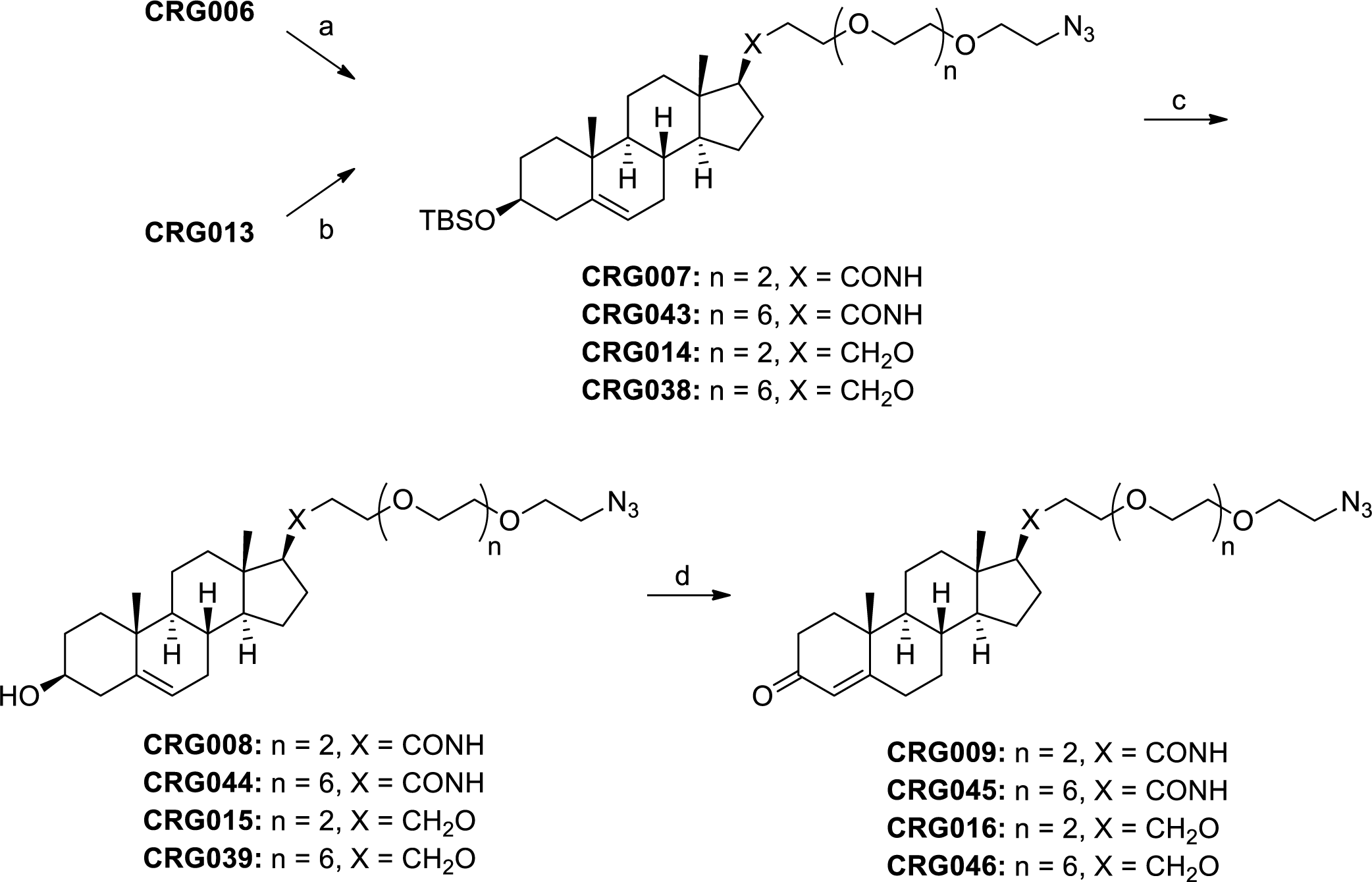
Synthesis of azides CRG009, CRG045, CRG016 and CRG046. Reagents and conditions: (**a**) Selected amine, EDC, HOBt, TEA, CH_2_Cl_2_, 0 °C to rt, 80% for CRG007, 92% for CRG043; (**b**) Selected sulfonate ester, NaH, THF, 65 °C, 24% for CRG014, 53% for CRG038; (**c**) TBAF, THF, reflux, 76-97%; (d) Al(O*^t^*Bu)_3_, acetone/toluene, reflux, 58-92%.

#### 1.4. Synthesis of probes CRG033, CRG047, CRG034 and CRG048

As depicted in Scheme S4, the synthesis of the alkyne-terminated phosphatidylcholine derivative CRG032 started with (*S*)-(−)-glycidol (**4**), which served as the source of chirality. After protection of **4** as the trityl ether CRG023, the epoxide was regioselectively opened with the *in situ* generated alcoxyde of hexadecanol, yielding alcohol CRG024 with retention of the configuration at the C2 position.

Treatment of CRG024 with an excess of 1,8-dibromooctane under basic conditions afforded an inseparable and enriched crude mixture of the bromo derivative 5, which was directly treated with the phenoxyde form of 2-hydroxy Nile Red,^1^ allowing the nucleophilic displacement of the remaining bromo group at the distal position of the side chain of the lipid structure and yielding compound CRG029 in 54% yield over two steps.

Acid-mediated removal of the trityl group gave compound CRG031 which was transformed to the zwitterionic aminophosphate CRG032 in a two-step sequence consisting of a phosphorylation of the primary hydroxyl at C1 followed by a nucleophilic opening of the resulting cyclic phosphate by 3-dimethylamino-1-propyne.^2, 3^

The desired probes were finally assembled by means of a Cu(I)-catalyzed azide-alkyne [3 + 2] dipolar cycloaddition (CuAAC) between the terminal alkyne in CRG032 and the azido group present in fragments CRG009, CRG045, CRG016 and CRG046, giving probes CRG033, CRG047, CRG034 and CRG048, respectively.

**Scheme S4.**
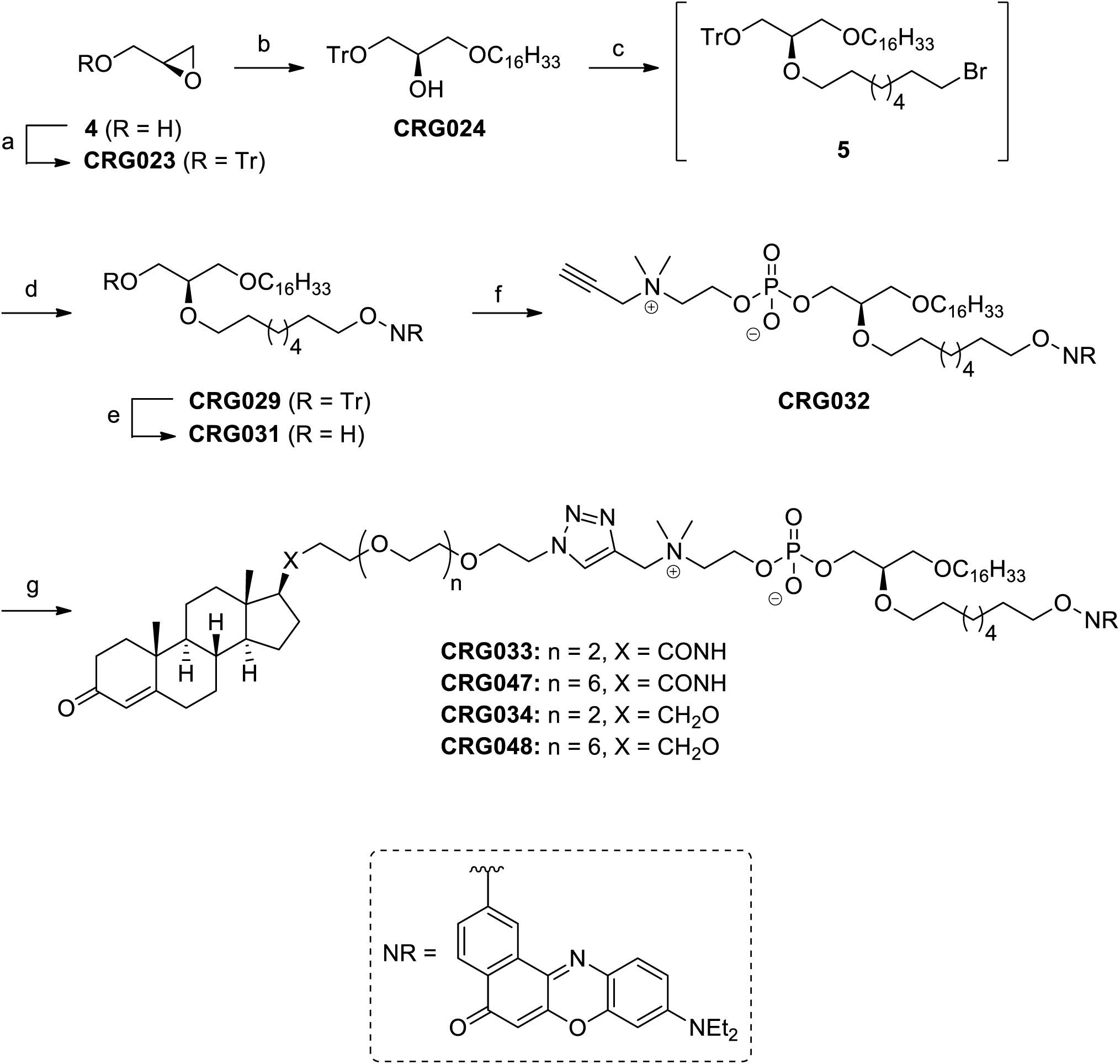
Synthesis of probes CRG033, CRG047, CRG034 and CRG048. Reagents and conditions: (**a**) TrCl, TEA, CH_2_Cl_2_, 0 °C to rt, 92%; (**b**) Hexadecanol, KO*^t^*Bu, DMF, 80 °C, 55%; (**c**) 1,8-dibromooctane, NaH, DMF, 65 °C; (**d**) 9-(diethylamino)-2-hydroxy-5*H*-benzo[*a*]phenoxazin-5-one (2-hydroxy Nile Red), K_2_CO_3_, DMF, 65 °C, 54% over two steps; (**e**) *p*-TsOH, MeOH/CHCl_3_, 0 °C to rt, 91%; (**f**) (I) 2-chloro-1,3,2-dioxaphospholane 2-oxide, TEA, DMAP, toluene, 0 °C to rt (II) 3-dimethylamino-1-propyne, MeCN, 80 °C, 42% over two steps; (**g**) Selected azide, CuI, DIPEA, H_2_O/MeCN (1:1), rt, 51-95%.

#### 1.5. Synthesis of carboxylic acid CRG049

Compound CRG049 was obtained (Scheme S5) by oxidation of the commercially available progesterone (**6**) following an adapted version of the synthetic protocol described by Lao *et al*.^4^

**Scheme S5.**
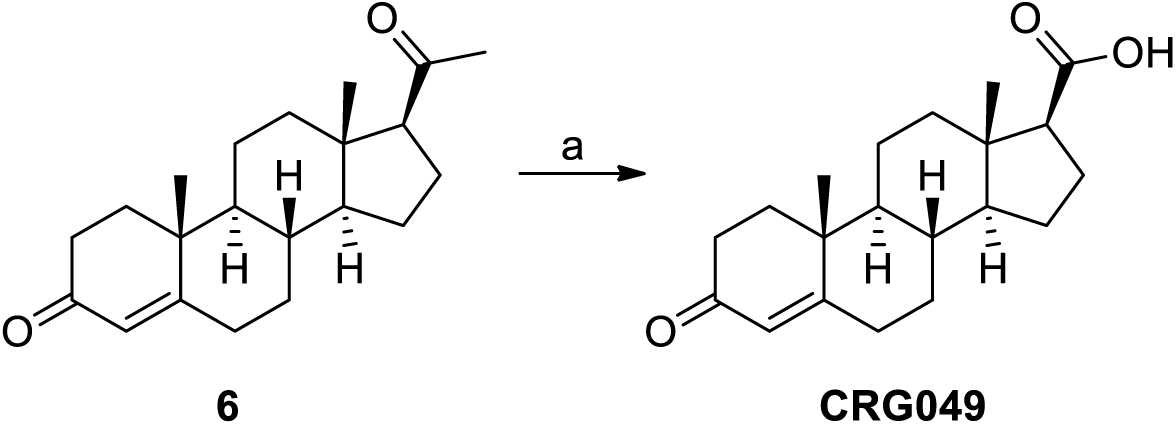
Synthesis of carboxylic acid CRG049. Reagents and conditions: (**a**) NaOBr, H_2_O/dioxane, 0 °C to rt, 74%.

### 2. Synthetic experimental part

#### 2.1. General remarks

Unless otherwise stated, reactions were carried out under nitrogen atmosphere. Dry solvents were obtained by passing through an activated alumina column on a Solvent Purification System (SPS). Acetone was dried over anhydrous CaCl_2_ and distilled prior to use. Commercially available reagents and solvents were used with no further purification. All reactions were monitored by TLC analysis using ALUGRAM^®^ SIL G/UV254 precoated aluminum sheets (Machery-Nagel). UV light was used as the visualizing agent and a 5% (w/v) ethanolic solution of phosphomolybdic acid as the developing agent. Flash column chromatography was carried out with the indicated solvents using flash-grade silica gel (37-70 μm). Yields refer to chromatographically and spectroscopically pure compounds, unless otherwise stated.

NMR spectra were recorded at room temperature on a Varian Mercury 400 instrument. The chemical shifts (δ) are reported in ppm relative to the solvent signal, and coupling constants (*J*) are reported in Hertz (Hz). ^31^P chemical shifts are relative to an 85 % H_3_PO_4_ external reference (0 ppm). In the case of NMR spectra recorded in CD_3_OD/CD_3_CN mixtures, chemical shifts are expressed relative to the residual peak of CD_3_OD. The following abbreviations are used to define the multiplicities in ^1^H NMR spectra: s = singlet, d = doublet, t = triplet, q = quartet, dd = doublet of doublets, ddd = doublet of doublet of doublets, dt = doublet of triplets, td = triplet of doublets, m = multiplet, br = broad signal and app = apparent. High Resolution Mass Spectrometry analyses were carried out on an Acquity UPLC system coupled to a LCT Premier orthogonal accelerated time-of-flight mass spectrometer (Waters) using the electrospray ionization (ESI) technique. Optical rotations were measured at room temperature on a Perkin Elmer 341 polarimeter.

#### 2.2. General synthetic methods

##### General procedure 1: Preparation of diazido derivatives from polyethylene glycols

MsCl (3.0 eq.) was added dropwise to a mixture of TEA (3.0 eq.) and the corresponding diol (15 mmol) in CH_2_Cl_2_ (100 mL) stirring at 0 °C. After the addition was complete, the reaction was allowed to proceed at 0 °C for 2 h and then at rt for a further 2 h. The reaction mixture was then poured into 0.5 M HCl and the mixture was extracted with CH_2_Cl_2_. The combined organic layers were washed with brine, dried over anhydrous MgSO_4_, filtered and evaporated to dryness. The resulting residue was dissolved in DMF (60 mL) and the mixture was treated with NaN_3_ (5.0 eq.) and stirred overnight at 65 °C. DMF was removed *in vacuo* and the residue was resuspended in Et_2_O, filtered over celite, and washed with Et_2_O. Finally, the combined filtrates were concentrated *in vacuo* to give the corresponding diazido derivatives, which were used without further purification.

##### General procedure 2: Monoreduction of diazido derivatives

To an ice-cooled solution of the corresponding diazido derivative (15 mmol) in 0.5 M HCl (60 mL) was added dropwise a solution of triphenyl phosphine (0.9 eq.) in Et_2_O (60 mL). After stirring overnight at rt, the reaction mixture was washed with EtOAc in order to remove the unreacted starting materials and triphenylphosphine oxide that was formed during the reaction. The aqueous layer was collected, cooled to 0 °C, and treated with 1M aq. KOH until the pH of the solution was around 12. This solution was finally extracted with CH_2_Cl_2_ and the combined organic layers were dried over anhydrous MgSO_4_, filtered and evaporated to dryness to give the required mono amines, which were used without further purification.

##### General procedure 3: Preparation of monoazido derivatives from polyethylene glycols

MsCl (0.9 eq.) was added dropwise to a mixture of TEA (1.5 eq.) and the corresponding diol (14 mmol) in CH_2_Cl_2_ (55 mL) stirring at 0 °C. After the addition was complete, the reaction was allowed to proceed at 0 °C for 1 h and then at rt for a further 2 h. The reaction mixture was then poured into 0.5 M HCl and the mixture was extracted with CH_2_Cl_2_. The combined organic layers were washed with brine, dried over anhydrous MgSO_4_, filtered and evaporated to dryness to give a mixture of mono- and di-mesylated derivatives. The resulting residue was dissolved in DMF (35 mL) and the mixture was treated with NaN_3_ (5.0 eq.) and stirred over night at 65 °C. DMF was removed *in vacuo* and the residue was resuspended in Et_2_O, filtered over celite, and washed with Et_2_O. Finally, the combined filtrates were concentrated under reduced pressure to give a crude mixture, which was purified as indicated for each compound.

##### General procedure 4: EDC/HOBt-mediated coupling for amide-bond formation

HOBt (1.2 eq.) and EDC (1.2 eq.) were added portionwise to a solution of **CRG006** (0.75 mmol) in CH_2_Cl_2_ (15 mL) at 0 °C. The resulting solution was stirred at the same temperature for 15 min and was added dropwise to an ice-cooled solution of the selected amine (1.1 eq.) and TEA (1.5 eq.) in CH_2_Cl_2_ (10 mL). After stirring at rt for 2h, the reaction mixture was diluted with water and was extracted with CH_2_Cl_2_. The combined organic layers were washed with brine, dried over anhydrous MgSO_4_, filtered and evaporated *in vacuo* to give a residue, which was purified as indicated for each compound.

##### General procedure 5: fluoride-mediated deprotection of TBS groups

To a solution of the corresponding TBS-protected alcohol (0.5 mmol) in THF (8.5 mL) was added dropwise TBAF (1 M in THF, 2 eq.) at 0 °C. The reaction mixture was refluxed for 4h, then cooled down to rt and quenched with saturated aq. NH_4_Cl (10 mL). The resulting mixture was extracted with Et_2_O and the combined organic layers were washed with brine, dried over anhydrous MgSO_4_, filtered, and evaporated to give the crude products. Purification by flash chromatography on silica gel using the indicated conditions afforded the required alcohols.

##### General procedure 6: Oppenauer oxidation of allylic alcohols

Aluminum *tert*-butoxide (1.5 eq.) was added in one portion to a solution of the corresponding allylic alcohol (0.5 mmol) in acetone (3 mL) and toluene (7 mL). The reaction mixture was refluxed for 7 h and cooled to room temperature. H_2_O was then added and the mixture was extracted with Et_2_O. The organic layers were combined, washed with brine, dried over anhydrous MgSO_4_, filtered and evaporated to give a crude mixture, which was purified as indicated for each compound.

##### General procedure 7: Williamson ether synthesis

To a solution of alcohol CRG013 (0.3 mmol) in THF (2 mL) was added portionwise NaH (60 wt.% in mineral oil, 2.0 eq.) at 0 °C and the mixture was stirred at the same temperature for 10 min. A solution of the corresponding sulfonate ester (1.0 eq.) in THF (1 mL) was then added dropwise and the resulting mixture was stirred at 65 °C overnight. After cooling down to rt, the reaction mixture was carefully diluted with water and extracted with Et_2_O. The combined organic layers were washed with brine, dried over anhydrous MgSO_4_, filtered and evaporated to give a residue, which was purified as indicated for each compound.

##### General procedure 8: CuAAC between CRG032 and selected azides

To a solution of CRG032 (0.025 mmol) and the selected azide (1.15 eq.) in H_2_O/MeCN (1:1) (2 mL), DIPEA (2 eq.) and CuI (0.15 eq.) were added. After stirring at rt for 1 h, the reaction mixture was diluted with water (5 mL) and was extracted with CHCl_3_. The combined organic layers were washed with brine, dried over anhydrous MgSO_4_ and concentrated to give a residue which was purified as indicated for each compound.

#### 2.3. Synthesis and characterization of compounds

##### 1- azido-2-(2-(2-(2-azidoethoxy)ethoxy)ethoxy)ethane (CRG002)

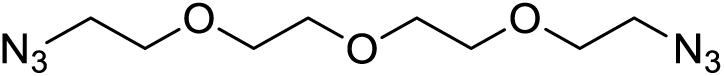

Compound **CRG002** (colourless oil, 3.66 g, 97%) was obtained from tetraethylene glycol (3.00 g, 15.45 mmol), MsCl (3.59 mL, 46.34 mmol), TEA (6.46 mL, 46.34 mmol) and NaN_3_ (5.02 g, 77.20 mmol), according to general procedure 1.

^1^H NMR (400 MHz, CDCl_3_) δ 3.71 – 3.64 (m, 12H), 3.39 (t, *J* = 5.1 Hz, 4H). ^13^C NMR (101 MHz, CDCl_3_) δ 70.8, 70.8, 70.2, 50.8. HRMS calcd. for C_8_H_16_N_6_O_3_Na ([M + Na]^+^): 267.1182, found: 267.1176.

##### 1,23-diazido-3,6,9,12,15,18,21-heptaoxatricosane (CRG041)

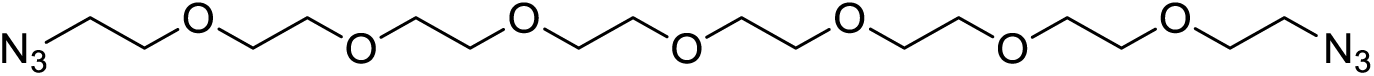

Compound **CRG041** (colourless oil, 205 mg, quant.) was obtained from octaethylene glycol (180 mg, 0.486 mmol), MsCl (113 µL, 1.46 mmol), TEA (203 µL, 1.46 mmol) and NaN_3_ (158 mg, 0.486 mmol), according to general procedure 1.

^1^H NMR (400 MHz, CDCl_3_) δ 3.69 – 3.62 (m, 28H), 3.38 (t, *J* = 5.1 Hz, 4H). ^13^C NMR (101 MHz, CDCl_3_) δ 70.7, 70.7, 70.6, 70.6, 70.0, 50.7. HRMS calcd. for C_16_H_32_N_6_O_7_Na ([M + Na]^+^): 443.2230, found: 443.2236.

##### 2-(2-(2-(2-azidoethoxy)ethoxy)ethoxy)ethanamine (CRG003)

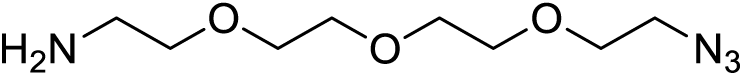

Compound **CRG003** (pale yellow oil, 2.44 g, 75%) was obtained from **CRG002** (3.65 g, 14.94 mmol) and PPh_3_ (3.53 g, 13.45 mmol), according to general procedure 2.

^1^H NMR (400 MHz, CDCl_3_) δ 3.70 – 3.61 (m, 10H), 3.51 (t, *J* = 5.2 Hz, 2H), 3.39 (t, *J* = 4.0 Hz, 2H),

2-86 (t, *J* = 5.2 Hz, 2H). ^13^C NMR (101 MHz, CDCl_3_) δ 73.5, 70.8, 70.7, 70.7, 70.4, 70.1, 50.8, 41.8. HRMS calcd. for C_8_H_19_N_4_O_3_ ([M + H]^+^): 219.1457, found: 219.1441.

##### 23-azido-3,6,9,12,15,18,21-heptaoxatricosan-1-amine (CRG042)

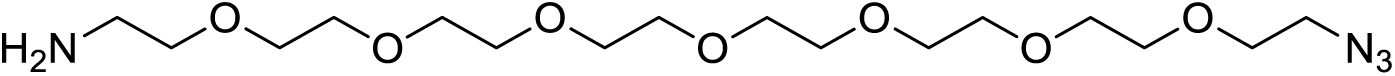

Compound **CRG042** (colourless oil, 168 mg, 72%) was obtained from **CRG041** (250 mg, 0.595 mmol) and PPh_3_ (141 mg, 0.535 mmol), according to general procedure 2.

^1^H NMR (400 MHz, CDCl_3_) δ 3.85 – 3.57 (m, 26H), 3.50 (t, *J* = 5.2 Hz, 2H), 3.37 (t, *J* = 4.0 Hz, 2H), 2.85 (t, *J* = 5.0 Hz, 2H). ^13^C NMR (101 MHz, CDCl_3_) δ 73.3, 70.7, 70.7, 70.6, 70.6, 70.6, 70.3, 70.0, 50.7, 41.8. HRMS calcd. for C_16_H_35_N_4_O_7_ ([M + H]^+^): 395.2506, found: 395.2492.

##### 2-(2-(2-(2-azidoethoxy)ethoxy)ethoxy)etanol (CRG011)

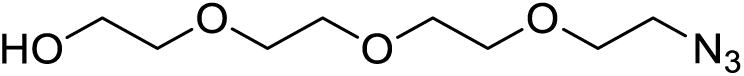

Compound **CRG011** (colourless oil, 1.09 g, 36%) was obtained from tetraethylene glycol (2.7 g, 13.90 mmol), MsCl (968 µL, 12.51 mmol), TEA (2.91 mL, 20.85 mmol) and NaN_3_ (4.52 g, 69.50 mmol), according to general procedure 3. The title compound was purified by flash chromatography on silica gel (from 0 to 2.5% MeOH in CH_2_Cl_2_).

^1^H NMR (400 MHz, CDCl_3_) δ 3.75 – 3.71 (m, 2H), 3.71 – 3.64 (m, 10H), 3.64 – 3.59 (m, 2H), 3.40 (t, *J* = 5.1 Hz, 2H), 2.13 (br s, 1H). ^13^C NMR (101 MHz, CDCl_3_) δ 72.6, 70.8, 70.7, 70.7, 70.4, 70.1, 61.8, 50.7. HRMS calcd. for C_8_H_17_N_3_O_4_Na ([M + Na]^+^): 242.1117, found: 242.1128.

##### 23-azido-3,6,9,12,15,18,21-heptaoxatricosan-1-ol (CRG036)

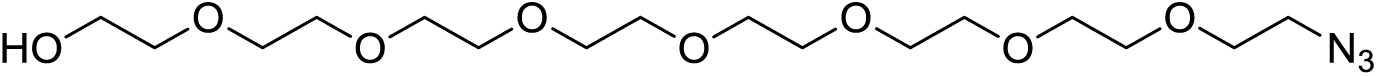

Compound **CRG036** (colourless oil, 110 mg, 41%) was obtained from octaethylene glycol (250 mg, 0.675 mmol), MsCl (47 µL, 0.607 mmol), TEA (141 µL, 1.01 mmol) and NaN_3_ (220 mg, 3.38 mmol), according to general procedure 3. The title compound was purified by flash chromatography on silica gel (from 0 to 6% MeOH in CH_2_Cl_2_).

^1^H NMR (400 MHz, CDCl_3_) δ 3.73 – 3.68 (m, 2H), 3.68 – 3.61 (m, 26H), 3.61 – 3.56 (m, 2H), 3.38 (t, *J* = 4.0 Hz, 2H). ^13^C NMR (101 MHz, CDCl_3_) δ 72.7, 70.7, 70.7, 70.7, 70.6, 70.6, 70.6, 70.6, 70.5, 70.3, 70.1, 61.7, 50.7. HRMS calcd. for C_16_H_34_N_3_O_8_ ([M + H]^+^): 396.2346, found: 396.2319.

##### 2-(2-(2-(2-azidoethoxy)ethoxy)ethoxy)ethyl methanesulfonate (CRG012)

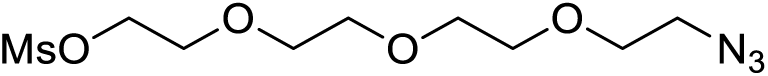

TEA (1.06 mL, 7.64 mmol) and MsCl (443 µL, 5.73 mmol) were added dropwise to a solution of **CRG011** (837 mg, 3.82 mmol) in CH_2_Cl_2_ (35 mL) at 0 °C. After stirring at rt for 3.5 h, the reaction mixture was poured into 0.5 M HCl and the mixture was extracted with CH_2_Cl_2_. The combined organic layers were washed with brine, dried over anhydrous MgSO_4_, filtered and evaporated to dryness. Purification of the residue by flash chromatography (from 0 to 70% EtOAc in hexane) gave **CRG012** (1.07 g, 94%) as a colourless oil.

^1^H NMR (400 MHz, CDCl_3_) δ 4.42 – 4.34 (m, 2H), 3.80 – 3.74 (m, 2H), 3.71 – 3.62 (m, 10H), 3.39 (t, *J* = 4.0 Hz, 2H), 3.07 (s, 3H). ^13^C NMR (101 MHz, CDCl_3_) δ 70.8, 70.7, 70.7, 70.1, 69.4, 69.1, 50.8, 37.8. HRMS calcd. for C_9_H_19_N_3_O_6_NaS ([M + Na]^+^): 320.0892, found: 320.0888.

##### 23-azido-3,6,9,12,15,18,21-heptaoxatricosyl 4-methylbenzenesulfonate (CRG037)

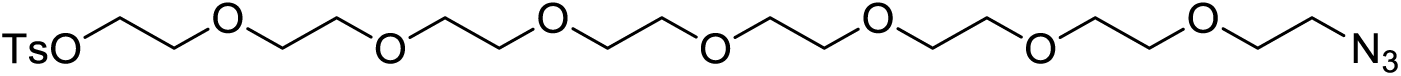

To an ice-cooled solution of **CRG036** (100 mg, 0.253 mmol) in CH_2_Cl_2_ (5 mL) was added KOH (57 mg, 1.01 mmol) followed by TsCl (72 mg, 0.379 mmol) and the reaction mixture was stirred at 0 °C for 4 h. A mixture of ice and water was then added and the mixture was extracted with CH_2_Cl_2_. The combined organic layers were dried over MgSO_4_, filtered and concentrated under reduced pressure. Flash chromatography of the residue (from 0 to 3.5% MeOH in CH_2_Cl_2_) gave **CRG037** (129 mg, 93%) as a colorless oil.

^1^H NMR (400 MHz, CDCl_3_) δ 7.80 (d, *J* = 8.3 Hz, 2H), 7.34 (d, *J* = 8.5 Hz, 2H), 4.19 – 4.12 (m, 2H), 3.71 – 3.54 (m, 28H), 3.39 (t, *J* = 4.0 Hz, 2H), 2.45 (s, 3H). ^13^C NMR (101 MHz, CDCl_3_) δ 144.9, 133.1, 129.9, 128.1, 70.9, 70.8, 70.8, 70.7, 70.7, 70.7, 70.6, 70.1, 69.4, 68.8, 50.8, 21.8. HRMS calcd. for C_23_H_40_N_3_O_10_S ([M + H]^+^): 550.2434, found: 550.2464.

##### 3β-(*tert*-Butyldimethylsilyloxy)-pregn-5-en-20-one (CRG005)

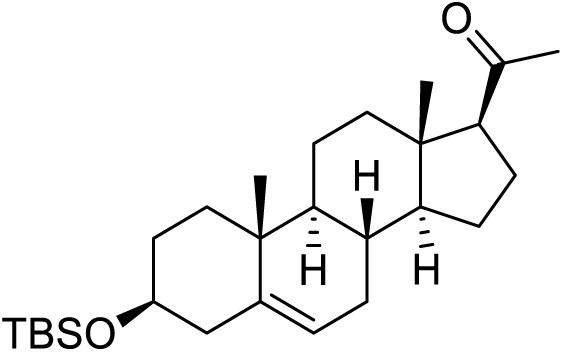

To a stirred solution of pregnenolone (**3**) (3.00 g, 9.48 mmol) in DMF (30 mL) was added imidazole (2.00 g, 29.39 mmol) and TBSCl (2.21 g, 14.69 mmol) at 0 °C. After stirring at rt for 16 h, the reaction mixture was diluted with water and extracted with Et_2_O. The combined organic layers were washed with brine, dried over anhydrous MgSO_4_, filtered and evaporated to dryness. The crude residue was purified by flash chromatography (from 0 to 5% EtOAc in hexane) to afford **CRG005** (3.82 g, 94%) as a white solid.

[α]^20^_D_ = +21.0 (*c* 1.0, CHCl_3_). ^1^H NMR (400 MHz, CDCl_3_) δ 5.32 (dd, *J* = 3.0, 2.3 Hz, 1H), 3.48 (ddd, *J* = 15.7, 10.9, 4.7 Hz, 1H), 2.53 (t, *J* = 9.0 Hz, 1H), 2.12 (s, 3H), 2.32 – 0.92 (m, 19H), 1.00 (s, 3H), 0.89 (s, 9H), 0.63 (s, 3H), 0.06 (s, 6H). ^13^C NMR (101 MHz, CDCl_3_) δ 209.6, 141.7, 121.0, 72.7, 63.9, 57.1, 50.2, 44.2, 42.9, 39.0, 37.5, 36.8, 32.2, 32.0, 32.0, 31.7, 26.1, 24.6, 23.0, 21.2, 19.6, 18.4, 13.4, −4.4. HRMS calcd. for C_27_H_48_O_2_Si ([M + H]^+^): 431.3345, found: 431.3337.

##### 3β-(*tert*-Butyldimethylsilyloxy)-androst-5-en-17β-carboxylic acid (CRG006)

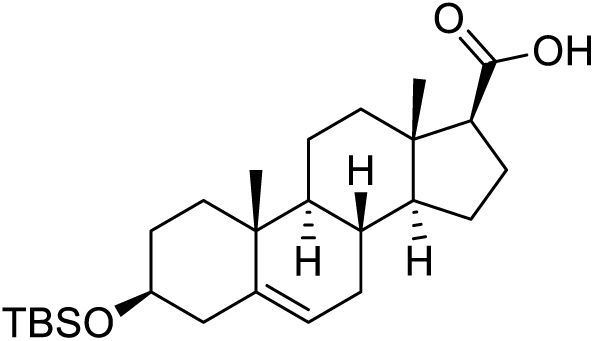

To a stirred solution of NaOH (6.41 g, 160.19 mmol) in water (55 mL) and dioxane (35 mL) was added dropwise Br_2_ (2.05 mL, 40.05 mmol) at 0 °C and the mixture was stirred at the same temperature for 10 min. 65 mL of this solution were added to an ice-cooled solution of **CRG005** (1.00 g, 2.32 mmol) in dioxane (65 mL) and water (8 mL) and the mixture was stirred overnight at rt. After cooling down to 0 °C, the reaction mixture was quenched with saturated aqueous Na_2_SO_3_, acidified with 1 M HCl and extracted with EtOAc. The combined organic layers were washed with brine, dried over anhydrous MgSO_4_, filtered and evaporated to dryness. Purification of the residue by flash chromatography (from 0 to 20% EtOAc in hexane) provided **CRG006** (830 mg, 83%) as a white solid.

[α]^20^_D_ = –17.8 (*c* 1.0, CHCl_3_). ^1^H NMR (400 MHz, CDCl_3_) δ 5.31 (dd, *J* = 3.3, 1.9 Hz, 1H), 3.48 (ddd, *J* = 15.7, 11.0, 4.8 Hz, 1H), 2.39 (t, *J* = 9.3 Hz, 1H), 2.33 – 0.92 (m, 19H), 1.00 (s, 3H), 0.89 (s, 9H), 0.74 (s, 3H), 0.06 (s, 6H). ^13^C NMR (101 MHz, CDCl_3_) δ 180.1, 141.8, 120.9, 72.7, 56.5, 55.3, 50.3, 44.3, 42.9, 38.2, 37.5, 36.8, 32.2, 32.0, 26.1, 24.7, 23.6, 21.0, 19.6, 18.4, 13.3, −4.4. HRMS calcd. for C_26_H_43_O_3_Si ([M – H]^−^): 431.2981, found: 431.2972.

##### 3β-(*tert*-Butyldimethylsilyloxy)-17β-(hydroxymethyl)-androst-5-ene (CRG013)

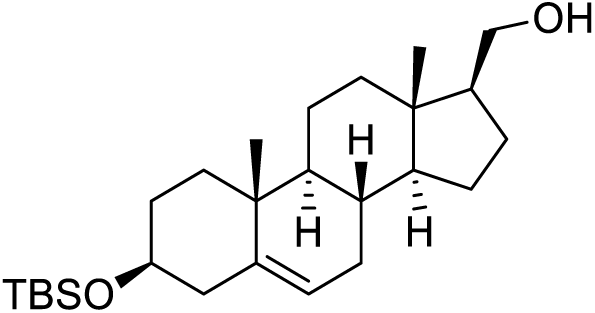

To a stirred suspension of LiAlH_4_ (179 mg, 4.71 mmol) in THF (3 mL) was added dropwise a solution of **CRG006** (680 mg, 1.57 mmol) in THF (6 mL) at 0 °C. The reaction mixture was stirred at rt for 16 h, cooled down to 0 °C and quenched by the dropwise addition of water (10 mL). The resulting white slurry was filtered over celite and washed with Et_2_O. The combined filtrates were dried over anhydrous MgSO_4_, filtered and concentrated under reduced pressure to give the crude product. Purification by flash chromatography (from 0 to 12% EtOAc in hexane) afforded **CRG013** (515 mg, 78%) as a white solid.

[α]^20^ = –45.3 (*c* 1.0, CHCl_3_). ^1^H NMR (400 MHz, CDCl_3_) δ 5.32 (dd, *J* = 3.2, 2.1 Hz, 1H), 3.73 (dd, *J* = 10.5, 6.9 Hz, 1H), 3.55 (dd, *J* = 10.5, 7.5 Hz, 1H), 3.48 (ddd, *J* = 15.6, 10.9, 4.7 Hz, 1H), 2.32 – 0.91 (m, 20H), 1.01 (s, 3H), 0.89 (s, 9H), 0.66 (s, 3H), 0.06 (s, 6H). ^13^C NMR (101 MHz, CDCl_3_) δ 141.8, 121.2, 72.8, 64.8, 56.4, 53.1, 50.6, 43.0, 41.8, 38.8, 37.6, 36.8, 32.2, 32.1, 31.8, 26.1, 25.7, 24.8, 20.9, 19.6, 18.4, 12.6, −4.4. HRMS calcd. for C_26_H_47_O_2_Si ([M + H]^+^): 419.3345, found: 419.3338.

##### *N*-(2-(2-(2-(2-azidoethoxy)ethoxy)ethoxy)ethyl)-3β-(*tert*-Butyldimethylsilyloxy)-androst-5-en-17β-carboxamide (CRG007)

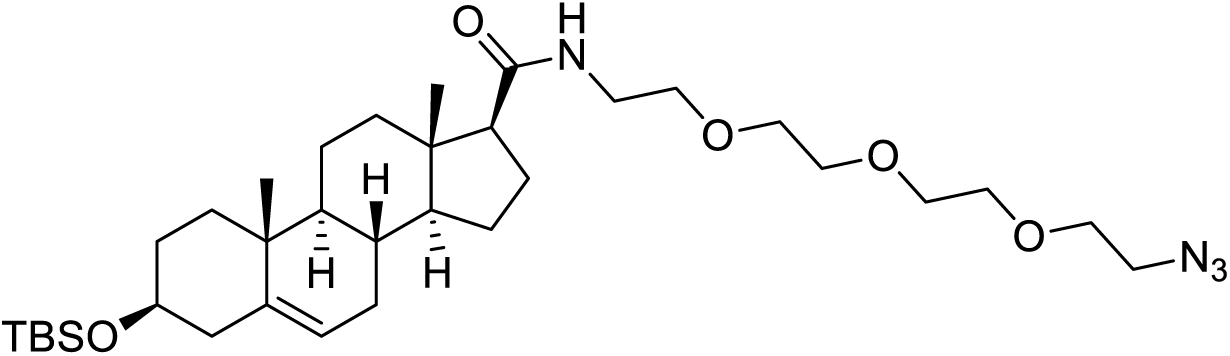

Compound **CRG007** (colourless oil, 382 mg, 80%) was obtained from **CRG006** (325 mg, 0.751 mmol), HOBt (122 mg, 0.901 mmol), EDC (173 mg, 0.901 mmol), **CRG003** (180 mg, 0.826 mmol) and TEA (157 µL, 1.13 mmol), according to general procedure 4. The title compound was purified by flash chromatography on silica gel (from 0 to 80% EtOAc in hexane).

[α]^20^_D_ = –32.9 (*c* 1.0, CHCl_3_). ^1^H NMR (400 MHz, CDCl_3_) δ 5.77 (br s, 1H), 5.31 (dd, *J* = 3.5, 1.8 Hz, 1H), 3.76 – 3.58 (m, 10H), 3.57 – 3.41 (m, 5H), 3.39 (t, *J* = 4.0 Hz, 2H), 2.31 – 0.91 (m, 20H), 1.00 (s, 3H), 0.88 (s, 9H), 0.68 (s, 3H), 0.05 (s, 6H). ^13^C NMR (101 MHz, CDCl_3_) δ 172.8, 141.6, 121.0, 72.7, 70.9, 70.8, 70.7, 70.3, 70.3, 70.2, 57.1, 56.6, 50.8, 50.3, 43.8, 42.9, 39.1, 38.5, 37.5, 36.8, 32.2, 32.1, 32.0, 26.1, 24.7, 23.6, 21.1, 19.6, 18.4, 13.2, −4.46, −4.46. HRMS calcd. for C_34_H_61_N_4_O_5_Si ([M + H]^+^): 633.4411, found: 633.4409.

##### *N*-(23-azido-3,6,9,12,15,18,21-heptaoxatricosyl)-3β-(*tert*-Butyldimethylsilyloxy)-androst-5-en-17β-carboxamide (CRG043)

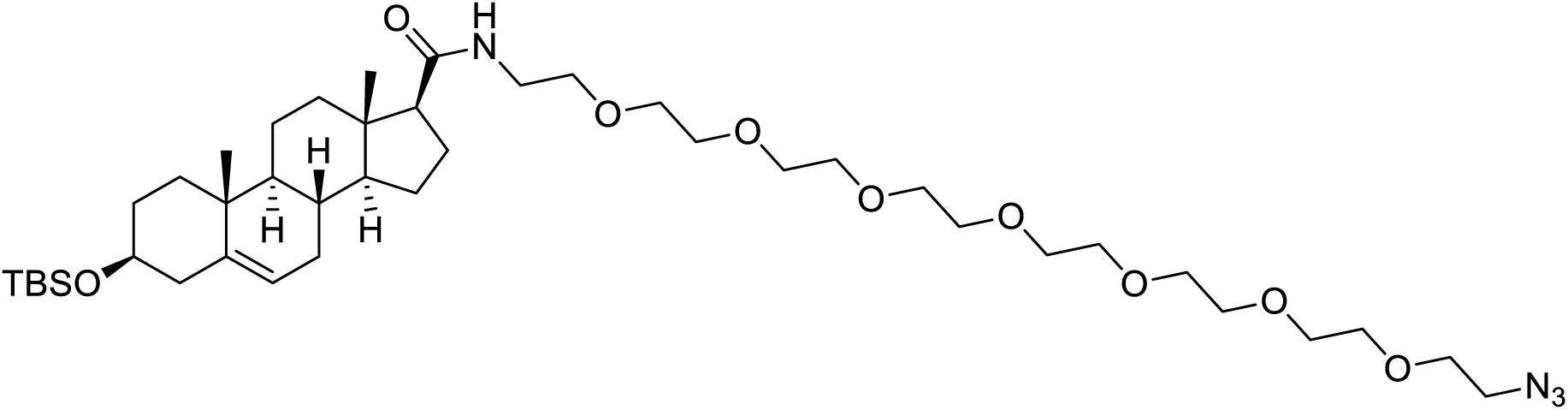

Compound **CRG043** (colourless oil, 274 mg, 92%) was obtained from **CRG006** (160 mg, 0.370 mmol), HOBt (60 mg, 0.444 mmol), EDC (85 mg, 0.444 mmol), **CRG042** (160 mg, 0.407 mmol) and TEA (77 µL, 0.555 mmol), according to general procedure 4. The title compound was purified by flash chromatography on silica gel (from 0 to 2.4% MeOH in CH_2_Cl_2_).

[α]^20^_D_ = –27.2 (*c* 1.0, CHCl_3_). ^1^H NMR (400 MHz, CDCl_3_) δ 5.84 (br s, 1H), 5.31 (dd, *J* = 3.4, 1.8 Hz, 1H), 3.73 – 3.58 (m, 25H), 3.58 – 3.39 (m, 6H), 3.38 (t, *J* = 4.0 Hz, 2H), 2.32 – 0.91 (m, 20H), 0.99 (s, 3H), 0.88 (s, 9H), 0.68 (s, 3H), 0.05 (s, 6H). ^13^C NMR (101 MHz, CDCl_3_) δ 172.8, 141.6, 121.0, 72.6, 70.8, 70.8, 70.7, 70.7, 70.6, 70.6, 70.3, 70.3, 70.1, 57.1, 56.5, 50.8, 50.3, 43.8, 42.9, 39.1, 38.5, 37.5, 36.7, 32.1, 32.0, 32.0, 26.0, 24.7, 23.6, 21.1, 19.5, 18.3, 13.2, −4.5, −4.5. HRMS calcd. for C_42_H_77_N_4_O_9_Si ([M + H]^+^): 809.5460, found: 809.5480.

##### *N*-(2-(2-(2-(2-azidoethoxy)ethoxy)ethoxy)ethyl)-3β-hydroxy-androst-5-en-17β-carboxamide (CRG008)

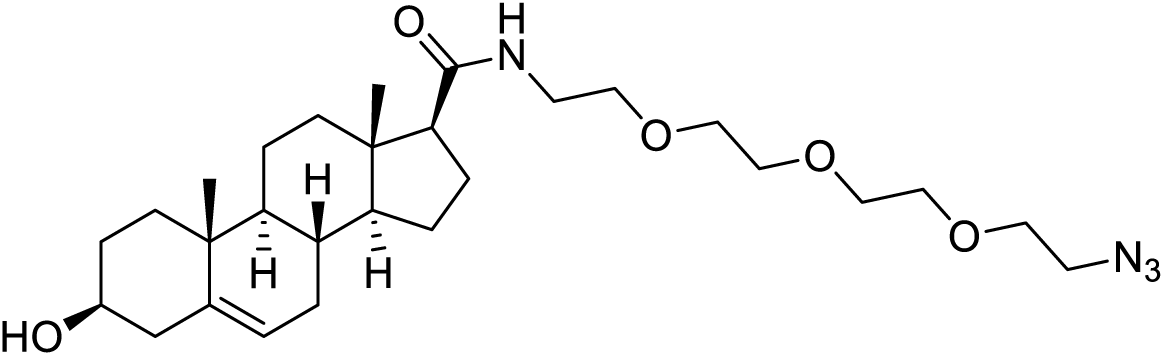

Compound **CRG008** (colourless oil, 279 mg, 92%) was obtained from **CRG007** (372 mg, 0.588 mmol) and TBAF (1.18 mL, 1.18 mmol), according to general procedure 5. The title compound was purified by flash chromatography on silica gel (from 0 to 100% EtOAc in hexane).

[α]^20^_D_ = –45.0 (*c* 1.0, CHCl_3_). ^1^H NMR (400 MHz, CDCl_3_) δ 5.79 (br s, 1H), 5.35 (app d, *J* = 5.1 Hz, 1H), 3.77 – 3.58 (m, 10H), 3.58 – 3.41 (m, 5H), 3.39 (t, *J* = 4.0 Hz, 2H), 2.34 – 0.92 (m, 20H), 1.01 (s, 3H), 0.69 (s, 3H). ^13^C NMR (101 MHz, CDCl_3_) δ 172.8, 140.9, 121.4, 71.7, 70.8, 70.7, 70.7, 70.3, 70.2, 70.2, 57.1, 56.5, 50.7, 50.2, 43.8, 42.3, 39.1, 38.4, 37.4, 36.6, 32.0, 31.9, 31.7, 24.6, 23.6, 21.1, 19.5, 13.2. HRMS calcd. for C_28_H_47_N_4_O_5_ ([M + H]^+^): 519.3546, found: 519.3559.

##### *N*-(23-azido-3,6,9,12,15,18,21-heptaoxatricosyl)-3β-hydroxy-androst-5-en-17β-carboxamide (CRG044)

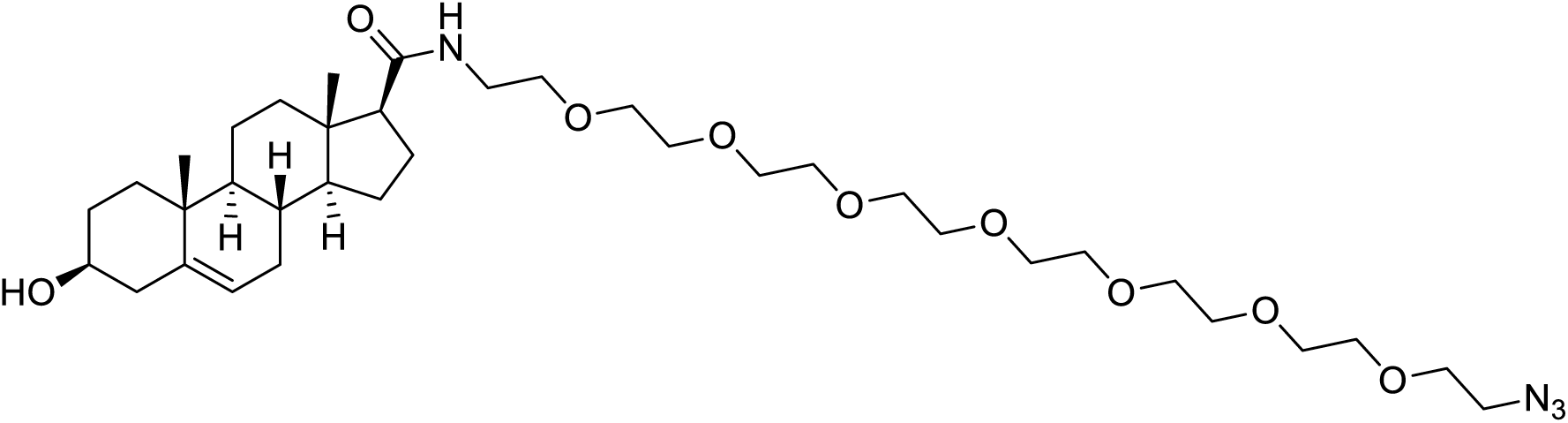

Compound **CRG044** (colourless oil, 213 mg, 97%) was obtained from **CRG043** (255 mg, 0.315 mmol) and TBAF (630 µL, 0.630 mmol), according to general procedure 5. The title compound was purified by flash chromatography on silica gel (from 0 to 5% MeOH in CH_2_Cl_2_).

[α]^20^ = –33.1 (*c* 0.9, CHCl_3_). ^1^H NMR (400 MHz, CDCl_3_) δ 5.86 (br s, 1H), 5.35 (dd, *J* = 3.6, 1.7 Hz, 1H), 3.73 – 3.40 (m, 31H), 3.38 (t, *J* = 4.0 Hz, 2H), 2.34 – 0.92 (m, 20H), 1.01 (s, 3H), 0.68 (s, 3H). ^13^C NMR (101 MHz, CDCl_3_) δ 172.9, 140.9, 121.5, 71.7, 70.8, 70.8, 70.7, 70.7, 70.6, 70.6, 70.3, 70.2, 70.1, 57.1, 56.5, 50.8, 50.2, 43.8, 42.3, 39.1, 38.5, 37.4, 36.7, 32.0, 31.9, 31.7, 24.7, 23.6, 21.1, 19.5, 13.2. HRMS calcd. for C_36_H_63_N_4_O_9_ ([M + H]^+^): 695.4595, found: 695.4606.

##### *N*-(2-(2-(2-(2-azidoethoxy)ethoxy)ethoxy)ethyl)-3-oxo-androst-4-en-17β-carboxamide (CRG009)

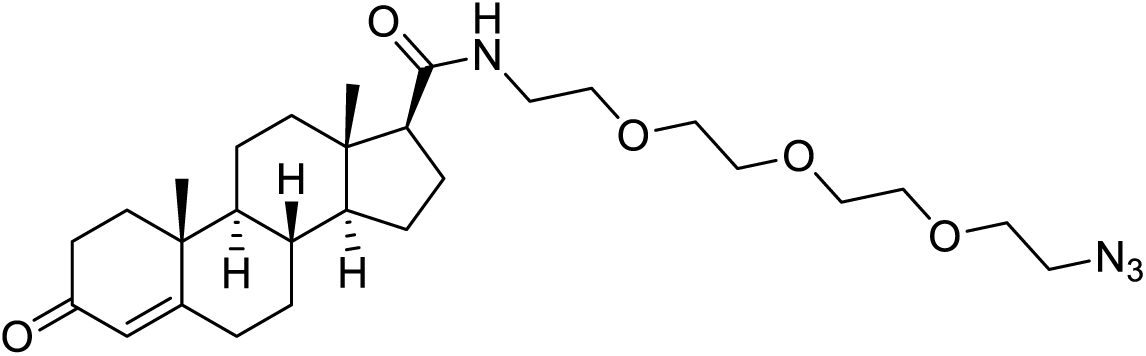

Compound **CRG009** (colourless oil, 162 mg, 63%) was obtained from **CRG008** (260 mg, 0.501 mmol) and Al(O*^t^*Bu)_3_ (185 mg, 0.752 mmol), according to general procedure 6. The title compound was purified by flash chromatography on silica gel (from 0 to 1.8% MeOH in CH_2_Cl_2_).

[α]^20^_D_ = +71.8 (*c* 0.6, CHCl_3_). ^1^H NMR (400 MHz, CDCl_3_) δ 5.79 (br s, 1H), 5.73 (s, 1H), 3.76 – 3.58 (m, 10H), 3.57 – 3.40 (m, 4H), 3.38 (t, *J* = 4.0 Hz, 2H), 2.49 – 0.91 (m, 20H), 1.19 (s, 3H), 0.73 (s, 3H). ^13^C NMR (101 MHz, CDCl_3_) δ 199.5, 172.6, 171.1, 124.0, 70.8, 70.7, 70.7, 70.3, 70.2, 70.2, 57.0, 55.6, 53.9, 50.7, 43.7, 39.1, 38.7, 38.3, 35.8, 35.7, 34.0, 32.9, 32.0, 24.5, 23.6, 21.0, 17.5, 13.3. HRMS calcd. for C_28_H_45_N_4_O_5_ ([M + H]^+^): 517.3390, found: 517.3397.

##### *N*-(23-azido-3,6,9,12,15,18,21-heptaoxatricosyl)-3-oxo-androst-4-en-17β-carboxamide (CRG045)

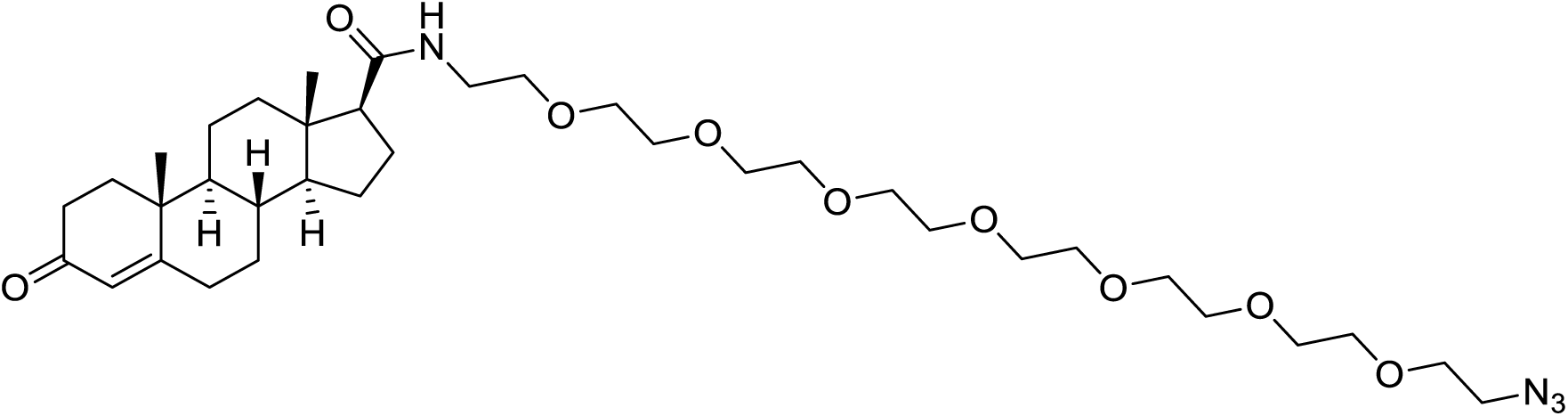

Compound **CRG045** (colourless oil, 98 mg, 58%) was obtained from **CRG044** (170 mg, 0.245 mmol) and Al(O*^t^*Bu)_3_ (90 mg, 0.367 mmol), according to general procedure 6. The title compound was purified by flash chromatography on silica gel (from 0 to 3% MeOH in CH_2_Cl_2_).

[α]^20^_D_ = +53.2 (*c* 1.0, CHCl_3_). ^1^H NMR (400 MHz, CDCl_3_) δ 5.89 (br s, 1H), 5.73 (s, 1H), 3.71 – 3.58 (m, 25H), 3.58 – 3.41 (m, 5H), 3.38 (t, *J* = 4.0 Hz, 2H), 2.49 – 0.91 (m, 20H), 1.18 (s, 3H), 0.72 (s, 3H). ^13^C NMR (101 MHz, CDCl_3_) δ 199.6, 172.7, 171.2, 124.0, 70.8, 70,8, 70.7, 70.7, 70,7, 70,7, 70.6, 70.3, 70.3, 70.2, 57.0, 55.6, 53.9, 50.8, 43.8, 39.2, 38.7, 38.3, 35.9, 35.8, 34.1, 32.9, 32.1, 24.6, 23.6, 21.1, 17.5, 13.4. HRMS calcd. for C_36_H_61_N_4_O_9_ ([M + H]^+^): 693.4439, found: 693.4421.

##### 3β-(*tert*-Butyldimethylsilyloxy)-17β-(13-azido-2,5,8,11-tetraoxatridecyl)-androst-5-ene (CRG014)

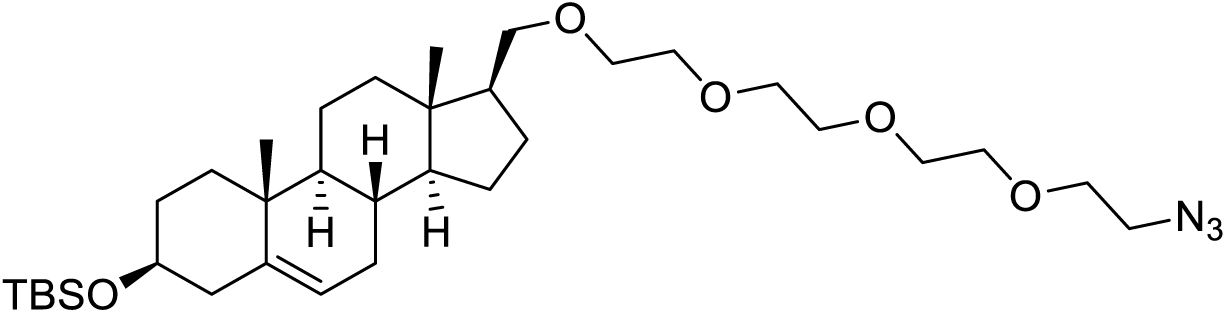

Compound **CRG014** (colourless oil, 37 mg, 24%) was obtained from **CRG013** (104 mg, 0.248 mmol), **CRG012** (74 mg, 0.248 mmol) and NaH (20 mg, 0.497 mmol), according to general procedure 7. The title compound was purified by flash chromatography on silica gel (from 0 to 25% EtOAc in hexane).

[α]^20^_D_ = –27.3 (*c* 1.0, CHCl_3_). ^1^H NMR (400 MHz, CDCl_3_) δ 5.31 (dd, *J* = 3.2, 2.0 Hz, 1H), 3.70 – 3.43 (m, 16H), 3.39 (t, *J* = 4.0 Hz, 2H), 3.33 (dd, *J* = 9.4, 7.4 Hz, 1H), 2.32 – 0.90 (m, 20H), 1.00 (s, 3H), 0.89 (s, 9H), 0.63 (s, 3H), 0.05 (s, 6H). ^13^C NMR (101 MHz, CDCl_3_) δ 141.7, 121.2, 73.2, 72.8, 70.9, 70.8, 70.8, 70.5, 70.2, 56.3, 50.8, 50.6, 49.9, 43.0, 41.8, 38.5, 37.6, 36.8, 32.2, 32.1, 31.8, 26.2, 26.1, 24.9, 20.9, 19.6, 18.4, 12.5, −4.4. HRMS calcd. for C_34_H_61_N_3_O_5_SiNa ([M + Na]^+^): 642.4278, found: 642.4277.

##### 3β-(*tert*-Butyldimethylsilyloxy)-17β-(25-azido-2,5,8,11,14,17,20,23-octaoxapentacosyl)- androst-5-ene (CRG038)

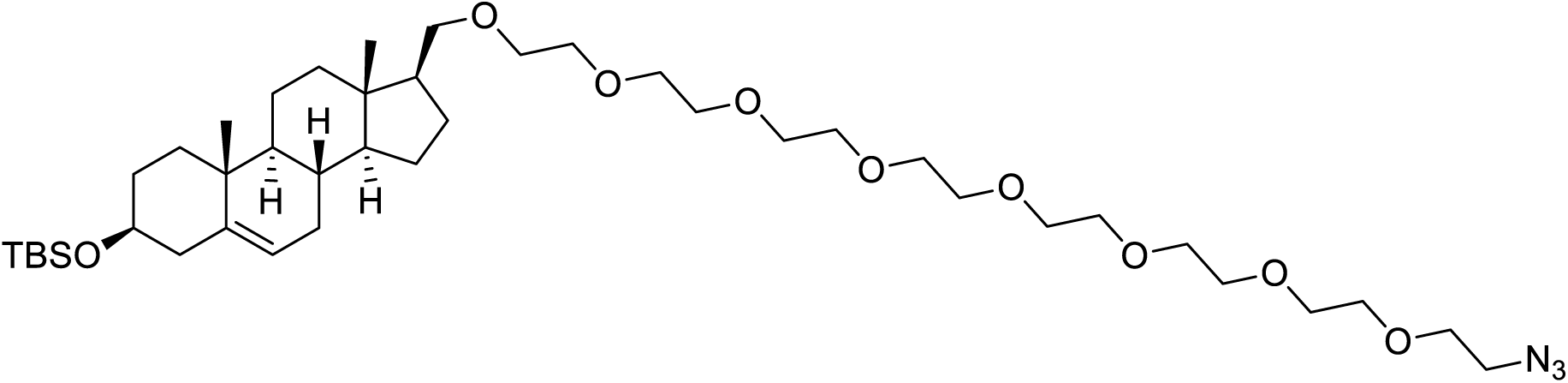

Compound **CRG038** (colourless oil, 135 mg, 53%) was obtained from **CRG013** (135 mg, 0.322 mmol), **CRG037** (177 mg, 0.322 mmol) and NaH (26 mg, 0.645 mmol), according to general procedure 7. The title compound was purified by flash chromatography on silica gel (from 0 to 100% EtOAc in hexane).

[α]^20^_D_ = –22.3 (*c* 0.6, CHCl_3_). ^1^H NMR (400 MHz, CDCl_3_) δ 5.31 (dd, *J* = 3.2, 2.0 Hz, 1H), 3.70 – 3.42 (m, 32H), 3.39 (t, *J* = 4.0 Hz, 2H), 3.33 (dd, *J* = 9.4, 7.4 Hz, 1H), 2.31 – 0.89 (m, 20H), 1.00 (s, 3H), 0.88 (s, 9H), 0.63 (s, 3H), 0.05 (s, 6H). ^13^C NMR (101 MHz, CDCl_3_) δ 141.7, 121.2, 73.2, 72.7, 70.8, 70.8, 70.8, 70.8, 70.7, 70.7, 70.4, 70.2, 56.3, 50.8, 50.6, 49.9, 43.0, 41.8, 38.5, 37.5, 36.8, 32.2, 32.1, 31.8, 26.1, 26.1, 24.9, 20.9, 19.6, 18.4, 12.5, −4.5. HRMS calcd. for C_42_H_77_N_3_O_9_SiNa ([M + Na]^+^): 818.5327, found: 818.5339.

##### 17β-(13-azido-2,5,8,11-tetraoxatridecyl)-androst-5-en-3β-ol (CRG015)

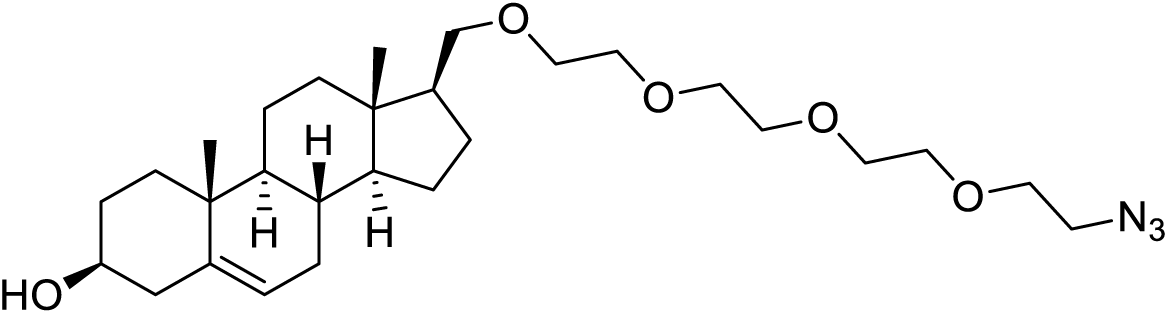

Compound **CRG015** (colourless oil, 81 mg, 86%) was obtained from **CRG014** (115 mg, 0.186 mmol) and TBAF (371 µL, 0.371 mmol), according to general procedure 5. The title compound was purified by flash chromatography on silica gel (from 0 to 45% EtOAc in hexane).

[α]^20^_D_ = –35.2 (*c* 1.0, CHCl_3_). ^1^H NMR (400 MHz, CDCl_3_) δ 5.35 (dd, *J* = 3.3, 1.9 Hz, 1H), 3.74 – 3.44 (m, 16H), 3.39 (t, *J* = 5.1 Hz, 2H), 3.33 (dd, *J* = 9.4, 7.4 Hz, 1H), 2.34 – 0.78 (m, 20H), 1.01 (s, 3H), 0.63 (s, 3H). ^13^C NMR (101 MHz, CDCl_3_) δ 140.9, 121.7, 73.2, 71.8, 70.8, 70.8, 70.7, 70.4, 70.2, 56.3, 50.8, 50.5, 49.9, 42.4, 41.8, 38.5, 37.4, 36.7, 32.1, 31.8, 31.8, 26.1, 24.8, 20.9, 19.5, 12.5. HRMS calcd. for C_28_H_47_N_3_O_5_Na ([M + Na]^+^): 528.3413, found: 528.3419.

##### 17β-(25-azido-2,5,8,11,14,17,20,23-octaoxapentacosyl)-androst-5-en-3β-ol (CRG039)

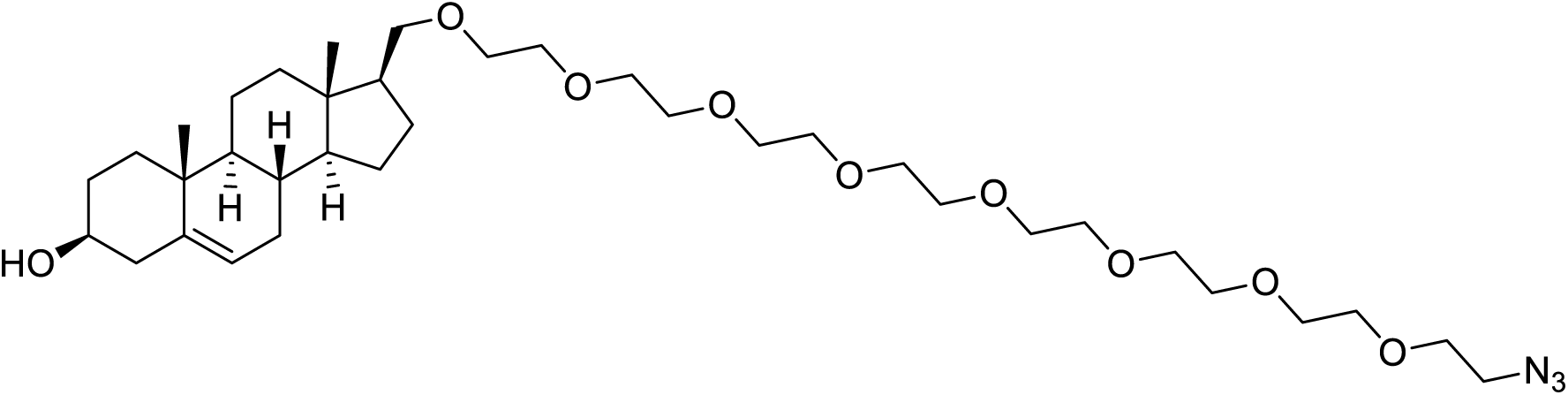

Compound **CRG039** (colourless oil, 53 mg, 76%) was obtained from **CRG038** (82 mg, 0.103 mmol) and TBAF (206 µL, 0.206 mmol), according to general procedure 5. The title compound was purified by flash chromatography on silica gel (from 0 to 4% MeOH in CH_2_Cl_2_).

[α]^20^_D_ = –23.9 (*c* 0.7, CHCl_3_). ^1^H NMR (400 MHz, CDCl_3_) δ 5.33 (dd, *J* = 3.2, 2.0 Hz, 1H), 3.85 – 3.40 (m, 32H), 3.37 (t, *J* = 4.0 Hz, 2H), 3.32 (dd, *J* = 9.4, 7.4 Hz, 1H), 2.31 – 0.90 (m, 20H), 0.99 (s, 3H), 0.62 (s, 3H). ^13^C NMR (101 MHz, CDCl_3_) δ 140.9, 121.7, 73.2, 71.8, 70.8, 70.8, 70.7, 70.7, 70.7, 70.7, 70.7, 70.4, 70.1, 56.3, 50.8, 50.5, 49.9, 42.4, 41.8, 38.4, 37.4, 36.7, 32.1, 31.8, 31.8, 26.1, 24.8, 20.9, 19.5, 12.5. HRMS calcd. for C_36_H_64_N_3_O_9_ ([M + H]^+^): 682.4643, found: 682.4656.

##### 17β-(13-azido-2,5,8,11-tetraoxatridecyl)-androst-4-en-3-one (CRG016)

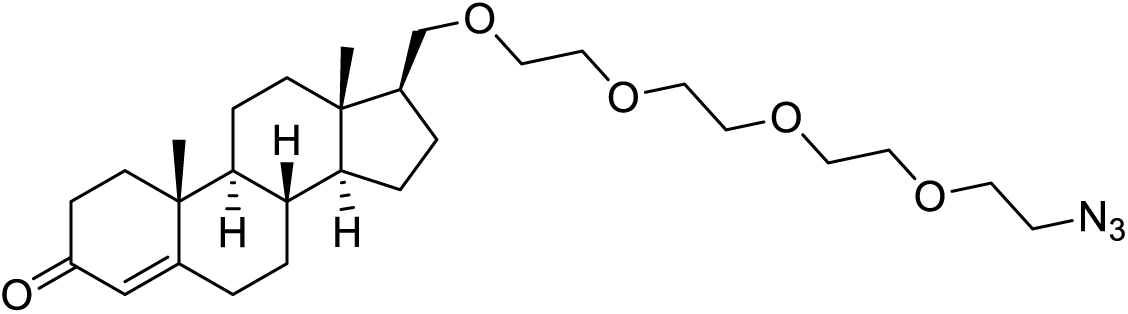

Compound **CRG016** (colourless oil, 62 mg, 89%) was obtained from **CRG015** (70 mg, 0.138 mmol) and Al(O*^t^*Bu)_3_ (51 mg, 0.208 mmol), according to general procedure 6. The title compound was purified by flash chromatography on silica gel (from 0 to 1% MeOH in CH_2_Cl_2_).

[α]^20^_D_ = +66.4 (*c* 1.0, CHCl_3_). ^1^H NMR (400 MHz, CDCl_3_) δ 5.72 (s, 1H), 3.73 – 3.47 (m, 15H), 3.39 (t, *J* = 4.0 Hz, 2H), 3.34 (dd, *J* = 9.4, 7.2 Hz, 1H), 2.48 – 0.90 (m, 20H), 1.18 (s, 3H), 0.67 (s, 3H). ^13^C NMR (101 MHz, CDCl_3_) δ 199.7, 171.6, 123.9, 73.1, 70.9, 70.8, 70.8, 70.5, 70.2, 55.4, 54.2, 50.8, 49.8, 41.9, 38.8, 38.4, 35.8, 35.5, 34.1, 33.1, 32.2, 26.0, 24.7, 20.9, 17.5, 12.5. HRMS calcd. for C_28_H_46_N_3_O_5_ ([M + H]^+^): 504.3437, found: 504.3435.

##### 17β-(25-azido-2,5,8,11,14,17,20,23-octaoxapentacosyl)-androst-4-en-3-one (CRG046)

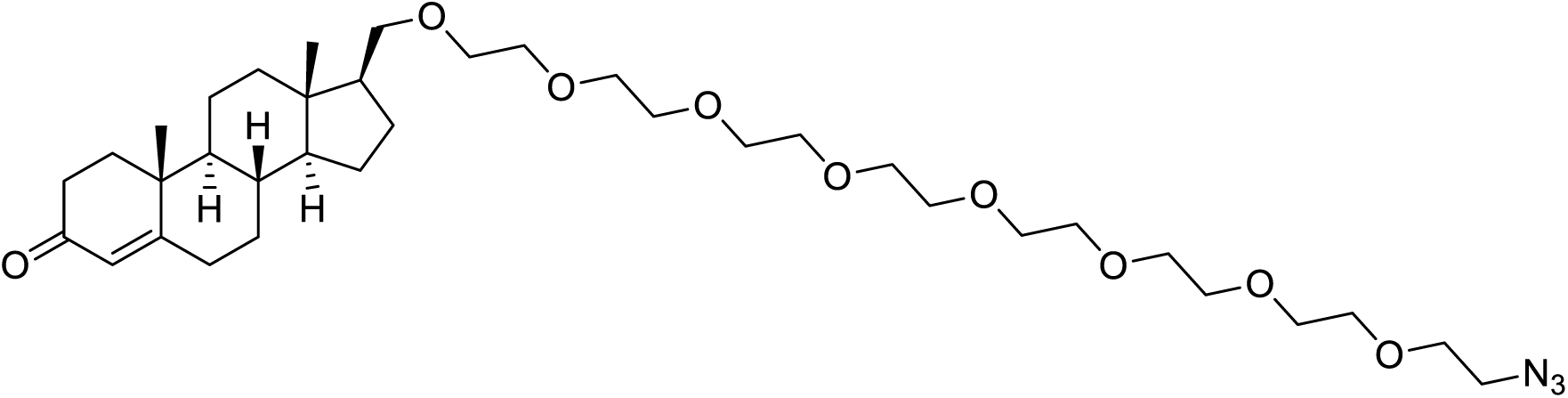

Compound **CRG046** (colourless oil, 75 mg, 92%) was obtained from **CRG039** (82 mg, 0.120 mmol) and Al(O*^t^*Bu)_3_ (44 mg, 0.180 mmol), according to general procedure 6. The title compound was purified by flash chromatography on silica gel (from 0 to 2% MeOH in CH_2_Cl_2_).

[α]^20^_D_ = +53.3 (*c* 0.7, CHCl_3_). ^1^H NMR (400 MHz, CDCl_3_) δ 5.72 (s, 1H), 3.85 – 3.44 (m, 31H), 3.38 (t, *J* = 4.0 Hz, 2H), 3.34 (dd, *J* = 9.4, 7.2 Hz, 1H), 2.48 – 0.90 (m, 20H), 1.18 (s, 3H), 0.66 (s, 3H). ^13^C NMR (101 MHz, CDCl_3_) δ 199.7, 171.6, 123.9, 73.1, 70.8, 70.8, 70.7, 70.7, 70.7, 70.7, 70.7, 70.4, 70.1, 55.4, 54.2, 50.8, 49.8, 41.9, 38.8, 38.3, 35.8, 35.5, 34.1, 33.0, 32.2, 25.9, 24.7, 20.9, 17.5, 12.5. HRMS calcd. for C_36_H_61_N_3_O_9_Na ([M + H]^+^): 702.4306, found: 702.4361.

##### (*R*)-Glycidol trityl ether (CRG023)

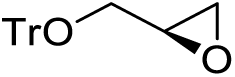

(*S*)-(−)-Glycidol (250 mg, 3.37 mmol) was added dropwise to a stirred solution of trityl chloride (1.03 g, 3.71 mmol) and TEA (517 µL, 3.71 mmol) in CH_2_Cl_2_ (3.5 mL) at 0 °C. After the addition was complete, the mixture was allowed to warm to rt and was stirred overnight. The reaction mixture was then poured into saturated aq. NH_4_Cl (10 mL) and the resulting mixture was extracted with CH_2_Cl_2_. The organic extracts were washed with brine, dried over anhydrous MgSO_4_, filtered and concentrated under reduced pressure. Flash chromatography of the residue (from 0 to 5% EtOAc in hexane) gave **CRG023** as a white solid (986 mg, 92%).

[α]^20^_D_ = +11.0 (*c* 1.0, CHCl_3_) [lit.^5^ [α]^23^_D_ = +11.2 (*c* 1.0, CHCl_3_]. ^1^H NMR (400 MHz, CDCl_3_) δ 7.50 – 7.42 (m, 6H), 7.33 – 7.21 (m, 9H), 3.37 – 3.28 (m, 1H), 3.19 – 3.09 (m, 2H), 2.81 – 2.75 (m, 1H), 2.63 (dd, *J* = 5.1, 2.3 Hz, 1H). ^13^C NMR (101 MHz, CDCl_3_) δ 143.9, 128.8, 128.0, 127.2, 86.8, 64.9, 51.2, 44.8. HRMS calcd. for C_22_H_20_O_2_Na ([M + Na]^+^): 339.1361, found: 339.1346.

##### (*R*)-1-(hexadecyloxy)-3-(trityloxy)propan-2-ol (CRG024)

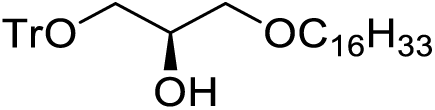

To a solution of **CRG023** (640 mg, 2.02 mmol) in DMF (15 mL) were added portionwise KO*^t^*Bu (681 mg, 6.07 mmol) and hexadecanol (736 mg, 3.03 mmol) at rt. The resulting heterogeneous mixture was stirred at 80 °C for 1h. After cooling down to rt, water (15 mL) was added and the resulting mixture was extracted with Et_2_O. The combined organic layers were washed with brine, dried over anhydrous MgSO_4_, filtered and evaporated to dryness. Purification of the residue by flash chromatography (from 0 to 18% Et_2_O in hexane) afforded **CRG024** as a white solid (623 mg, 55%).

[α]^20^_D_ = +3.1 (*c* 1.0, CHCl_3_) [lit.^6^ [α]^26^_D_ = +2.3 (*c* 1.02, CHCl_3_]. ^1^H NMR (400 MHz, CDCl_3_) δ 7.46 – 7.41 (m, 6H), 7.33 – 7.20 (m, 9H), 3.99 – 3.90 (m, 1H), 3.54 (dd, *J* = 9.7, 4.3 Hz, 1H), 3.50 – 3.38 (m, 3H), 3.25 – 3.14 (m, 2H), 2.41 (d, *J* = 4.6 Hz, 1H), 1.58 – 1.49 (m, 2H), 1.37 – 1.16 (m, 26H), 0.88 (t, *J* = 6.8 Hz, 3H). ^13^C NMR (101 MHz, CDCl_3_) δ 144.0, 128.8, 128.0, 127.2, 86.8, 72.2, 71.8, 70.0, 64.8, 32.1, 29.9, 29.8, 29.8, 29.8, 29.8, 29.7, 29.5, 26.3, 22.9, 14.3. HRMS calcd. for C_38_H_54_O_3_Na ([M + Na]^+^): 581.3971, found: 581.3977.

##### (*R*)-9-(diethylamino)-2-((8-((1-(hexadecyloxy)-3-(trityloxy)propan-2-yl)oxy)octyl)oxy)-5*H*-benzo[*a*]phenoxazin-5-one (CRG029)

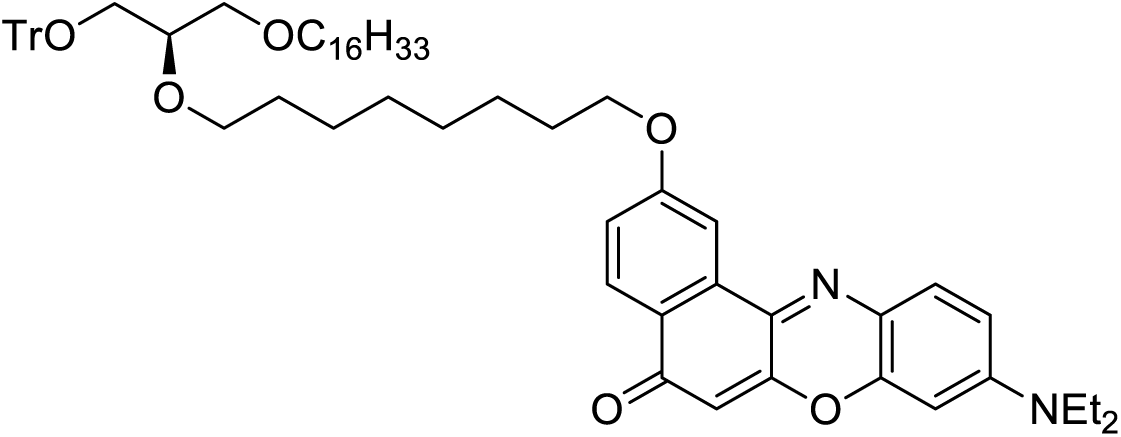

To a solution of **CRG024** (300 mg, 0.537 mmol) in DMF (5.5 mL) was added portionwise NaH (60 wt.% in mineral oil, 64mg, 1.61 mmol) at 0 °C and the mixture was stirred at the same temperature for 10 min. 1,8-dibromooctane (494 µL, 2.68 mmol) was then added dropwise and the resulting mixture was stirred at 65 °C overnight. After cooling down to rt, the reaction mixture was carefully diluted with water and extracted with Et_2_O. The combined organic layers were washed with brine, dried over anhydrous MgSO_4_, filtered and concentrated under reduced pressure. The residue was dissolved in CH_2_Cl_2_ and was filtered through a short pad of silica gel firstly eluting with hexane in order to remove the excess of 1,8-dibromooctane and then with hexane/EtOAc (95:5), affording the corresponding bromo derivative **5** as a crude intermediate. The resulting residue was dissolved in DMF (2 mL) and was directly added to a solution of 2-hydroxy Nile Red^1^ (216 mg, 0.645 mmol) and K_2_CO_3_ (149 mg, 1.07 mmol) in DMF (4 mL) at rt. The reaction mixture was stirred overnight at 65 °C, cooled to rt and diluted with water (10 mL). The resulting dark red solution was extracted with Et_2_O (5x) and the combined organic layers were washed with brine, dried over anhydrous MgSO_4_, filtered and evaporated to dryness. Purification of the residue by flash chromatography (from 0 to 40% EtOAc in hexane) gave **CRG029** as a dark violet oil (289 mg, 54% over two steps).

^1^H NMR (400 MHz, CDCl_3_) δ 8.18 (d, *J* = 8.7 Hz, 1H), 8.01 (d, *J* = 2.6 Hz, 1H), 7.56 (d, *J* = 9.0 Hz, 1H), 7.49 – 7.37 (m, 6H), 7.30 – 7.16 (m, 9H), 7.12 (dd, *J* = 8.7, 2.6 Hz, 1H), 6.61 (dd, *J* = 9.1, 2.7 Hz, 1H), 6.42 (d, *J* = 2.7 Hz, 1H), 6.27 (s, 1H), 4.10 (t, *J* = 6.5 Hz, 2H), 3.59 – 3.34 (m, 11H), 3.20 – 3.10 (m, 2H), 1.86 – 1.77 (m, 2H), 1.63 – 1.12 (m, 44H), 0.85 (t, *J* = 6.8 Hz, 3H). ^13^C NMR (101 MHz, CDCl_3_) δ 183.4, 162.0, 152.2, 150.8, 147.0, 144.3, 140.3, 134.2, 131.2, 128.9, 127.8, 127.0, 125.6, 124.9, 118.4, 109.6, 106.8, 105.4, 96.5, 86.6, 78.5, 71.8, 71.3, 70.8, 68.5, 63.8, 45.2, 32.1, 30.3, 29.8, 29.8, 29.8, 29.7, 29.6, 29.6, 29.5, 29.4, 26.3, 26.3, 26.2, 22.8, 14.3, 12.8. HRMS calcd. for C_66_H_87_N_2_O_6_ ([M + H]^+^): 1003.6564, found: 1003.6587.

##### (*S*)-9-(diethylamino)-2-((8-((1-(hexadecyloxy)-3-hydroxypropan-2-yl)oxy)octyl)oxy)-5*H*-benzo[*a*]phenoxazin-5-one (CRG031)

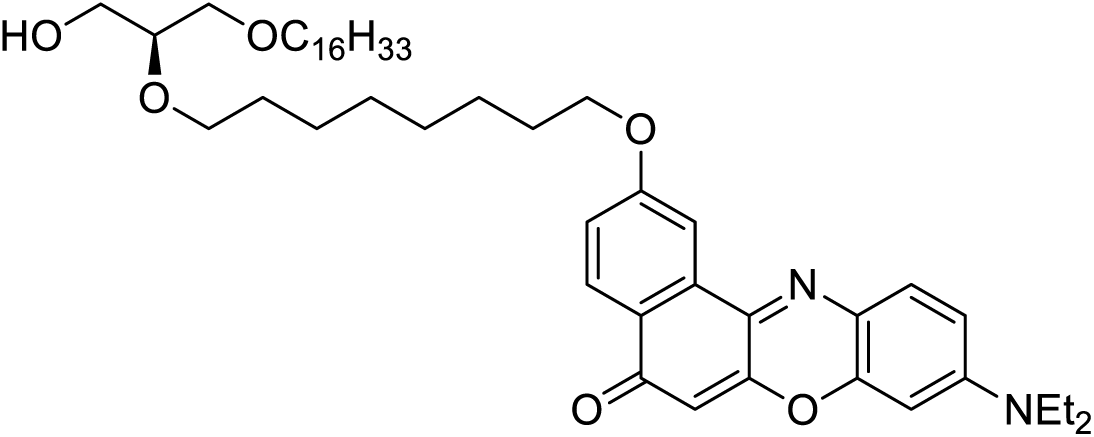

To an ice-cooled solution of **CRG029** (160 mg, 0.160 mmol) in MeOH (3 mL) and CHCl_3_ (250 µL), *p*-TsOH (55 mg, 0.319 mmol) was added in one portion. After stirring at rt for 2.5 h, the reaction was carefully quenched with saturated aqueous NaHCO_3_ (15 mL) and the resulting mixture was extracted with CH_2_Cl_2_. The combined organic extracts were washed with brine, dried over anhydrous MgSO_4_, filtered and evaporated *in vacuo*. The residue was purified by flash chromatography (from 0 to 70% EtOAc in hexane) to yield **CRG031** (110 mg, 91%) as a dark violet oil.

^1^H NMR (400 MHz, CDCl_3_) δ 8.19 (d, *J* = 8.7 Hz, 1H), 8.01 (d, *J* = 2.6 Hz, 1H), 7.57 (d, *J* = 9.0 Hz, 1H), 7.14 (dd, *J* = 8.7, 2.6 Hz, 1H), 6.62 (dd, *J* = 9.1, 2.7 Hz, 1H), 6.42 (d, *J* = 2.7 Hz, 1H), 6.27 (s, 1H), 4.15 (t, *J* = 6.5 Hz, 2H), 3.72 (dd, *J* = 11.3, 3.7 Hz, 1H), 3.66 – 3.57 (m, 2H), 3.57 – 3.37 (m, 10H), 1.90 – 1.80 (m, 2H), 1.68 – 1.09 (m, 44H), 0.87 (t, *J* = 6.9 Hz, 3H). ^13^C NMR (101 MHz, CDCl_3_) δ 183.4, 162.0, 152.1, 150.8, 146.9, 140.2, 134.2, 131.2, 127.8, 125.6, 124.8, 118.4, 109.6, 106.7, 105.4, 96.4, 78.4, 72.0, 71.0, 70.5, 68.5, 63.2, 45.2, 32.1, 30.2, 29.8, 29.8, 29.8, 29.8, 29.7, 29.6, 29.5, 29.5, 29.4, 26.2, 26.2, 26.2, 22.8, 14.3, 12.8. HRMS calcd. for C_47_H_73_N_2_O_6_ ([M + H]^+^): 761.5469, found: 761.5490.

##### (*R*)-2-((8-((9-(diethylamino)-5-oxo-5*H*-benzo[*a*]phenoxazin-2-yl)oxy)octyl)oxy)-3-(hexadecyloxy)propyl (2-(dimethyl(prop-2-yn-1-yl)ammonio)ethyl) phosphate (CRG032)

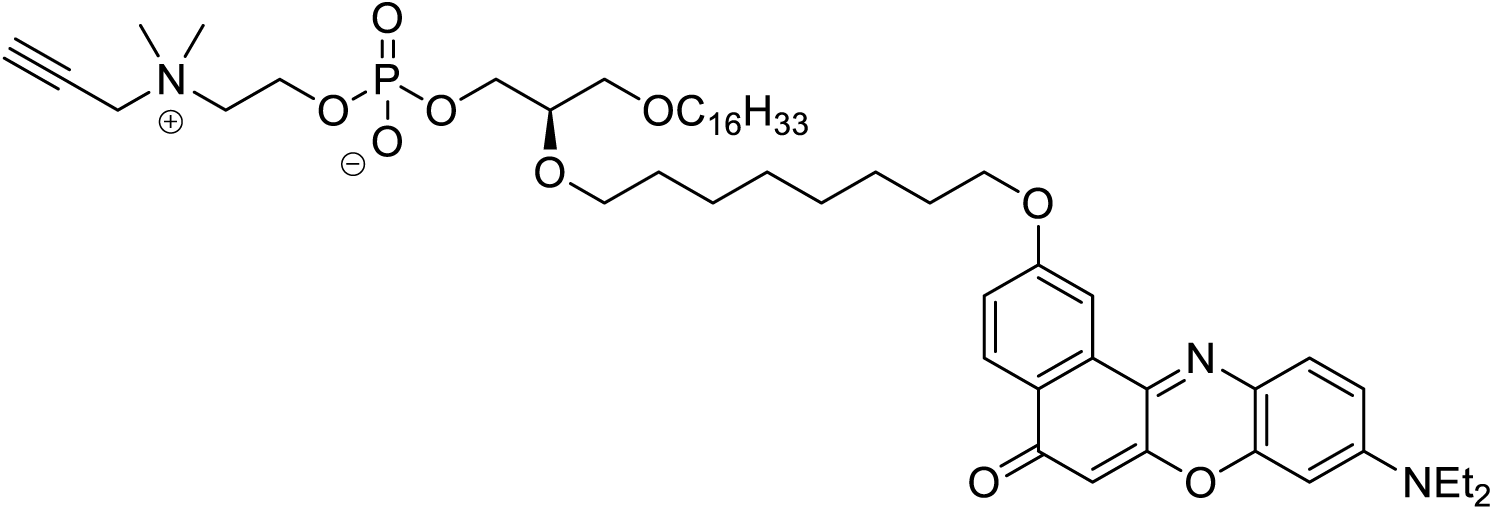

A solution of **CRG031** (110 mg, 0.145 mmol) in toluene (1.5 mL) at 0 °C was successively treated with TEA (60 µL, 0.434 mmol), DMAP (4 mg, 0.029 mmol) and 2-chloro-1,3,2-dioxaphospholane 2-oxide (40 µL, 0.434 mmol). After stirring at 0 °C for 1.5 h, the reaction mixture was transferred to a sealed tube *via* cannula under positive nitrogen pressure. The crude solution was diluted with MeCN (13 mL) before 3-dimethylamino-1-propyne (621 µL, 5.77 mmol) was added dropwise. The tube was flushed with argon, sealed and the mixture stirred at 80 °C for 48 h. The reaction mixture was then transferred to a round-bottom flask, concentrated *in vacuo* and purified by flash chromatography on silica gel (from 90:10:0 to 67:30:3 CHCl_3_/MeOH/H_2_O), yielding the zwitterionic amino phosphate **CRG032** (58 mg, 42% over two steps) as a dark violet oil.

^1^H NMR (400 MHz, CD_3_OD/CD_3_CN (1:1)) δ 8.09 (d, *J* = 8.8 Hz, 1H), 8.00 (s, 1H), 7.58 (d, *J* = 9.1 Hz, 1H), 7.18 (d, *J* = 8.8 Hz, 1H), 6.83 (d, *J* = 7.1 Hz, 1H), 6.56 (s, 1H), 6.20 (s, 1H), 4.37 (d, *J* = 2.5 Hz, 2H), 4.28 (br s, 2H), 4.17 (t, *J* = 6.4 Hz, 2H), 3.89 (td, *J* = 10.5, 5.3 Hz, 2H), 3.73 – 3.67 (m, 2H), 3.67 – 3.40 (m, 12H), 3.25 (s, 6H), 1.92 – 1.82 (m, 2H), 1.63 – 1.18 (m, 44H), 0.89 (t, *J* = 6.9 Hz, 4H). ^13^C NMR (101 MHz, CD_3_OD/CD_3_CN (1:1)) δ 184.0, 163.0, 153.5, 152.6, 147.9, 139.3, 135.2, 132.1, 128.2, 126.0, 125.9, 119.0, 111.6, 107.2, 104.8, 97.0, 83.0, 79.1 (d, *J*_C-P_ = 7.9 Hz), 72.4, 72.2, 71.4, 71.2, 69.3, 66.2 (d, *J*_C-P_ = 5.1 Hz), 65.2 (d, *J*_C-P_ = 6.8 Hz), 60.0 (d, *J*_C-P_ = 4.2 Hz), 56.3, 52.1, 46.0, 32.8, 31.0, 30.7, 30.6, 30.6, 30.6, 30.4, 30.4, 30.3, 30.1, 27.1, 27.0, 26.9, 23.5, 14.4, 12.9, 9.2. ^31^P NMR (162 MHz, CD_3_OD/CD_3_CN (1:1)) δ 0.69. HRMS calcd. for C_54_H_85_N_3_O_9_P ([M + H]^+^): 950.6023, found: 950.6133.

##### Probe CRG033

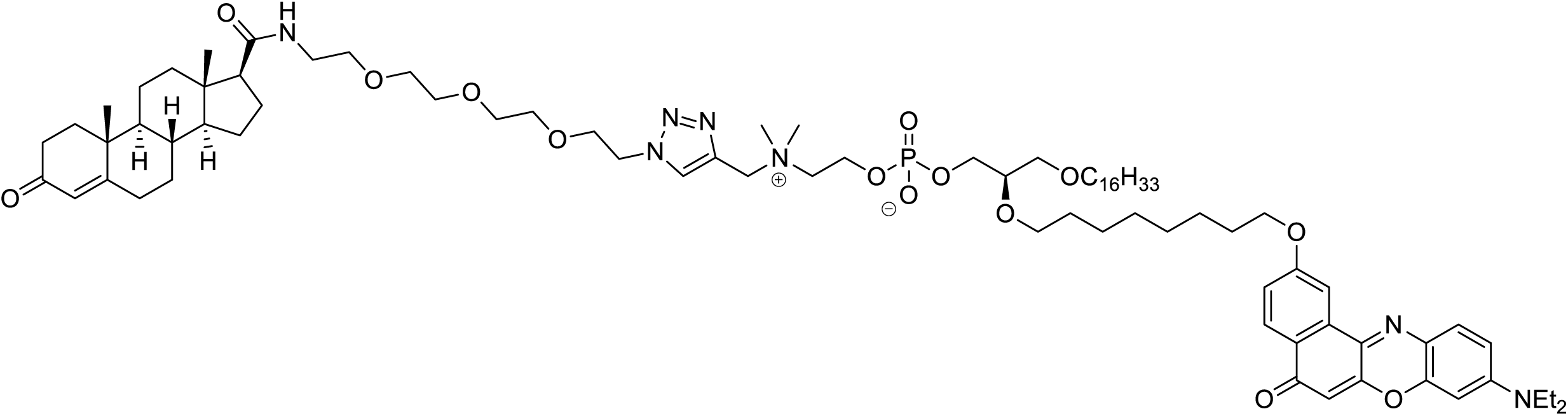

Compound **CRG033** (dark violet oil, 19 mg, 95%) was obtained from **CRG032** (13 mg, 0.014 mmol), **CRG009** (8 mg, 0.016 mmol), DIPEA (5 µL, 0.027 mmol) and CuI (0.391 mg, 0.002 mmol), according to general procedure 8. The title compound was purified by flash chromatography on silica gel (CHCl_3_/MeOH/H_2_O (78:20:2)).

^1^H NMR (400 MHz, CD_3_OD/CD_3_CN (1:1)) δ 8.32 (s, 1H), 8.06 (d, *J* = 8.8 Hz, 1H), 7.95 (d, *J* = 2.5 Hz, 1H), 7.54 (d, *J* = 9.1 Hz, 1H), 7.14 (dd, *J* = 8.8, 2.5 Hz, 1H), 7.09 – 7.02 (m, 1H), 6.79 (dd, *J* = 9.2, 2.6 Hz, 1H), 6.52 (d, *J* = 2.6 Hz, 1H), 6.16 (s, 1H), 5.66 (s, 1H), 4.73 (s, 2H), 4.63 (t, *J* = 5.1 Hz, 2H), 4.34 (br s, 2H), 4.14 (t, *J* = 6.5 Hz, 2H), 3.94 – 3.85 (m, 4H), 3.68 – 3.36 (m, 24H), 3.29 – 3.22 (m, 1H), 3.17 (s, 6H), 2.50 – 0.83 (m, 66H), 1.18 (s, 3H), 0.88 (t, *J* = 6.9 Hz, 3H), 0.68 (s, 3H). ^13^C NMR (101 MHz, CD_3_OD/CD_3_CN (1:1)) δ 201.5, 184.4, 174.7, 174.2, 163.1, 153.7, 152.6, 148.0, 139.5, 136.2, 135.3, 132.2, 130.1, 128.4, 126.1, 125.9, 124.1, 119.1, 111.6, 107.4, 105.0, 97.0, 79.1 (d, *J*_C-P_ = 7.9 Hz), 72.4, 71.5, 71.3, 71.2, 71.2, 71.0, 70.5, 69.9, 69.4, 66.1 (d, *J*_C-P_ = 5.1 Hz), 64.9 (d, *J*_C-P_ = 9.1 Hz), 60.4, 60.0 (d, *J*_C-P_ = 4.2 Hz), 57.4, 56.4, 54.9, 52.0, 51.4, 46.0, 44.9, 39.9, 39.7, 38.8, 36.5, 34.6, 33.6, 33.0, 32.9, 31.0, 30.7, 30.6, 30.6, 30.4, 30.3, 30.3, 30.3, 30.1, 27.1, 27.0, 26.9, 25.3, 24.3, 23.6, 21.8, 17.7, 17.7, 14.4, 13.8, 12.9. ^31^P NMR (162 MHz, CD_3_OD/CD_3_CN (1:1)) δ 0.35. HRMS calcd. for C_82_H_129_N_7_O_14_P ([M + H]^+^): 1466.9335, found: 1466.9370.

##### Probe CRG047

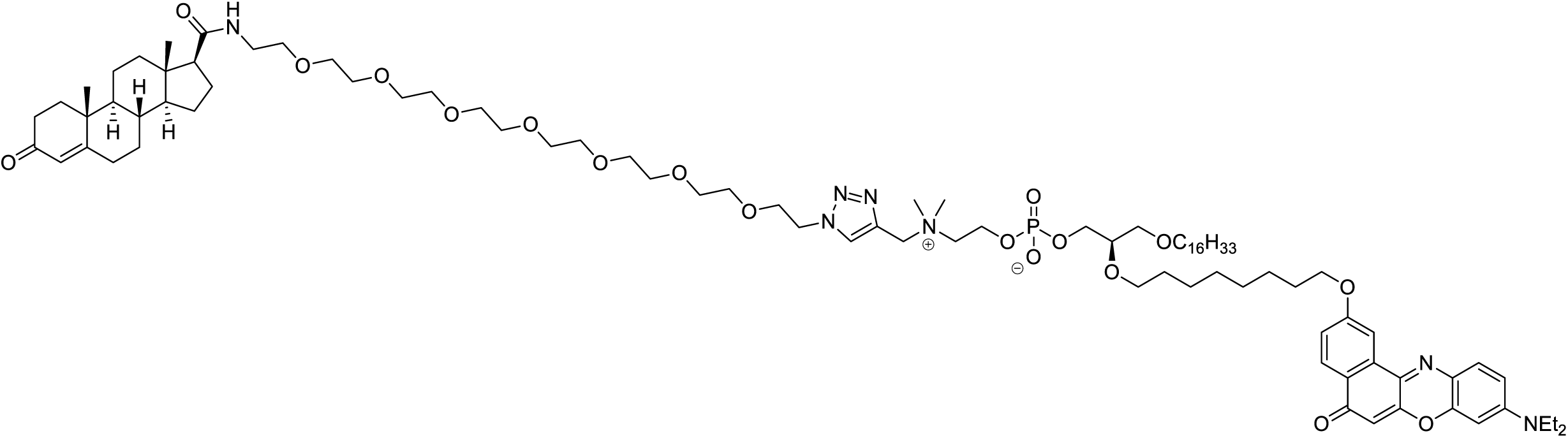

Compound **CRG047** (dark violet oil, 22 mg, 61%) was obtained from **CRG032** (25 mg, 0.026 mmol), **CRG045** (21 mg, 0.030 mmol), DIPEA (9 µL, 0.053 mmol) and CuI (0.752 mg, 0.004 mmol), according to general procedure 8. The title compound was purified by flash chromatography on silica gel (CHCl_3_/MeOH/H_2_O (78:20:2)).

^1^H NMR (400 MHz, CD_3_OD) δ 8.40 (s, 1H), 8.04 (d, *J* = 8.7 Hz, 1H), 7.94 (d, *J* = 2.4 Hz, 1H), 7.54 (d, *J* = 9.1 Hz, 1H), 7.51 – 7.45 (m, 1H), 7.12 (dd, *J* = 8.8, 2.5 Hz, 1H), 6.79 (dd, *J* = 9.2, 2.6 Hz, 1H), 6.53 (d, *J* = 2.6 Hz, 1H), 6.16 (s, 1H), 5.66 (s, 1H), 4.77 (s, 2H), 4.67 – 4.62 (m, 2H), 4.36 (br s, 2H), 4.12 (t, *J* = 6.3 Hz, 2H), 3.95 – 3.87 (m, 4H), 3.71 – 3.35 (m, 40H), 3.29 – 3.23 (m, 1H), 3.20 (s, 6H), 2.50 – 0.82 (m, 66H), 1.18 (s, 3H), 0.87 (t, *J* = 6.9 Hz, 3H), 0.68 (s, 3H). ^13^C NMR (101 MHz, CD_3_OD) δ 202.0, 184.9, 175.2, 174.8, 163.4, 153.9, 152.9, 148.3, 139.5, 136.5, 135.5, 132.5, 130.5, 128.5, 126.3, 124.2, 119.3, 111.8, 107.5, 105.0, 97.2, 79.4 (d, *J*_C-P_ = 4.0 Hz), 72.6, 71.8, 71.6, 71.6, 71.5, 71.5, 71.4, 71.4, 71.2, 70.8, 70.2, 69.5, 66.5 (d, *J*_C-P_ = 5.1 Hz), 65.0 (d, *J*_C-P_ = 5.1 Hz), 60.7, 60.3 (d, *J*_C-P_ = 3.0 Hz), 57.6, 56.7, 55.3, 52.0, 51.6, 46.2, 45.1, 40.3, 39.9, 39.0, 36.8, 36.8, 34.7, 33.9, 33.2, 33.1, 31.2, 30.9, 30.8, 30.8, 30.8, 30.7, 30.6, 30.5, 30.4, 27.3, 27.2, 27.2, 25.5, 24.5, 23.8, 22.0, 17.7, 14.5, 13.9, 13.0. ^31^P NMR (162 MHz, CD_3_OD) δ 0.69. HRMS calcd. for C_90_H_146_N_7_O_18_P ([M + 2H]^2+^): 822.0231, found: 822.0209.

##### Probe CRG034

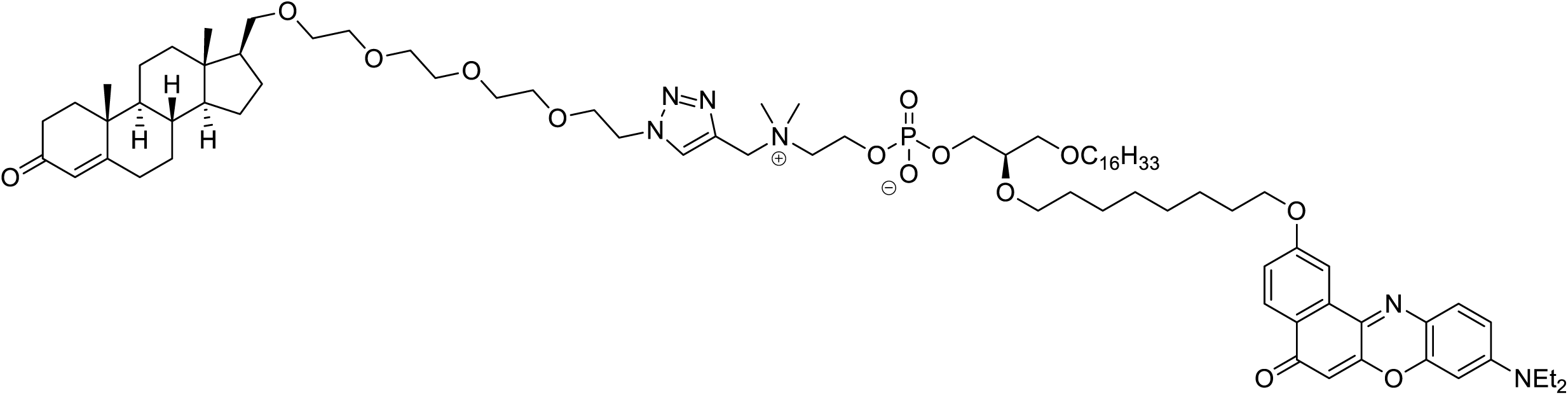

Compound **CRG034** (dark violet oil, 15 mg, 75%) was obtained from **CRG032** (13 mg, 0.014 mmol), **CRG016** (8 mg, 0.016 mmol), DIPEA (5 µL, 0.027 mmol) and CuI (0.391 mg, 0.002 mmol), according to general procedure 8. The title compound was purified by flash chromatography on silica gel (CHCl_3_/MeOH/H_2_O (83.5:15:1.5)).

^1^H NMR (400 MHz, CD_3_OD) δ 8.39 (s, 1H), 8.04 (d, *J* = 8.8 Hz, 1H), 7.93 (d, *J* = 2.5 Hz, 1H), 7.53 (d, *J* = 9.1 Hz, 1H), 7.12 (dd, *J* = 8.8, 2.6 Hz, 1H), 6.78 (dd, *J* = 9.2, 2.7 Hz, 1H), 6.52 (d, *J* = 2.6 Hz, 1H), 6.15 (s, 1H), 5.63 (s, 1H), 4.76 (s, 2H), 4.67 – 4.62 (m, 2H), 4.35 (br s, 2H), 4.12 (t, *J* = 6.4 Hz, 2H), 3.95 – 3.87 (m, 4H), 3.67 – 3.40 (m, 26H), 3.29 – 3.23 (m, 1H), 3.20 (s, 6H), 2.48 – 0.75 (m, 66H), 1.15 (s, 3H), 0.87 (t, *J* = 6.9 Hz, 3H), 0.62 (s, 3H). ^13^C NMR (101 MHz, CD_3_OD) δ 202.1, 184.8, 174.9, 163.4, 153.9, 152.9, 148.2, 139.5, 136.5, 135.5, 132.5, 130.5, 128.5, 126.3, 126.2, 124.2, 119.3, 111.8, 107.5, 105.0, 97.2, 79.4 (d, *J*_C-P_ = 8.1 Hz), 73.9, 72.6, 71.8, 71.6, 71.6, 71.5, 71.5, 71.4, 70.2, 69.5, 66.4 (d, *J*_C-P_ = 6.1 Hz), 64.9 (d, *J*_C-P_ = 8.1 Hz), 60.6, 60.3 (d, *J*_C-P_ = 5.1 Hz), 56.6, 55.5, 52.0, 51.6, 51.0, 46.2, 42.9, 39.9, 39.5, 36.7, 36.5, 34.7, 33.9, 33.3, 33.1, 31.2, 30.9, 30.9, 30.8, 30.8, 30.7, 30.7, 30.6, 30.5, 30.4, 27.4, 27.2, 27.2, 26.8, 25.5, 23.8, 21.9, 17.7, 14.5, 13.0, 12.8. ^31^P NMR (162 MHz, CD_3_OD) δ 0.72. HRMS calcd. for C_82_H_130_N_6_O_14_P ([M + H]^+^): 1453.9383, found: 1453.9371.

##### Probe CRG048

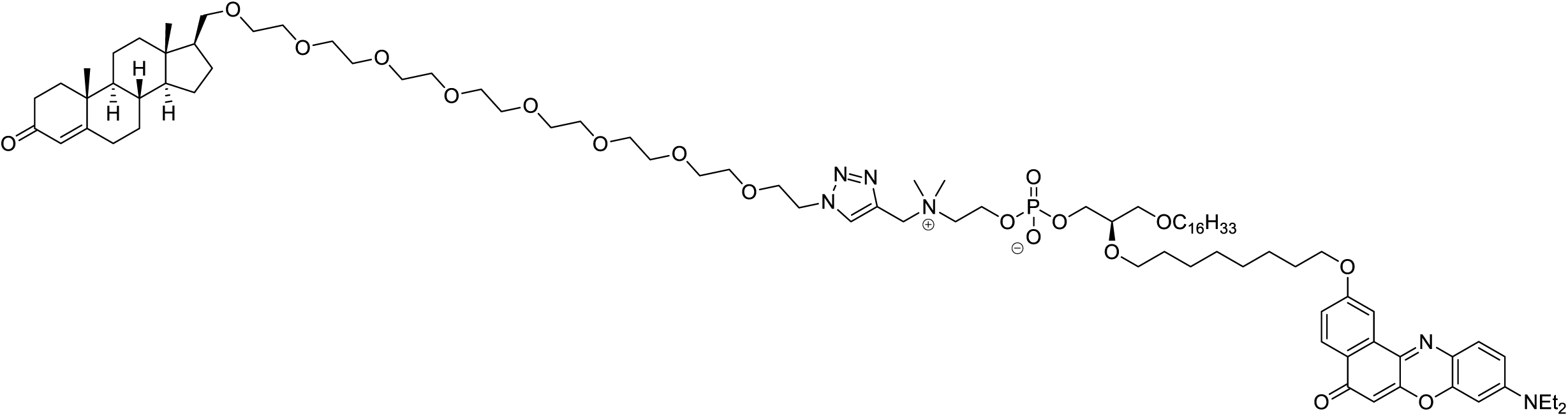

Compound **CRG048** (dark violet oil, 34 mg, 79%) was obtained from **CRG032** (25 mg, 0.026 mmol), **CRG046** (21 mg, 0.030 mmol), DIPEA (9 µL, 0.053 mmol) and CuI (0.752 mg, 0.004 mmol), according to general procedure 8. The title compound was purified by flash chromatography on silica gel (CHCl_3_/MeOH/H_2_O (78.5:20:1.5)).

^1^H NMR (400 MHz, CD_3_OD) δ 8.40 (s, 1H), 8.06 (d, *J* = 8.8 Hz, 1H), 7.98 (d, *J* = 2.5 Hz, 1H), 7.57 (d, *J* = 9.1 Hz, 1H), 7.15 (dd, *J* = 8.8, 2.6 Hz, 1H), 6.81 (dd, *J* = 9.2, 2.6 Hz, 1H), 6.56 (d, *J* = 2.6 Hz, 1H), 6.18 (s, 1H), 5.65 (s, 1H), 4.76 (s, 2H), 4.67 – 4.62 (m, 2H), 4.36 (br s, 2H), 4.14 (t, *J* = 6.4 Hz, 2H), 3.96 – 3.87 (m, 4H), 3.71 – 3.38 (m, 41H), 3.20 (s, 6H), 2.49 – 2.37 (m, 66H), 1.17 (s, 3H), 0.87 (t, *J* = 6.9 Hz, 3H), 0.65 (s, 3H). ^13^C NMR (101 MHz, CD_3_OD) δ 202.1, 184.8, 175.0, 163.4, 153.9, 152.9, 148.2, 139.5, 136.5, 135.5, 132.5, 130.5, 129.9, 128.5, 126.3, 124.2, 119.2, 111.8, 107.5, 105.0, 97.2, 79.4 (d, *J*_C-P_ = 9.1 Hz), 73.9, 72.6, 71.8, 71.6, 71.6, 71.6, 71.6, 71.5, 71.5, 71.4, 71.4, 70.2, 69.5, 66.4 (d, *J*_C-P_ = 5.1 Hz), 65.0 (d, *J*_C-P_ = 5.1 Hz), 60.7, 60.3 (d, *J*_C-P_ = 4.1 Hz), 56.6, 55.6, 52.0, 51.6, 51.0, 46.2, 42.9, 40.0, 39.5, 36.7, 36.6, 34.7, 33.9, 33.3, 33.1, 31.3, 30.9, 30.9, 30.8, 30.8, 30.7, 30.6, 30.5, 30.4, 27.4, 27.2, 27.2, 26.8, 25.5, 23.8, 21.9, 17.7, 14.5, 13.0, 12.8. ^31^P NMR (162 MHz, CD_3_OD) δ 0.79. HRMS calcd. for C_90_H_147_N_6_O_18_P ([M + 2H]^2+^): 815.5255, found: 815.5259.

##### 3- oxo-androst-4-en-17β-carboxylic acid (CRG049)

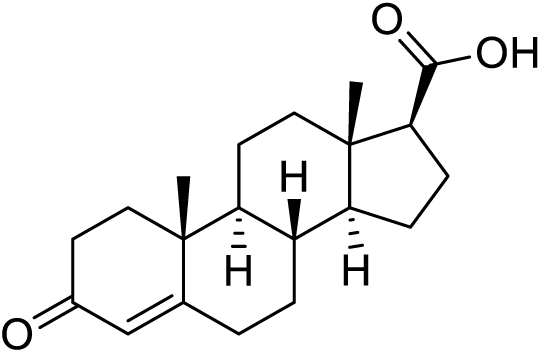

To a stirred solution of NaOH (827 mg, 20.67 mmol) in water (7 mL) was added dropwise Br_2_ (277 µL, 5.41 mmol) at 0 °C and the mixture was stirred at the same temperature for 10 min. The resulting yellowish solution was added dropwise to an ice-cooled solution of progesterone (**6**) (500 mg, 1.59 mmol) in dioxane (15 mL) and water (5 mL) and the mixture was stirred at rt for 1.5 h. A saturated aqueous Na_2_SO_3_ solution (5 mL) was added dropwise until the solution turned colourless and the resulting mixture was refluxed for 15 min. After cooling down to 0 °C, the pH of the solution was adjusted to 2-3 with 1 M HCl. Dioxane was carefully evaporated under reduced pressure and the resulting precipitate was filtered, washed with water, and dried to give the crude product. Purification by flash chromatography (from 0 to 70% EtOAc in hexane) provided **CRG049** (372 mg, 74%) as a pale-yellow solid.

[α]^20^_D_= +155.5 (*c* 1.0, CHCl_3_) [lit.^7^ [α]^25^_D_ = +140.3 (*c* 1.0, CHCl_3_]. ^1^H NMR (400 MHz, CDCl_3_) δ 5.74 (s, 3H), 2.49 – 0.92 (m, 20H), 1.19 (s, 3H), 0.78 (s, 3H). ^13^C NMR (101 MHz, CDCl_3_) δ 199.8, 179.5, 171.3, 124.1, 55.5, 55.1, 53.8, 44.2, 38.8, 38.0, 35.9, 35.8, 34.1, 32.9, 32.0, 24.5, 23.5, 21.0, 17.5, 13.4. HRMS calcd. for C_20_H_29_O_3_ ([M + H]^+^): 317.2117, found: 317.2116.

### 3. Fluorescence quantum yield measurements

Relative quantum yields were measured on a SpectraMax M5 (Molecular Devices) spectrophotometer using a 1 cm path length quartz cuvette. Rhodamine B was obtained from Sigma-Aldrich and was used without further purification. Absolute ethanol (analytical quality) was deoxygenated prior to use. Fluorescence quantum yields were calculated using the following equation:

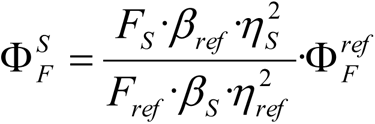

Where 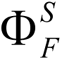 and 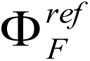 are the fluorescence quantum yield of the sample and that of the standard (Rhodamine B, 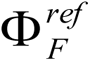= 0.7),^8^ respectively. F_S_ and F_ref_ represent the area of fluorescent emission in units of photons. *η_S_* and *η*_*ref*_ are the refractive indices of the solvent used (ethanol, *η* = 1.361). *β_S_* and *β_ref_* are the correction absorption factors, *β* = 1 −10^−^ *^A^* where A = absorbance.

In order to minimize reabsorption effects, the solutions for quantum yield measurements were prepared such that the optical density was generally about 0.04 at λ_ex_ = 550 nm.

## REFERENCES

1 Gronemeyer, H. Control of transcription activation by steroid hormone receptors. FASEB J 6, 2524–2529, doi:10.1096/fasebj.6.8.1592204 (1992).

2 Beato, M., Herrlich, P. & Schütz, G. Steroid hormone receptors: many actors in search of a plot. Cell 83, 851–857, doi:10.1016/0092-8674(95)90201-5 (1995).

3 Beato, M., Wright, R. H. G. & Dily, F. L. 90 YEARS OF PROGESTERONE: Molecular mechanisms of progesterone receptor action on the breast cancer genome. J Mol Endocrinol 65, T65–t79, doi:10.1530/jme-19-0266 (2020).

4 Pietras, R. J. & Szego, C. M. Specific binding sites for oestrogen at the outer surfaces of isolated endometrial cells. Nature 265, 69–72, doi:10.1038/265069a0 (1977).

5 Migliaccio, A. et al. Tyrosine kinase/p21ras/MAP-kinase pathway activation by estradiol-receptor complex in MCF-7 cells. Embo j 15, 1292–1300 (1996).

6 Migliaccio, A. et al. Activation of the Src/p21ras/Erk pathway by progesterone receptor via cross-talk with estrogen receptor. Embo j 17, 2008–2018, doi:10.1093/emboj/17.7.2008 (1998).

7 Ballaré, C. et al. Two domains of the progesterone receptor interact with the estrogen receptor and are required for progesterone activation of the c-Src/Erk pathway in mammalian cells. Mol Cell Biol 23, 1994–2008, doi:10.1128/mcb.23.6.1994-2008.2003 (2003).

8 Boonyaratanakornkit, V. et al. Progesterone receptor contains a proline-rich motif that directly interacts with SH3 domains and activates c-Src family tyrosine kinases. Mol Cell 8, 269–280, doi:10.1016/s1097-2765(01)00304-5 (2001).

9 Pedram, A. et al. A conserved mechanism for steroid receptor translocation to the plasma membrane. J Biol Chem 282, 22278–22288, doi:10.1074/jbc.M611877200 (2007).

10 Vicent, G. P. et al. Induction of progesterone target genes requires activation of Erk and Msk kinases and phosphorylation of histone H3. Mol Cell 24, 367–381, doi:10.1016/j.molcel.2006.10.011 (2006).

11 Zaurin, R. et al. A set of accessible enhancers enables the initial response of breast cancer cells to physiological progestin concentrations. Nucleic Acids Res 49, 12716–12731, doi:10.1093/nar/gkab1125 (2021).

12 Siiteri, P. K. et al. The serum transport of steroid hormones. Recent Prog Horm Res 38, 457–510, doi:10.1016/b978-0-12-571138-8.50016-0 (1982).

13 Zheng, J., Ali, A. & Ramirez, V. D. Steroids conjugated to bovine serum albumin as tools to demonstrate specific steroid neuronal membrane binding sites. J Psychiatry Neurosci 21, 187–197 (1996).

14 Balleza, D., Garcia-Arribas, A. B., Sot, J., Ruiz-Mirazo, K. & Goñi, F. M. Ether-versus ester-linked phospholipid bilayers containing either linear or branched apolar chains. Biophys J 107, 1364–1374, doi:10.1016/j.bpj.2014.07.036 (2014).

15 Goswami, L. N., Houston, Z. H., Sarma, S. J., Jalisatgi, S. S. & Hawthorne, M. F. Efficient synthesis of diverse heterobifunctionalized clickable oligo(ethylene glycol) linkers: potential applications in bioconjugation and targeted drug delivery. Org Biomol Chem 11, 1116–1126, doi:10.1039/c2ob26968f (2013).

16 Nieves, I. et al. Fluorescent polyene ceramide analogues as membrane probes. Langmuir 31, 2484–2492, doi:10.1021/la505017x (2015).

17 Goretta, S. A. et al. Effects of chemical modification of sphingomyelin ammonium group on formation of liquid-ordered phase. Bioorg Med Chem 20, 4012–4019, doi:10.1016/j.bmc.2012.05.015 (2012).

18 Sandbhor, M. S., Key, J. A., Strelkov, I. S. & Cairo, C. W. A modular synthesis of alkynyl-phosphocholine headgroups for labeling sphingomyelin and phosphatidylcholine. J Org Chem 74, 8669–8674, doi:10.1021/jo901824h (2009).

19 Bhabak, K. P., Proksch, D., Redmer, S. & Arenz, C. Novel fluorescent ceramide derivatives for probing ceramidase substrate specificity. Bioorg Med Chem 20, 6154–6161, doi:10.1016/j.bmc.2012.08.035 (2012).

20 Wichmann, O., Wittbrodt, J. & Schultz, C. A small-molecule FRET probe to monitor phospholipase A2 activity in cells and organisms. Angew Chem Int Ed Engl 45, 508–512, doi:10.1002/anie.200500751 (2006).

21 Pinkert, T., Furkert, D., Korte, T., Herrmann, A. & Arenz, C. Amplification of a FRET Probe by Lipid-Water Partition for the Detection of Acid Sphingomyelinase in Live Cells. Angew Chem Int Ed Engl 56, 2790–2794, doi:10.1002/anie.201611706 (2017).

22 Williams, S. P. & Sigler, P. B. Atomic structure of progesterone complexed with its receptor. Nature 393, 392–396, doi:10.1038/30775 (1998).

23 Di Croce, L. et al. Two-step synergism between the progesterone receptor and the DNA-binding domain of nuclear factor 1 on MMTV minichromosomes. Mol Cell 4, 45–54, doi:10.1016/s1097-2765(00)80186-0 (1999).

24 Grill, H. J., Manz, B., Elger, W. & Pollow, K. 3H-cyproterone acetate: binding characteristics to human uterine progestagen receptors. J Endocrinol Invest 8, 135–141, doi:10.1007/bf03350668 (1985).

25 Gong, W., Chávez, S. & Beato, M. Point mutation in the ligand-binding domain of the progesterone receptor generates a transdominant negative phenotype. Mol Endocrinol 11, 1476–1485, doi:10.1210/mend.11.10.9991 (1997).

26 Mauvais-Jarvis, F., Lange, C. A. & Levin, E. R. Membrane-initiated estrogen, androgen and progesterone receptor signaling in health and disease. Endocr Rev, doi:10.1210/endrev/bnab041 (2021).

27 Quiles, I. et al. Mutational analysis of progesterone receptor functional domains in stable cell lines delineates sets of genes regulated by different mechanisms. Mol Endocrinol 23, 809–826, doi:10.1210/me.2008-0454 (2009).

28 Ballaré, C. et al. Nucleosome-driven transcription factor binding and gene regulation. Mol Cell 49, 67–79, doi:10.1016/j.molcel.2012.10.019 (2013).

29 Gerdes, D., Wehling, M., Leube, B. & Falkenstein, E. Cloning and tissue expression of two putative steroid membrane receptors. Biol Chem 379, 907–911, doi:10.1515/bchm.1998.379.7.907 (1998).

30 Alonso, A., Sáez, R., Villena, A. & Goñi, F. M. Increase in size of sonicated phospholipid vesicles in the presence of detergents. J Membr Biol 67, 55–62, doi:10.1007/bf01868647 (1982).

31 Böttcher, C. J. F., Pries, C. & van Gent, C. M. A rapid and sensitive colorimetric microdetermination of free and bound choline. Recueil des Travaux Chimiques des Pays-Bas 80, 1169–1178, doi:https://doi.org/10.1002/recl.19610801102 (1961).

32 Angelova, M. I. & Dimitrov, D. S. Liposome electroformation. Faraday Discussions of the Chemical Society 81, 303–311, doi:10.1039/DC9868100303 (1986).

33 Montes, L. R. et al. Electroformation of giant unilamellar vesicles from native membranes and organic lipid mixtures for the study of lipid domains under physiological ionic-strength conditions. Methods Mol Biol 606, 105–114, doi:10.1007/978-1-60761-447-0_9 (2010).

34 Wang, X., Spandidos, A., Wang, H. & Seed, B. PrimerBank: a PCR primer database for quantitative gene expression analysis, 2012 update. Nucleic Acids Research 40, D1144–D1149, doi:10.1093/nar/gkr1013 (2011).

35 Ferrari, R. et al. TFIIIC Binding to Alu Elements Controls Gene Expression via Chromatin Looping and Histone Acetylation. Molecular Cell 77, 475–487.e411, doi:https://doi.org/10.1016/j.molcel.2019.10.020 (2020).

36 Bolger, A. M., Lohse, M. & Usadel, B. Trimmomatic: a flexible trimmer for Illumina sequence data. Bioinformatics 30, 2114–2120, doi:10.1093/bioinformatics/btu170 (2014).

37 Lander, E. S. et al. Initial sequencing and analysis of the human genome. Nature 409, 860–921, doi:10.1038/35057062 (2001).

38 Dobin, A. et al. STAR: ultrafast universal RNA-seq aligner. Bioinformatics 29, 15–21, doi:10.1093/bioinformatics/bts635 (2012).

39 Bligh, E. G. & Dyer, W. J. A rapid method of total lipid extraction and purification. Can J Biochem Physiol 37, 911–917, doi:10.1139/o59-099 (1959).

40 Strutt, H. & Paro, R. Mapping DNA target sites of chromatin proteins in vivo by formaldehyde crosslinking. Methods Mol Biol 119, 455–467, doi:10.1385/1-59259-681-9:455 (1999).

## References

(1) S. J. Briggs, M.; Bruce, I.; N. Miller, J.; J. Moody, C.; C. Simmonds, A.; Swann, E. Synthesis of functionalised fluorescent dyes and their coupling to amines and amino acids. J. Chem. Soc. Perkin Trans. 1 1997, No. 7, 1051–1058.

(2) Goretta, S. A.; Kinoshita, M.; Mori, S.; Tsuchikawa, H.; Matsumori, N.; Murata, M. Effects of chemical modification of sphingomyelin ammonium group on formation of liquid-ordered phase. Bioorg. Med. Chem. 2012, 20 (13), 4012–4019.

(3) Sandbhor, M. S.; Key, J. A.; Strelkov, I. S.; Cairo, C. W. A Modular Synthesis of Alkynyl-Phosphocholine Headgroups for Labeling Sphingomyelin and Phosphatidylcholine. J. Org. Chem. 2009, 74 (22), 8669–8674.

(4) Lao, K.; Sun, J.; Wang, C.; Lyu, W.; Zhou, B.; Zhao, R.; Xu, Q.; You, Q.; Xiang, H. Design, synthesis and biological evaluation of novel androst-3,5-diene-3-carboxylic acid derivatives as inhibitors of 5α-reductase type 1 and 2. Steroids 2017, 124, 29–34.

(5) White, J. D.; Lincoln, C. M.; Yang, J.; Martin, W. H. C.; Chan, D. B. Total Synthesis of Solandelactones A, B, E, and F Exploiting a Tandem Petasis−Claisen Lactonization Strategy. J. Org. Chem. 2008, 73 (11), 4139–4150.

(6) Nakamura, N.; Miyazaki, H.; Ohkawa, N.; Oshima, T.; Koike, H. An efficient synthesis of platelet-activating factor (PAF) via 1-O-alkyl-2-O-(3-isoxazolyl)-SN-glycero-3-phospho-choline, a new PAF agonist utilization of the 3-isoxazolyloxy group as a protected hydroxyl. Tetrahedron Lett. 1990, 31 (5), 699–702.

(7) Rey, J.; O’Riordan, T. J. C.; Hu, H.; Snyder, J. P.; White, A. J. P.; Barrett, A. G. M. Design and Diastereoselective Synthesis of C-2,C-20-Diaryl Steroidal Derivatives. Eur. J. Org. Chem. 2012, 2012 (20), 3781–3794.

(8) Arbeloa, F. L.; Ojeda, P. R.; Arbeloa, I. L. Flourescence self-quenching of the molecular forms of Rhodamine B in aqueous and ethanolic solutions. J. Lumin. 1989, 44 (1), 105– 112.

